# Modeling Dragons: Using linked mechanistic physiological and microclimate models to explore environmental, physiological, and morphological constraints on the early evolution of dinosaurs

**DOI:** 10.1101/790980

**Authors:** David M. Lovelace, Scott A. Hartman, Paul D. Mathewson, Benjamin J. Linzmeier, Warren P. Porter

**Author notes:** Earth and Planetary Sciences Department, Northwestern University, Evanston, Illinois, United States of America (Benjamin J. Linzmeier).

## Abstract

We employed the widely-tested biophysiological modeling software, Niche Mapper™ to investigate the metabolic function of Late Triassic dinosaurs *Plateosaurus* and *Coelophysis* during global greenhouse conditions. We tested them under a variety of assumptions about resting metabolic rate, evaluated within six microclimate models that bound paleoenvironmental conditions at 12° N paleolatitude, as determined by sedimentological and isotopic proxies for climate within the Chinle Formation of the southwestern United States. Sensitivity testing of metabolic variables and simulated “metabolic chamber” analyses support elevated “ratite-like” metabolic rates and intermediate “monotreme-like” core temperature ranges in these species of early saurischian dinosaur. Our results suggest small theropods may have needed partial to full epidermal insulation in temperate environments, while fully grown prosauropods would have likely been heat stressed in open, hot environments and should have been restricted to cooler microclimates such as dense forests (under any vegitative cover) or those seen at higher latitudes and elevations. This is in agreement with the Late Triassic fossil record and may have contributed to the latitudinal gap in the Triassic prosauropod record.

## Introduction

Paleontologists have long inferred the biology of extinct organisms from morphological correlates and paleoenvironmental context. Hypotheses derived from morphology rely on extant phylogenetic bracketing and modern analogs for support [1, 2]; lack of inferential specificity may come from trimmed phylogenetic trees or increased distance from extant relatives [3]. Spatiotemporally derived hypotheses suffer from confounding factors related to bias in the stratigraphic record [4–6]. These biases can be tempered by physics-based constraints to understand a broad range of paleobiological phenomena.

All animals, living and extinct are constrained by physics. Gravity exerts control on the maximum size attained by terrestrial clades, from spiders to sauropods [7], and biological thermodynamics constrains the rate heat is produced and how it is transferred to and from the environment [8]. Robust biophysiological models, such as Niche Mapper™ are built on the fundamental properties of heat and mass flux into and out of individuals [9]. Morphology (e.g. posture, insulation, color, and body part dimensions), behavior (e.g. seeking shade, sunning, fur or feather erection, varying activity level and location), and physiology (e.g. metabolic rate, peripheral blood flow, respiratory and cutaneous water loss) can accelerate or retard heat exchange with the surrounding environment that sets the temporal and spatial constraints (boundary conditions) for animal function.

For decades ecologists have been modeling the physics of heat transfer to understand ecological and biogeographic constraints of modern organisms [10–25]. Biophysiological models have only been sparsely applied to deep time investigations [e.g. 26, 27]. For paleoecologists, the paleobiogeographic distribution of an extinct organism is harder to test with respect to organismal physiology. For instance, it has been noted that there is an absence of large (>∼1000 kg) prosauropod dinosaurs in the well-studied tropical to subtropical latitudes during the Late Triassic (e.g., the Chinle Formation of southwestern U.S.), while smaller (<∼100 kg) theropod dinosaurs and their closest relatives are quite common [28]. In contrast, both large prosauropod and small theropod remains are well represented in temperate latitudes. It has been hypothesized that high temperatures and environmental fluctuations including increased severity and frequency of wildfires, aridity, and precipitation patterns precluded large prosauropods from inhabiting these tropical environments [28], this hypothesis is hard to test explicitly without the use of biophysical modeling.

We applied Niche Mapper to test a range of physiological possibilities for two Late Triassic dinosaurs, the ∼20 kg theropod *Coelophysis bauri* (Cope 1887) and the ∼1000 kg prosauropod *Plateosaurus engelhardti* (von Meyer 1837) to quantitatively test the thermal constraint hypothesis. We performed sensitivity analyses on the models to test the relative contributions of climate, body mass, shape, diet, insulation, and the efficiencies of respiratory, digestive, and muscle systems, as well as feasible daily core temperature range and resting metabolic rate on energy requirements. The integration of biophysiological and microclimate models offers a potentially powerful means of testing feasible physiologies and behaviors of extinct organisms derived for a given paleoenvironmental.

## Materials and methods

We employed the mechanistic modeling program Niche Mapper, developed at UW-Madison [29]. Niche Mapper is compartmentalized into generic microclimate and animal submodels that each contain momentum, heat, and mass transfer equations. The microclimate model has 51 variables relating to seasonality, insolation, shade, wind, air temperature, humidity, cloud, and soil properties. The biophysical model is composed of 270 morphological, physiological, and behavioral parameters previously described in detail and tested over a wide range of animal taxa including reptiles, amphibians, birds, mammals and insects [e.g., 12,13,15,16,23,30–39]. The user is able to control how many days (1 to 365) and how those days are distributed throughout the year, and how many of the variables (such as air temperature, feather density, or body mass) vary for each modeled day (see S1 Appendix).

Fundamentally, Niche Mapper calculates hourly energy and mass expenditures that can predict survivorship based on reasonable bounds of thermal stress and resource availability. Paleoenvironmental bounds for the microclimate model are derived from environmental proxies preserved within the rock record and data from analogous modern environments, when applicable. Measurements of skeletal dimensions parameterize a simple geometric volume that approximates the shape of the animal. The range of metabolic rates known from modern tetrapods bounds modeled rates. These boundaries allowed us to explore a reasonable parameter space although the model could easily be extended to explore unique circumstances and test novel hypotheses.

As an additional means of testing our Triassic microclimate models and as a point of comparison for dinosaurs modeled with a squamate-like metabolism, we created a model for the largest extant terrestrial ecotherm, *Varanus komodoensis* (Komodo dragon), using the same methodologies implemented in our dinosaur models along with experimental and observational data for *V. komodoensis* [40–42]. We compared calculated vs. observed agreement between activity patterns, food requirements, body size and body temperature (S2 Appendix).

### The microclimate model

The microclimate model calculates hourly air temperatures, wind speeds, and relative humidity profiles, solar and thermal long wavelength infrared radiation, ground surface and subsurface temperatures available to an animal [11,29,43,44]. Local environments at mean animal height are used for heat balance calculations (Fig. 1).

**Figure 1.**
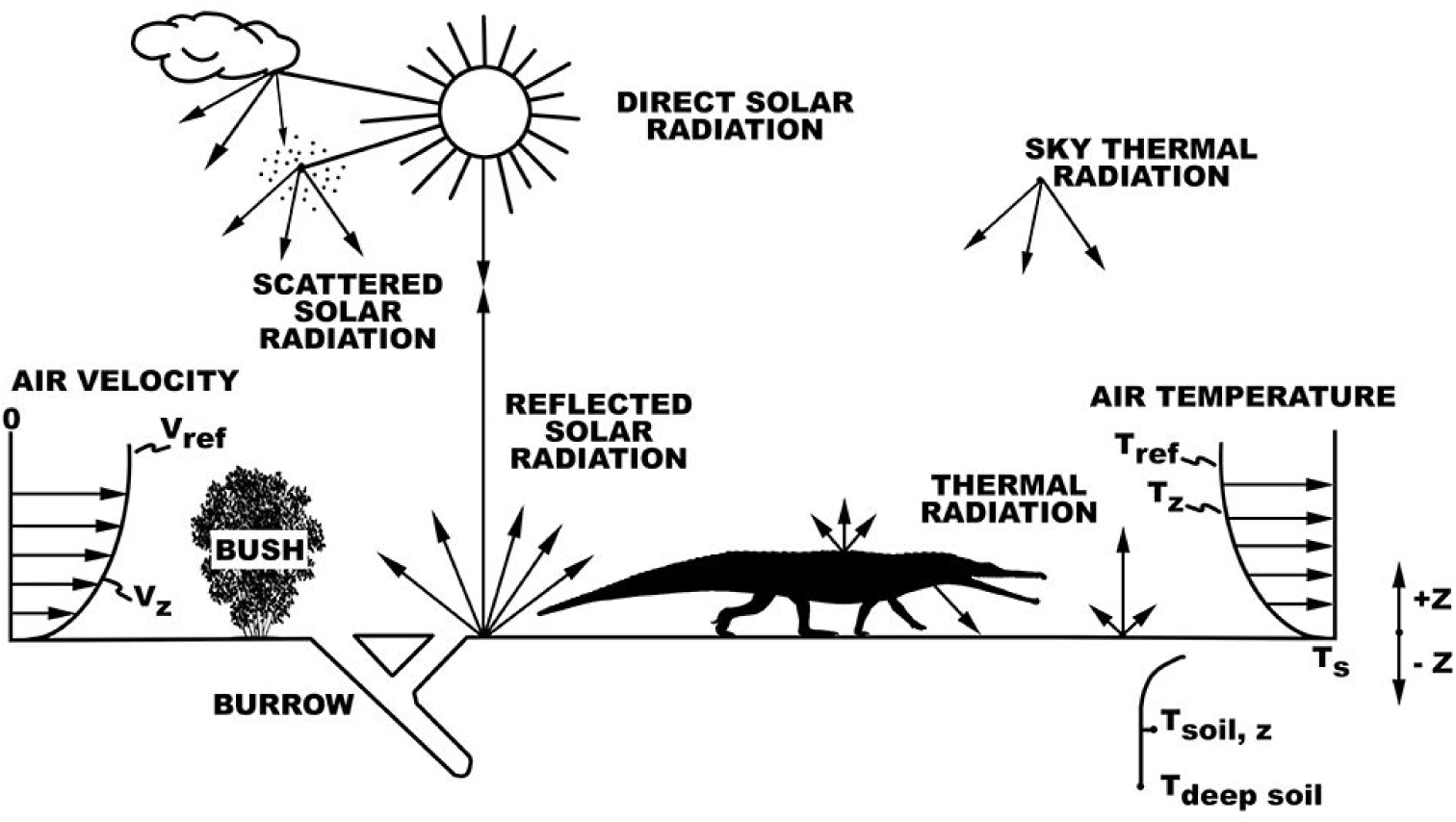
Organism-environment heat balance interactions. The reptile’s heat and mass balances that determine body temperature are determined by where it chooses to be each hour to remain within its preferred body temperature range. Niche Mapper allows it to find a location each hour where it can remain active, or not, if necessary, to optimize its body temperature and/or water balance.

The microclimate model fits a sinusoidal curve to user-specified maximum and minimum daily air temperatures, wind speeds, cloud cover, and relative humidity to estimate hourly values. We set minimum air temperature, minimum wind speed, maximum relative humidity and cloud cover to occur at sunrise. Maximum air temperature, maximum wind speed, and minimum relative humidity and cloud cover are set to occur one hour after solar noon [45]. Clear sky solar radiation is calculated based on date, hour of day, and geographic location adjusted for cloud cover and overhead vegetation [43, 46]. Paleosolar calculations for insolation were computed from Laskar et al. [47] using the *palinsol* program in R [48]. Because of uncertainties in deep time insolation we used modern values as a first step. However, we recognize there is variability in insolation not only in deep time, but also on shorter timescales due to precession, obliquity, and eccentricity. Cloud cover also reduces solar radiation intensity at ground level and provides thermal cover by trapping longwave radiation that would otherwise escape to the sky, increasing the sky’s radiant temperature [29,44,45,49].

Long wavelength thermal radiation from clear sky and clouds were computed using empirical air temperature correlations from the literature [50, 51]. Substrate thermal radiation was computed hourly from the numerical solution of a one dimensional finite difference transient heat balance equation for the ground. Hourly outputs from the microclimate model specify above and belowground local microclimates in the sunniest and shadiest sites specified by the user.

### Microclimate Model Parameterization in Deep Time

In order to test the hypothesis that large dinosaurs such as *Plateosaurus* were restricted from tropical latitudes primarily due to thermal constraints we chose to use the well sampled Late Triassic strata of the Chinle Formation of western North America as a model for our Late Triassic paleoenvironment. Although these strata are currently located at 35 degrees north, the Chinle was originally deposited between 5 and 10 degrees north paleolatitude [52]. We used published local and regional paleoclimate data derived from the Chinle Formation; these data include sedimentary proxies for paleotemperature and precipitation [52–54], global climate models [55], and global geochemical compilations [56, 57]. We used these data to constrain our microclimate model to best represent a tropical Late Triassic environment and considered this the ‘hot’ microclimate model. This is not the most extreme temperatures proposed for the Late Triassic Chinle Formation [58], making our modeled environment a conservative estimate for testing whether high temperatures excluded large dinosaurs from the region. Our cold microclimate is an conservative approximation of paleoclimates in upper Triassic strata in central and northern Europe (∼35-45 °N; [55,59,60]) where *Plateosaurus* is a relatively common constituent in fossil assemblages. In order to aid comparison and avoid interactions of variables, annual distribution of microclimate data was maintained across the hot and cold microclimates and only the temperature values were adjusted to model a cooler temperate microclimate as a first-order approximation of higher latitudes. A moderate microclimate was also modeled to represent areas intermediate between hot and cold microclimates. Air temperature are modeled at the average height of the organisms being examined for each hour (Fig. 2). Paleolatitude is modeled as 12 °N, with an elevation of 150 m [54].

**Figure 2.**
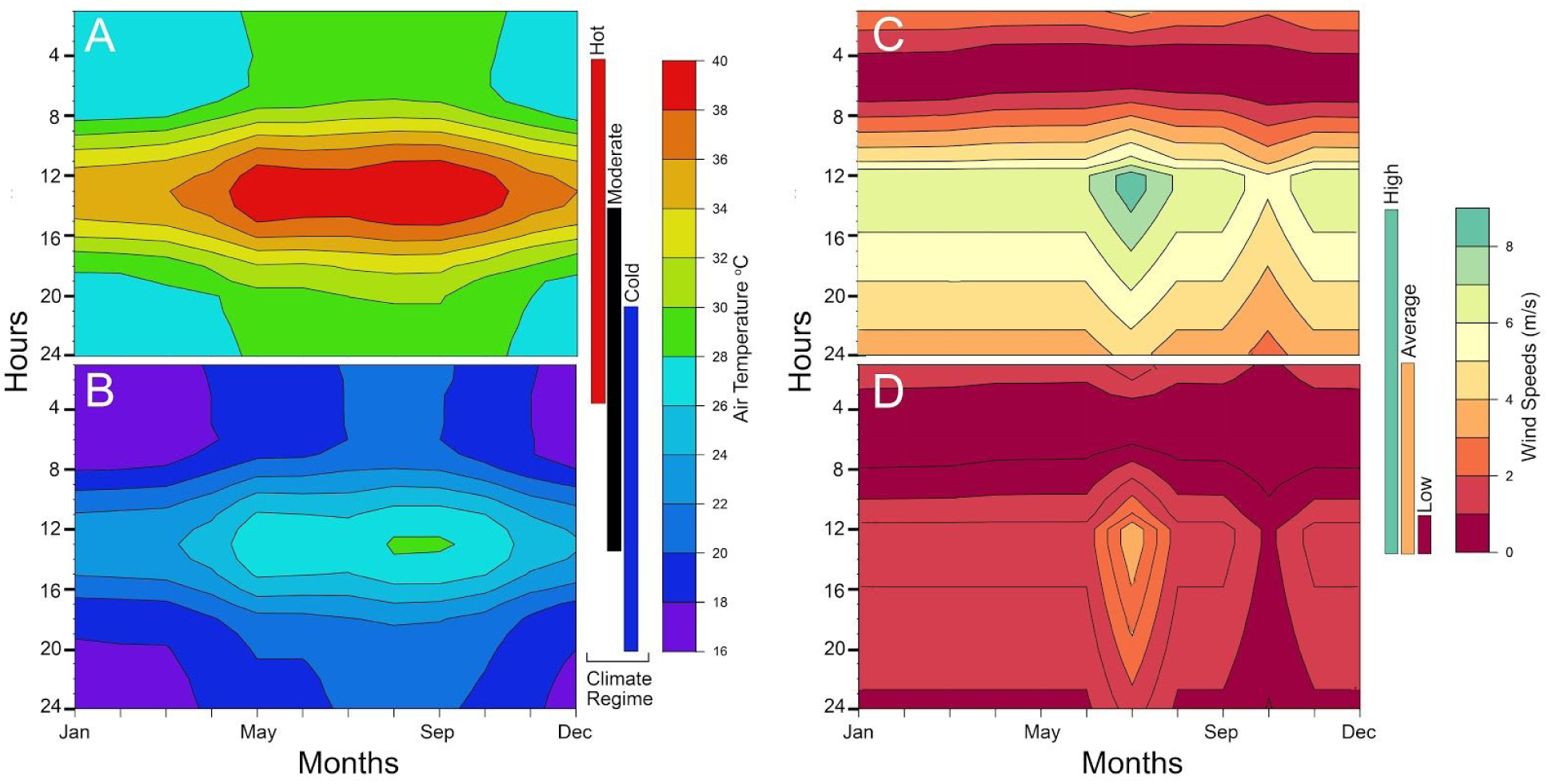
Heatmaps of microclimate air temperature and wind speed at average animal height for each modeled hour. We use a turbulent velocity and temperature profile where the most significant changes occur within the first 15 cm from the ground. The microclimates are the same for both *Coelophysis* (shown) and *Plateosaurus*. (A) hot and B) cold climate regimes, C) high and D) low wind speeds.

Because the resolution of paleoclimate proxies are time-averaged it is difficult to determine an annual pattern for the microclimate model. As such, historic climate data from modern analogues provide a means to establish realistic annual patterns otherwise indiscernible in the rock record. We selected regions similar to paleogeographic reconstructions of the Chinle formation with respect to elevation, temperature and precipitation regimes, latitude, and general position on the continent; two locations in western Africa (Tamale, Ghana and Timbuktu, Mali) act as modern analogues for this study. Numeric values for the microclimate model were determined by multi-proxy data in the geological record whenever possible (see Table 1).

**Table 1.**
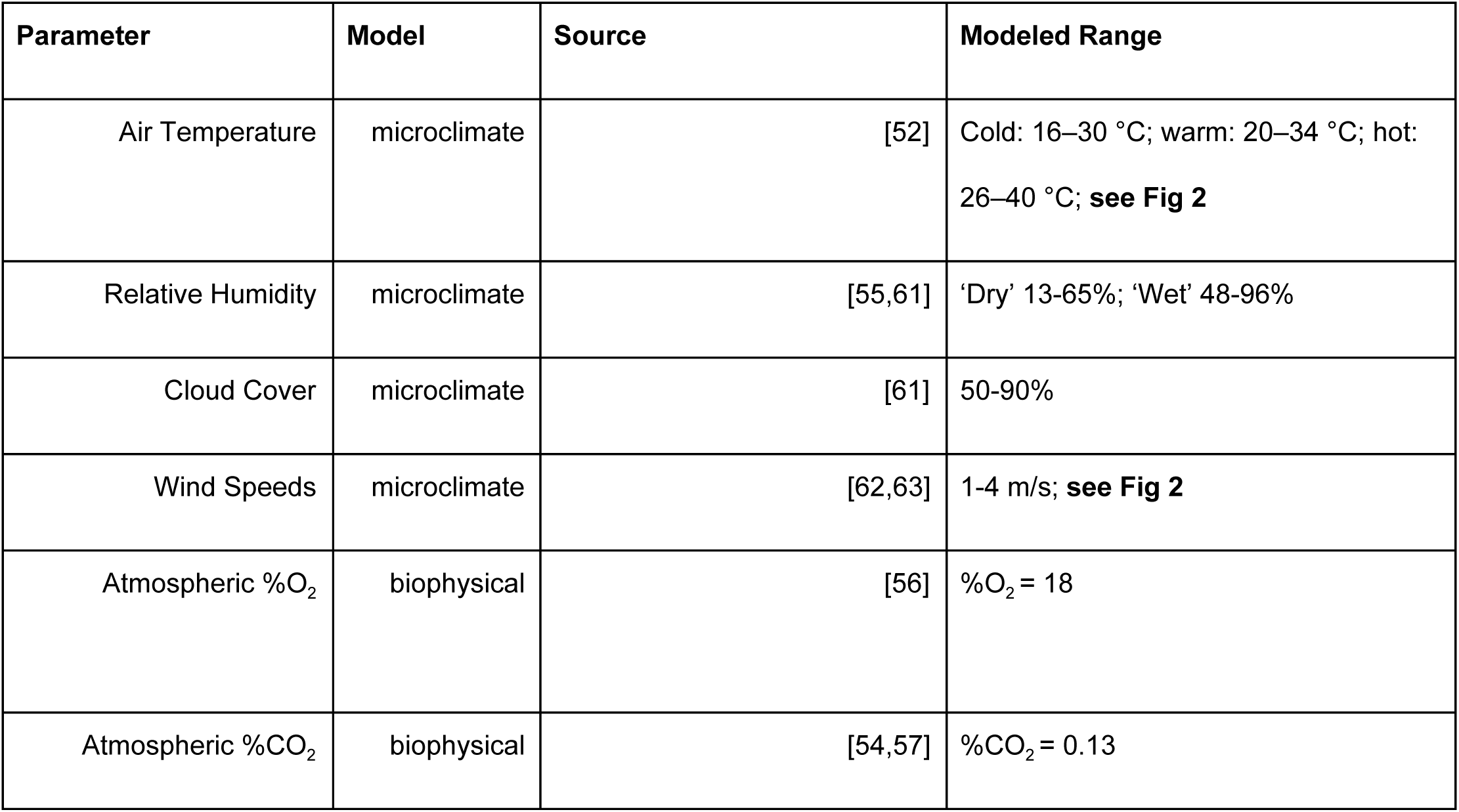
Microclimate parameters inferred from geologic proxies, GCM’s, and modern analogues.

### The biophysical model

Gross organismal morphology (head, neck, torso, legs, tail) are modeled as simple geometric shapes (e.g., cylinders, spheres, truncated cones, or ellipsoids; S3 Appendix). These geometries have known or measurable heat transfer properties and temperature profile equations that simplify solving the heat balance equation when there is distributed internal heat generation [e.g., 36,64,66]. Each body part can be modeled with up to three concentric layers: 1) a solid central geometry of tissue uniformly generating heat; 2) if present, a surrounding layer of insulating subcutaneous fat is modeled as a hollow heat conducting geometry; 3) a surrounding layer of insulating fur or feathers, modeled as a hollow porous medium (see below). Net metabolic heat produced by the central flesh layer must be transferred through the fat layer to the skin surface, where is it either dissipated via cutaneous evaporation, convection and infrared thermal radiation (naked), or transferred through the fur/feather layer if present, then lost by convection and infrared thermal radiation to the environment. Heat is transferred through the fur/feathers by parallel thermal radiation and conduction through the air between the insulation elements and through the fur or feathers [36, 66] and is lost to the environment via thermal radiation and convection. If the animal is lying down and has ventral insulation, it is compressed by an amount defined by the user and heat is conducted to or from the substrate at the insulation surface. Heat from solar radiation can also be absorbed through the skin (naked) or fur/feather layer (insulated), contributing to the heat load that must be dissipated (Fig. 3).

**Figure 3.**
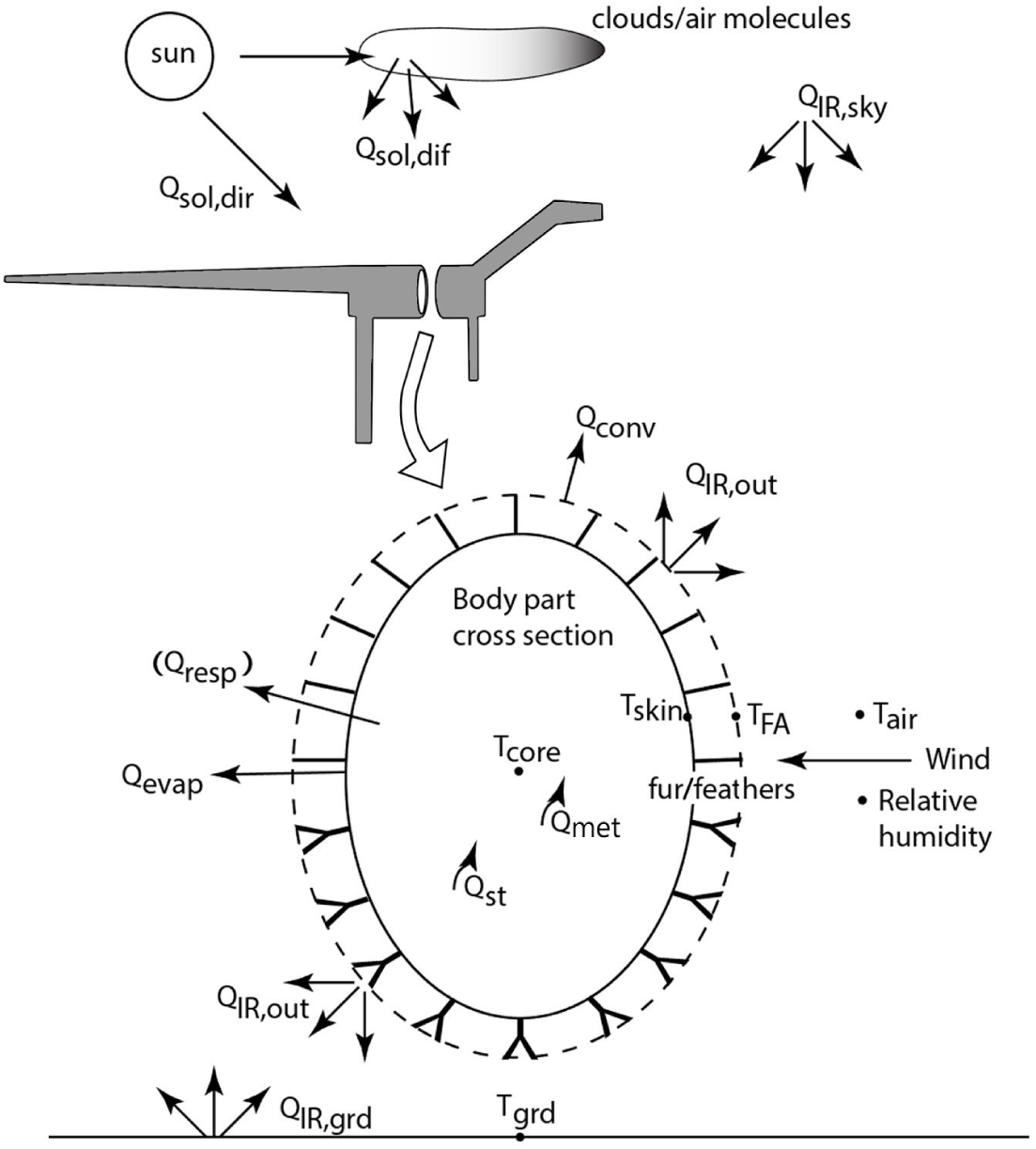
Heat transfer pathways between modeled organism and environment. Cross-section of a body segment (e.g., elliptical cylindrical torso of distributed heat generating flesh surrounded by an optional layer of fat (not shown), then skin surrounded by porous insulation whose properties may be the same or different dorsally vs. ventrally). The flesh is generating metabolic heat throughout the body (*Qmet)* and exchanging (*Qresp*, *Qevap*, *QIRnet*, *Qconv*, *Qsol*) heat with its environment as modeled by Niche Mapper (adapted from Porter et al. [23]). The transient model also includes a heat storage term, *Qst*, for the flesh. A full list of symbols and abbreviations can be found in the text.

Provided with the local environmental conditions from the microclimate model and biophysical properties of the organism, the animal model calculates radiative (*Q_rad_*), convective (*Q_conv_*), solar (*Q_sol_*), and evaporative (respiratory: *Q_resp_* and cutaneous: *Q_evap_*) heat fluxes between the animal and its microenvironment to solve a heat balance equation (1) for a metabolic rate, *Q_met_*, that satisfies (1) and is consistent with the status of its core and skin temperatures and environmental conditions:

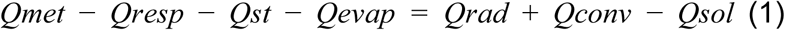

If the animal has a fur or feather layer, an additional parameter (*Qfur*) must be added to account for heat flow through the insulating fur or feather layer:

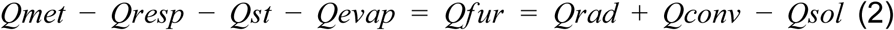

where *Q_fur_* represents the heat flux through the fur or feather layer via parallel conductive and radiative processes. For more detailed explanations of heat flux through porous media such as fur or feathers and solving for steady state conditions see Porter et al. [36], Porter and Kearney [67] and Mathewson and Porter [17].

For each hour of every model day the heat balance equation is solved for individual body parts and summed to provide the total metabolic rate (W) for the entire animal that will allow it to maintain a target core temperature in that hour’s range of environmental conditions. Users can specify a basal metabolic rate multiplier to simulate activity in the heat balance calculations, as well as muscle efficiency, which is the proportion of that additional activity expenditure contributes to the animal’s heat balance (i.e., 0% means that the mechanical work (activity) is 100% efficient with no excess heat produced; 99% means that 99% of the metabolic effort is lost as heat and needs to be considered in the heat balance). We assumed a mammal-like 20% muscle efficiency for activity with 80% of the chemical energy for activity going to heat. Although a ∼35% muscle efficiency may be more reasonable for archosaurs, it is less well documented [68]; we performed a sensitivity analysis to test the effect of differing muscle efficiencies (see below).

If the total animal metabolic rate deviates from user-specified variation in the target metabolic rate (i.e., expected basal metabolic rate x activity multiplier) for the hour being modeled, physiological options, followed by behavioral options, are engaged to prevent the animal from being too hot or too cold by decreasing or increasing metabolic expenditure on heat production respectively.

User selected physiological options are engaged in the following order when individually enabled: 1) incrementally erect fur or feathers to increase insulation; 2) incrementally increase or decrease flesh thermal conductivity, simulating vasodilation or vasoconstriction of peripheral blood vessels; 3) incrementally increase or decrease core temperature, simulating temporary, bounded positive or negative heat storage; 4) incrementally increase the amount of surface area that is wet to increase evaporative heat loss, simulating sweating (if allowed); and 5) incrementally decrease oxygen extraction efficiency to increase respiratory heat loss, simulating panting.

If physiological changes are not sufficient to thermoregulate, behavioral thermoregulation options are engaged and the animal can seek shade, swim, wade, climb, or enter a burrow to achieve cooler environmental conditions. The user defines which behaviors are possible for the modeled organism; for instance, it is unlikely that an 850 kilogram prosauropod is burrowing or climbing trees to behaviorally thermoregulate, so these options would not be utilized. If the animal is too cold (i.e. the requisite metabolism is greater than the resting metabolic rate x the activity multiplier), the user can allow the animal to enter a burrow or seek vegitative shelter or get out of the wind. These options also reduce radiant heat loss to the sky by providing overhead structures (e.g. forest canopy or burrow ceiling) with warmer radiant temperatures than the open sky. Users can also allow model animals to make postural changes such as curling up to minimize surface area for heat exchange with the environment if the animal is too cold.

The heat balance is re-solved after each incremental thermoregulatory change until either 1) the metabolic rate that balances the heat budget equation is within the percent error of the target metabolic rate or 2) thermoregulatory options are exhausted. In the case of the latter, the metabolic rate that balances the heat budget equation that is closest to the target rate is used for that hour. Hourly metabolic rates and water losses are integrated over the day to calculate daily metabolic rate and water loss, which can then be used to calculate food and water requirements. The day’s water and energy requirements are then used to compute the respiratory and digestive system inputs and outputs using molar balances as described below.

The heat and mass balance of an organism are connected by metabolic rate, a ‘biological fire’ that requires fuel and oxygen. The daily metabolic rate that releases heat (*Qmet*) sets the daily mass balance requirements for the respiratory and digestive systems (Fig. 4). Diet composition (proteins, carbohydrates, lipids, percent water) specify how much mass must be absorbed (**m_abs_**) from the gut to meet metabolic demands. Digestive efficiency divided into the mass absorbed determines the mass of food that must be ingested per day (***m_in_***) to meet energy requirements. Mass excreted (***m_out_***) is the difference between mass in and mass absorbed.

**Figure 4.**
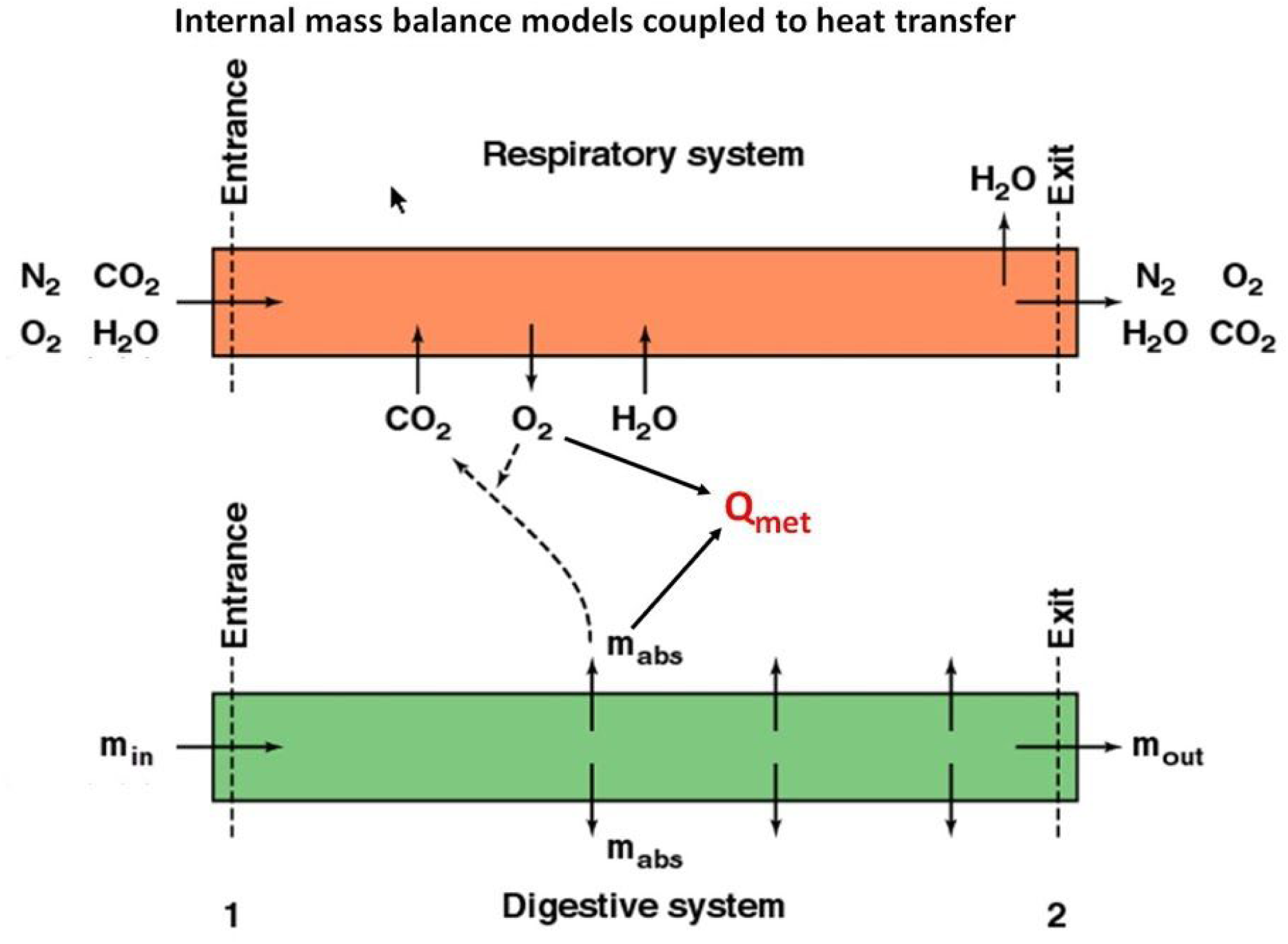
Internal mass balance models coupled to heat transfer. System diagram for the respiratory and digestive system driven by the metabolic rate, *Qmet*.

The respiratory system functions in an analogous manner. Diet utilization and activity rates determine the amount of oxygen needed and carbon dioxide produced. Oxygen required divided by the respiratory extraction coefficient specifies the mass of air that must enter the respiratory system, given the atmospheric composition of oxygen per unit volume. Humidity of the incoming air is increased to saturated air at lung (body) temperature, so respiratory water loss can also be computed. Recovery of water vapor during exhalation through cooler nostrils is also calculated.

### Biophysical Model Parameterization in Deep Time

Key *morphological* and *behavioral* model inputs are summarized in Tables 2-3.

**Table 2.** Morphological parameters of porous insulation modeled and skin or insulation surface solar reflectivity. Insulation element density estimated assuming that 1) feathers evolved by selection for a follicle that would grow an emergent tubular appendage [74] early ‘feathers’ were similar to stage 1 or 2 in the developmental pattern of modern feathers, i.e. cylindrical in shape. 3) that the minimum hair density that substantially reduces heat loss (reduces metabolic heat generation that sets body temperature) is approximately 800-1000 ‘hairs’ /cm^2^ [37 Fig 25].

| Variable |  |  | No<br>Insulation | Top-only<br>Insulation | Full<br>Insulation | Reference and<br>comment |
| --- | --- | --- | --- | --- | --- | --- |
| Insulation | Length<br>(mm) | Dorsal* | 0 | 30 | 30 | [Based on 70-73] |
|  |  | Ventral* | 0 | 0 | 30 |  |
|  | Depth<br>(mm) | Dorsal* | 0 | 10 | 10 |  |
|  |  | Ventral* | 0 | 0 | 10 |  |
|  | Density<br>(elements/cm<sup>2</sup>) |  | 0 | 2000 | 2000 | [37 Fig 25,74] |
|  | Insulation/skin<br>reflectivity (%) |  | 15 | 15 | 15 | Porter, marine<br>iguana, unpub. data |
|  | Pilo/ptiloerection |  | No | Yes** | Yes** |  |

\* head, neck, torso, and tail
\*\*only when behaviors are enabled

**Table 3.**
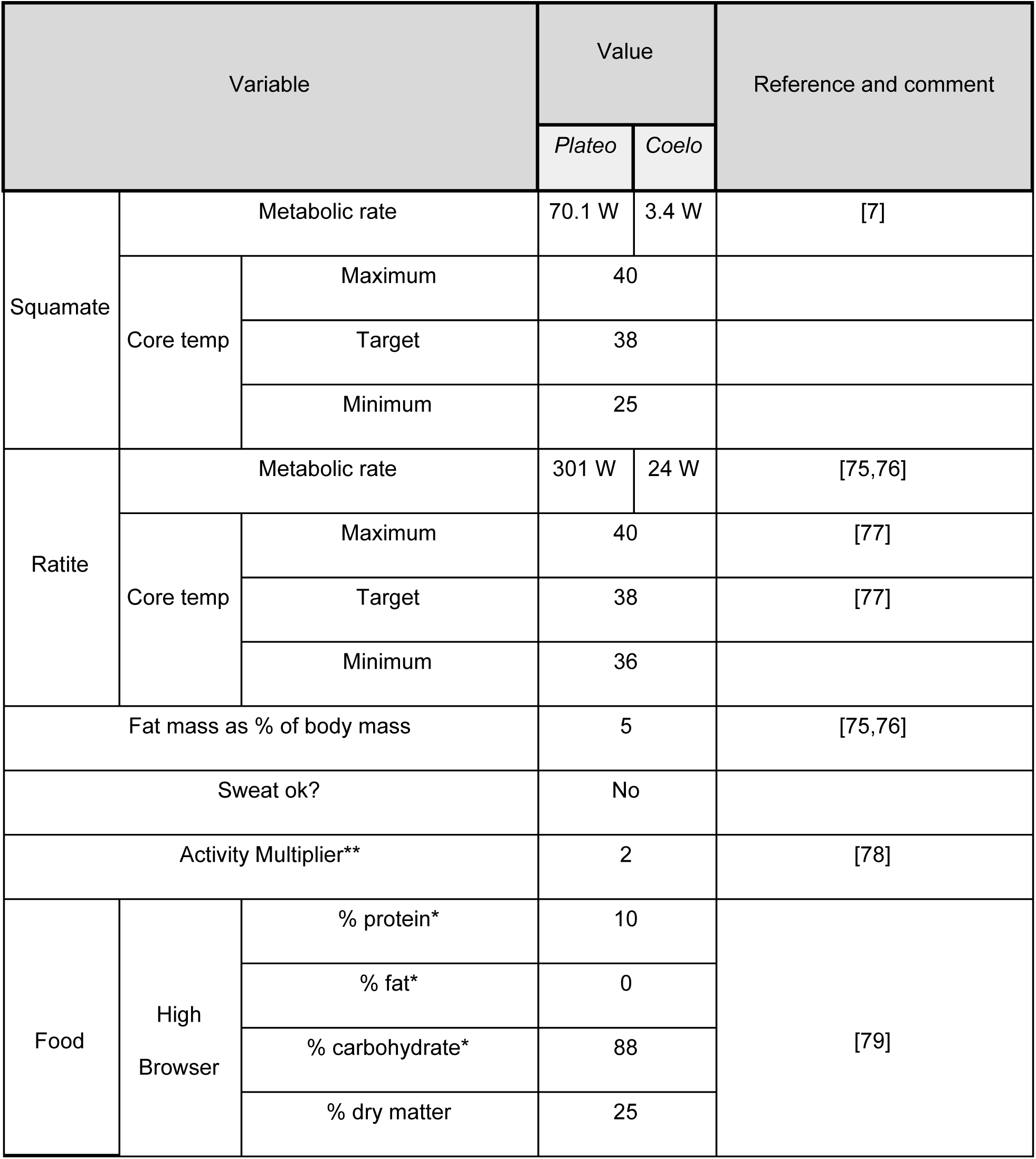
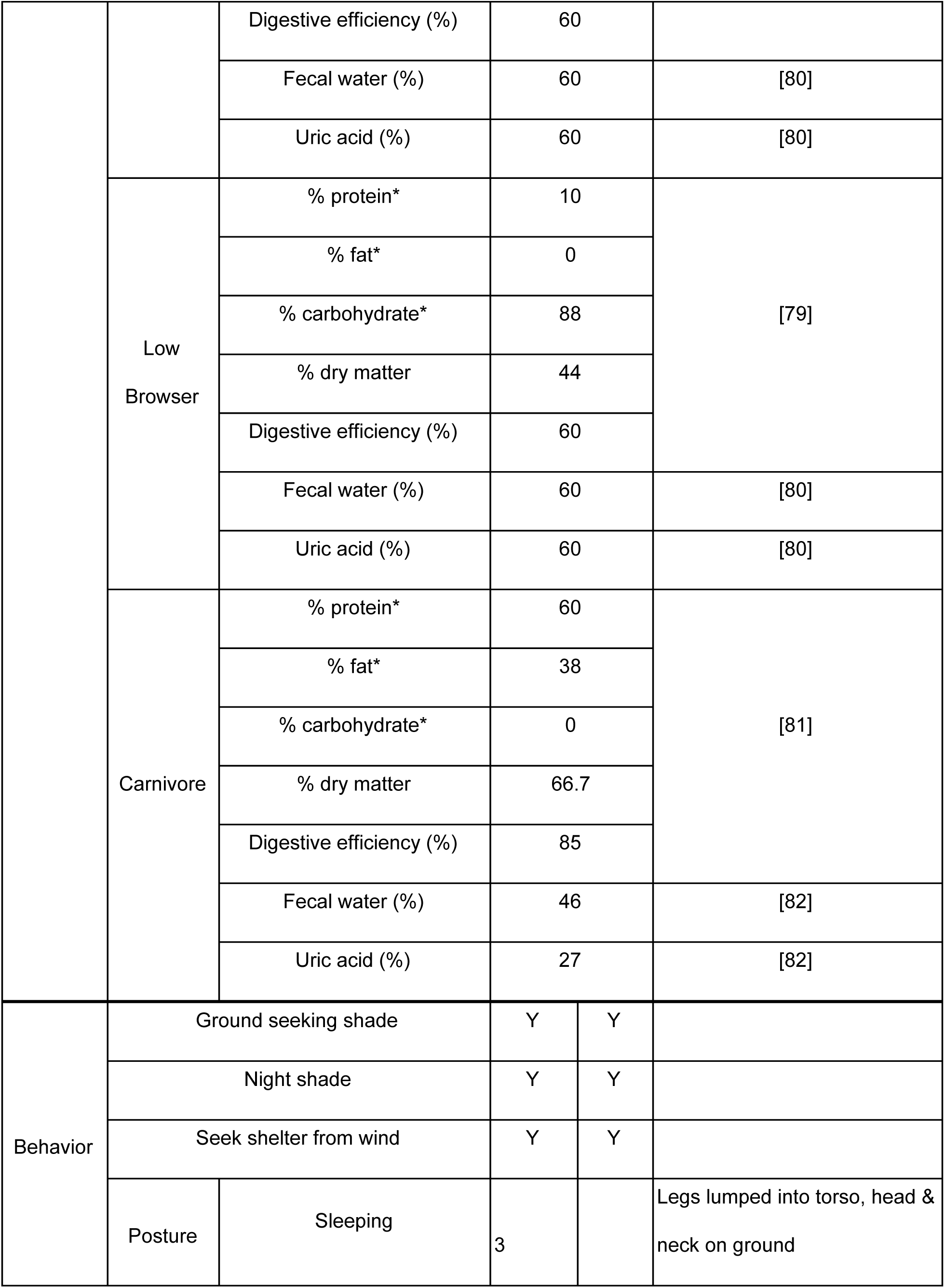
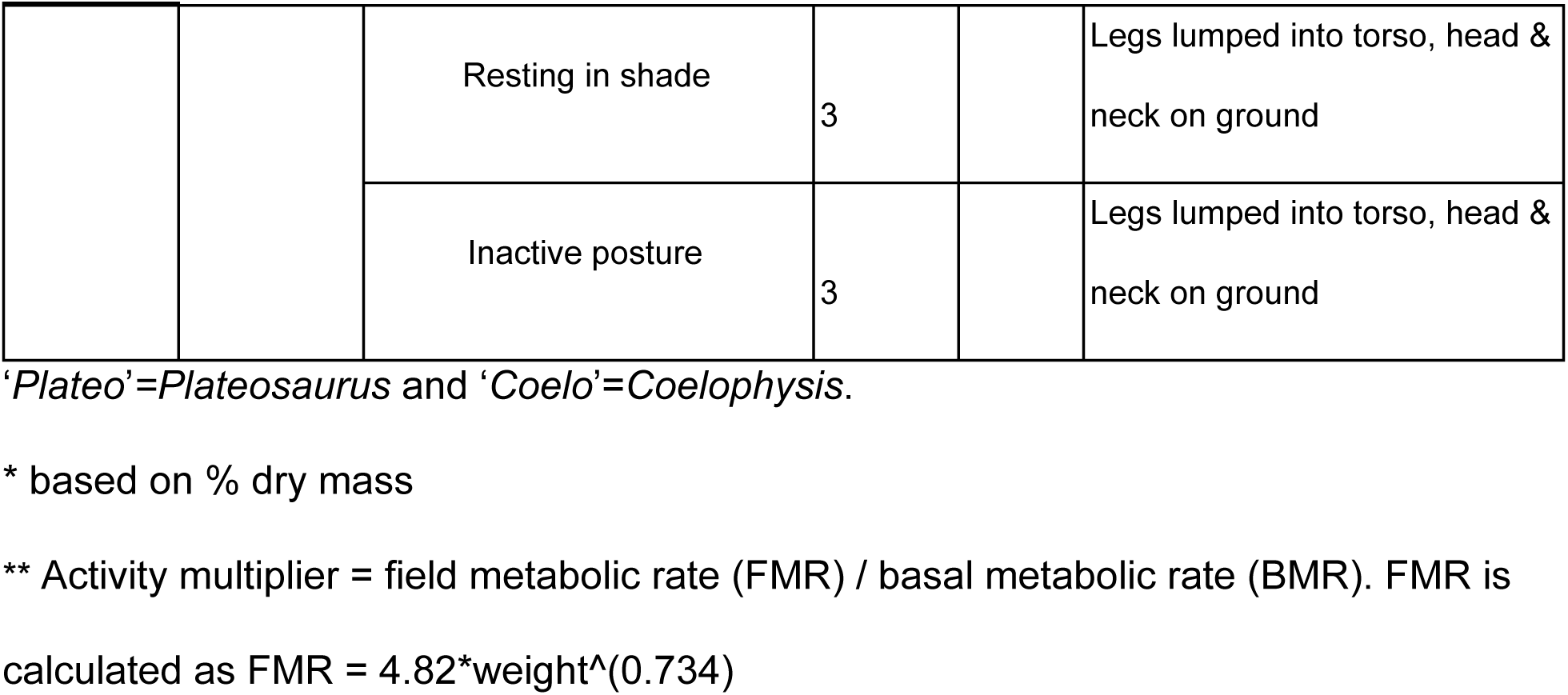
Parameters for metabolic rates, diet, and behavior.

For *behavioral* thermoregulation, we allowed animals to hide from wind (if too cool), seek shade during the day (if too hot), seek shade at night (e.g., simulating vegitative cover slowing the rate of radiative heat loss seen in open sky conditions), be active in the shade (day and night), postural changes to minimize surface area if cold and inactive, or maximize surface area if hot. For *physiological* thermoregulation we allowed for panting (too hot), increased and decreased flesh conductivity (to simulate vasoconstriction/vasodilation when cold/hot), if dermal insulation (fur/feathers) is present they can piloerect/ptiloerect, or changes in regulated body temperature within the user specified maximum and minimum body temperature range. Sweating was not enabled due to phylogenetic constraints. Sensitivity analyses were performed to test which behavior, or interaction of multiple behaviors, had the strongest effect. Since legs and tail consist of mostly muscle, bone and tendon, we allowed the temperature in limb and tail segments to reach 50% of the difference between the torso-segment junction and ambient air or ground temperatures [c.f.,12,69].

### Determining rates of metabolism

To determine a range of metabolic rates for extinct taxa within our modeled microclimate, we simulated a spectrum of different metabolic rates. We evaluated 5 different resting metabolic rates (RMRs) ranging from a typical ectothermic squamate to endothermic eutherian metabolisms. An RMR from the lower avian range (ratite) and two lower mammal (monotreme and eutherian) RMRs were calculated from empirical regressions utilizing phylogenetic and ecological constraints [e.g.,75, 76], while the squamate, and an additional eutherian RMR were calculated using empirical models derived from oxygen consumption or CO_2_ production measurements with an assumed respiratory quotient [7, 83]. These regressions implicitly include the presence or absence of epidermal insulation of extant species as well as their size and shape, but the data provide a range of values for estimating the span of modern mass-specific metabolic rates(S1 Table).

Using the calculated masses (S3 Appendix) and the empirical equations above we generated five different mass-specific resting metabolic rates for *Coelophysis* and *Plateosaurus*, labeled: squamate, monotreme, marsupial, ratite, and eutherian. We analysed the thermoneutral range for each RMR in a virtual metabolic chamber simulation within Niche Mapper. We chose to conduct the remainder of the model simulations with low (squamate), moderate (monotreme), and high (ratite) RMRs representing a possible range of metabolic rates based on phylogenetic position. It is unlikely that basal dinosaurs had metabolic rates elevated above extant ratites, or below extant squamates.

### Diets

In the Niche Mapper model, the primary outputs influenced by diet are daily values for discretionary water (kg/day) and food requirement (kg/day). The diet (required caloric intake) is calculated based on daily energy expenditure, user-supplied values for percent protein, fat, carbohydrates, and dry mass of the food, and the animal’s assimilation efficiency (Table 3). The amount of water initially available to the organism is calculated from the diet-assigned dry mass and the amount of food consumed (e.g., total food mass - dry mass = total available free water from food). Metabolic water production is computed from diet composition and metabolic rate [84 p. 489, 695]. Daily water loss is the total of cutaneous water evaporation (if sweating were allowed; we did not allow sweating) and water lost through respiration and excretion (in mammal-based models this includes water loss through feces and urine, in non-mammal models water loss through urine is ignored). If daily water budget is negative, the organism must drink water to make up the volume; thus, discretionary water reflects the debt or credit of the total water budget after all modeled physiological needs are met. A user defined digestive efficiency controls the amount of incoming calories (food mass) that is required by the model organism to meet its metabolic needs (Table 3).

*Coelophysis* has long been considered a predatory theropod [85, 86] and *Plateosaurus* is usually described as a herbivorous prosauropod [6]. To assess the impact of varying inferences of diet on food and water requirements *Coelophysis* and *Plateosaurus* were both modeled as carnivores and as high and low browsing herbivores. The diet in the high browsing scenario is comprised of primarily high % dry matter (e.g., 40-50% dry mass) such as conifers (8.3 MJ/kg dry mass), ginkgos (8.6 MJ/kg dry mass), and cycads (6.1 MJ/kg dry mass) [79, 87]. Low browsing diets were primarily composed of ferns (7.7 MJ/kg dry mass) and *Equisetum* (11.6 MJ/kg dry mass) which contain much higher water content (e.g., 25-30% dry mass) [79, 87]. A positive discretionary water budget would indicate the animal is getting most of its water from its food source as well as metabolic water and may not require regular access to drinking water extending its potential geographic range.

### Energy requirements

We developed an R script to interface Niche Mapper with a modifiable database containing climate (Table 1) and physiological variables (Tables 2-3) for each of the experimental simulations, which were assessed for 6 unique climates [hot, moderate, cold] x [arid, humid]. For each simulation Niche Mapper calculated hourly interactions between the organisms and their environment over a 24 hour period at mid-month for a calendar year (12 total model days).

In all of our modeled simulations, each species is given the potential ability to be active every hour of the day (24 hours). The amount of metabolic heat production (W) needed to maintain the target core temperature throughout the day is determined by multiplying the resting metabolic rate (Table 4) by an activity multiplier (2.0 in our study; [78]). The resultant is the daily target metabolic rate (MJ/day):

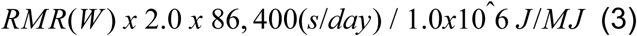

**Table 4.** Annual predicted energy budget (MJ/year) for both dinosaur species.

| Daily/Annual target ME (MJ) |  |  |  |  |  |  |  |  |  |  |
| --- | --- | --- | --- | --- | --- | --- | --- | --- | --- | --- |
|  |  |  |  | Daily target metabolic output (MJ) |  |  |  | Annual target metabolic output (MJ) |  |  |
|  | Mass (kg) | Ecto RMR* | Mono RMR** | Ratite RMR** | Ectotherm | Monotreme | Ratite | Ecto YMR | Mono YMR | Ratite YMR |
| Coelophysis | 15 | 2.6 | 9.7 | 17.5 | 0.45 | 1.67 | 3.02 | 163.08 | 610.29 | 1102.58 |
|  | 21 | 3.4 | 12.3 | 22.2 | 0.59 | 2.13 | 3.83 | 214.36 | 778.64 | 1397.75 |
|  | 30 | 4.5 | 16.0 | 28.5 | 0.79 | 2.76 | 4.92 | 286.62 | 1008.05 | 1797.36 |
|  | 40 | 5.7 | 19.7 | 34.9 | 0.99 | 3.40 | 6.03 | 362.43 | 1241.48 | 2201.49 |
|  | 50 | 6.9 | 23.1 | 40.9 | 1.19 | 4.00 | 7.06 | 434.87 | 1459.16 | 2576.55 |
|  | 60 | 8.0 | 26.4 | 46.5 | 1.38 | 4.56 | 8.03 | 504.72 | 1665.06 | 2929.96 |
| Plateosaurus | 600 | 52.7 | 139.8 | 235.5 | 9.11 | 24.16 | 40.70 | 3325.34 | 8819.21 | 14854.63 |
|  | 850 | 70.1 | 179.9 | 301.1 | 12.12 | 31.09 | 52.03 | 4422.46 | 11348.73 | 18989.15 |
|  | 1150 | 89.9 | 224.0 | 372.6 | 15.53 | 38.70 | 64.38 | 5668.06 | 14125.16 | 23499.42 |
|  | 1600 | 117.8 | 284.4 | 470.3 | 20.36 | 49.15 | 81.26 | 7430.36 | 17940.35 | 29659.90 |
|  | 2000 | 141.5 | 334.3 | 550.4 | 24.44 | 57.77 | 95.10 | 8921.95 | 21085.98 | 34712.93 |
|  | 3000 | 197.2 | 448.4 | 732.5 | 34.08 | 77.48 | 126.57 | 12440.32 | 28280.29 | 46199.32 |
|  |  | * McMahan model |  |  |  |  |  |  |  |  |
|  |  | ** McNab model |  |  |  |  |  |  |  |  |

Thus any decrease in activity hours represent periods of the modeled day when the animal is heat stressed and must decrease activity to lower its metabolic heat production. If the animal is cold stressed or within its active thermoneutral zone it will maintain 24 hour activity. However, if it is cold stressed the animals metabolic heat production and by extension, food consumption, must increase accordingly.

We define the *active* thermoneutral zone as the zone where an activity multiplier > 1 is expanding the temperature range in which the animals internal heat production balances the heat loss to the external environment (steady state condition). In contrast, a *resting* thermoneutral zone is the temperature range when the activity multiplier is 1.

Four physiological conditions were used to test the viability of each modeled organism under six microclimate conditions mentioned above (see Table 1). Low (squamate) and high (ratite) resting metabolic rates were calculated based on equations from McNab [75, 76] and McMahon [7], each of which were analyzed with a broad squamate-like core temperature range (CTR), which ranged from 26-40°C, moderate monotreme-like CTR (32-40°C), and a narrow ratite-like CTR (36-40°C). All CTRs were assigned a target core temperature of 38°C.

### Metabolic Chamber

Metabolic chamber simulations in Niche Mapper were used to evaluate the specific impact of different physiological inferences affecting the temperatures in which model animals were predicted to be cold or heat stressed. In the metabolic chamber simulations, all temperatures (ground, sky, and air) are set equal to one another, no solar input is allowed, a constant, negligible wind speed of 0.1 m/s is used, along with a constant 5% relative humidity. Animals are modeled in a standing posture with no activity multiplier. Heat balance calculations were then performed along a range of temperatures (0-51°C) that exceeded the minimum and maximum air temperatures experienced where each organism, with each modeled metabolic rate, could maintain thermoneutrality [65] in order to identify lower and upper critical temperature boundaries.

### Sensitivity Analyses

Niche Mapper is an effective tool for modeling extant organisms where direct measurements can be applied. Modeling organisms in deep-time is faced with a number of challenges where direct measurements are not possible. Variables such as air temperature, core temperature range, resting metabolic rate, and insulatory structures have a significant effect on the modeled organisms annual metabolic energy and are tested for and visualized in each modeled experiment. Additional parameters, such as muscle efficiency, digestion, and respiration, as well as mass estimates, skin reflectivity, and insolation factors related to latitude are independently tested for model sensitivity. In order to determine how sensitive the model was to these additional parameters, a bounded range that includes our hypothesized values were modeled for each parameter.

For instance, there is uncertainty as to the color of skin or insulatory structures in most extinct animals. For our purposes, it is known that a lighter color absorbs less solar radiation, a darker color absorbs more and this variable could be easily selected for in a given environment [88–90]. To test the effect of color on total metabolic energy requirements we modeled the skin of *Plateosaurus* and the uninsulated *Coelophysis* as well as the proto-feathers for the top-only and fully insulated *Coelophysis* with 5 states ranging from high to low reflectivity (light to dark color, respectively) in both our cold and hot microclimate.

Similarly we tested main and interactive effects of parameters related to climate (i.e.,temperature, wind, relative humidity, cloud cover). To assess the relative effect of these parameters we used the metric of total annual energy for each species and determined how annual metabolic expenditure would change relative to the target value for all variables individually and combined. This approach allowed us to evaluate the main effects of each of these variables as well as possible interactions between them. To determine main effects and interactions of the variables on annual metabolic expenditures for *Plateosaurus* and *Coelophysis* we used a 2^4^ (climate) full factorial design and Yates’ algorithm for analysis of effects [91]; minimum and maximum data are outlined in Table 1.

## Results

### Sensitivity Analyses

The strength of our modeled results, in part, relies on understanding how the model responds to ranges of values for variables that are not directly measurable in deep-time. We conducted the following analyses to quantify the advantage or disadvantage our chosen values would impact on the model results: skin/insulation reflectivity, muscle efficiency, respiratory efficiency, digestive efficiency, the effect of latitude, and mass estimates; a summary of these results follows - further details and figures are provided in supplemental data (S4 Appendix).

Skin and insulation color (reflectivity) was analyzed from 10 – 60% (15% was our chosen model value). It was observed that the disparity in ME between high and low reflectivity values increased with increasing cold stress. For example, the more cold-stressed the model was (i.e., >4-5 x RMR), the greater advantage low reflectivity values (darker color) had. However, there was a negligible effect of reflectivity for models whose annual ME was near target (e.g., between 2x and 3x RMR). Similarly, muscle efficiency ranged from 20 – 50% efficient (20% was our chosen model value) and the disparity between the lowest and highest values for a given model increased as cold-stress increased (see S4 Appendix); there was a negligible effect for models whose RMR was between 2 and 3x resting. An analysis of respiratory efficiency ranged from 10 – 30% (we chose a min-max value of 15-20%) and there was virtually no change in annual ME regardless of which parameter was used.

Digestive efficiency was analyzed with a range of 70 – 85% efficiency (we chose 85%) for the carnivorous diet, and 30-70% (we chose 60%) for the herbivorous diets (see S4 Appendix for discussion). All diet parameters are independent of metabolic calculations and thus did not affect annual ME. Varying the efficiency of digestion provides us with a range of annual wet food requirements. These values can be used to compare with reasonable rates of browsing (or prey acquisition/consumption) relative to modern analogs. For instance, even at the lower extreme, a 30% digestive efficiency for *Plateosaurus* would require ∼8000 kg wet-food per year; this is ∼22 kg of wet-food per day, which is on par with similarly sized extant browsing mammals such as the black rhinoceros [92, 93].

Because we are using our cold microclimate as a proxy for higher latitudes we also tested our models at 45°N [e.g., 55,59,60]. The primary effect of increasing latitude was a result of increased daylight hours midyear and decreased daylight hours during the winter months. This is most apparent in the increased hours/day that core temperature was maintained, midyear, and decreased during the winter months relative to those observed at 12°N. The model is more sensitive to microclimate temperatures than variance in insolation due to increased latitude between12 and 45°N.

The parameters outlined above had relatively small effects on metabolic needs of the modeled organisms that were able to maintain an annual ME between 2 and 3x RMR. However, we realize these effects can be cumulative and are more significant at the boundaries of a modeled organisms’ temperature tolerance where small changes can be the difference between survival or death. There is also the potential effect of interaction between parameters. To test the main and interactive effects of four primary climate parameters (temperature, humidity, wind speed, and cloud cover) a 2^4^ factorial design and Yates’ algorithm [91] was analyzed (S4 Figure 6). Temperature had 2-10 times the effect of wind, and both humidity and cloud cover were insignificant. Variables that have the greatest impact, such as temperature, CTR, RMR, and insulation are presented below with a range of inputs for each experiment.

### Mass estimates

Mass estimates can vary widely for a given taxon depending on methodology [94–99]. Niche Mapper uses a user-supplied mass and distributes that mass, with assigned densities, among each body segment (head, neck, torso, front legs, hind legs, and tail). We tested our modeled organisms with 6 mass estimates from a low to an extreme high mass in both the hot and cold microclimate. In addition to increasing the estimated mass we also accounted for the necessary increase in RMR with mass (see Table 4).

Linear measurements of a *Coelophysis* specimen (AMNH 7224) yield a skeletal length of 2.61 m. Mass estimates for a x meter long specimen are…xxx. With an estimated average density of 0.97 kg/l our *Coelophysis* model has a mass of 21 kg. This is consistent with previously assigned masses [94, pg. 260] of 15-20 kg for a ‘gracile’ and ‘robust’ skeleton, respectively. It is unlikely that the mass exceeds 30 kg for the skeletal dimensions used for this analysis of mass estimates. We tested the effect of increased mass via increasing the diameter of the model’s body segments (i.e., making it thicker) with mass assignments of 15, 21, 30, 40, 50, and 60 kg (see S4 Figure 7). Results for monthly metabolic energy with a high (ratite-like) RMR and CTR (relative to target) for the uninsulated, top-only insulated, and fully insulated *Coelophysis* model demonstrate the effect of mass and insulation in hot and cold microclimates (Fig. 5). Results for varied RMR and CTR for the 21 kg *Coelophysis* follow.

**Figure 5.**
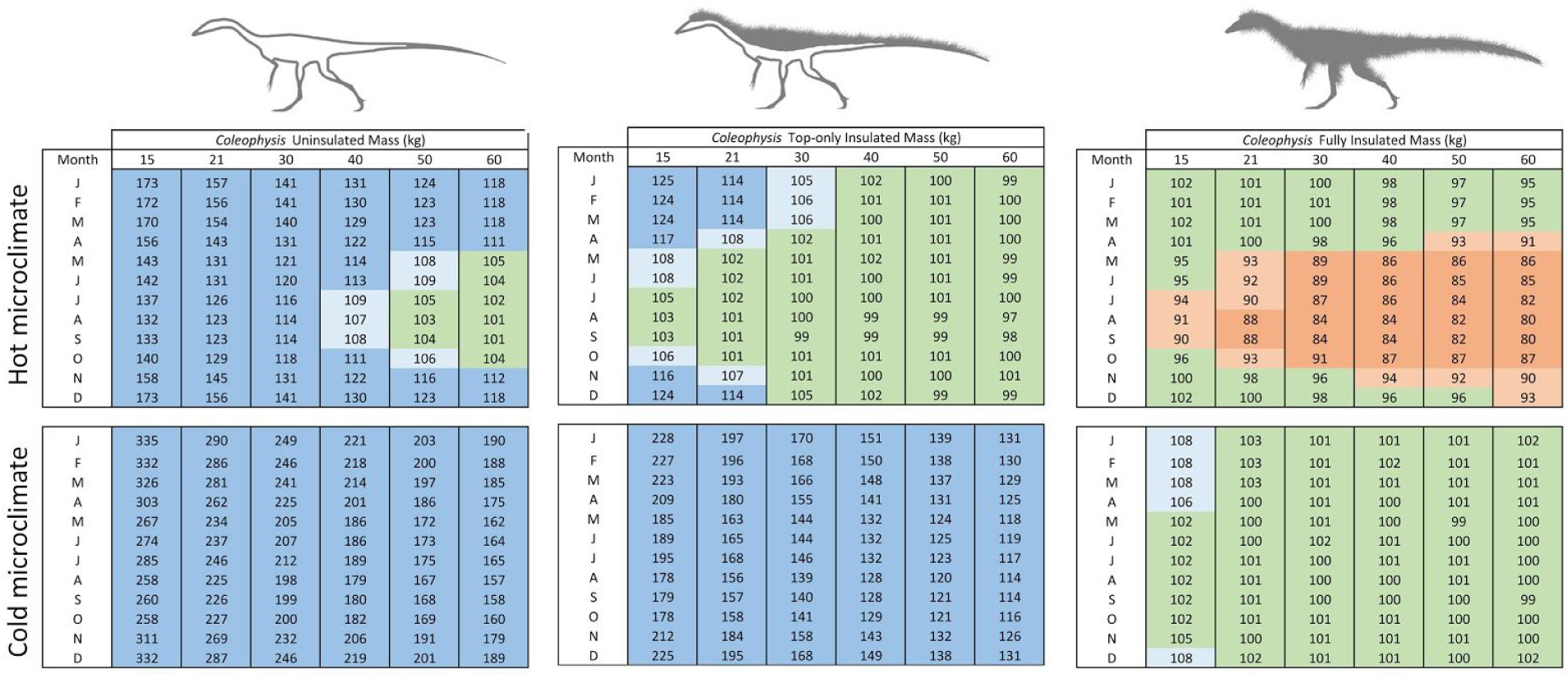
Effect of mass estimate (*Coelophysis*) on annual energy. The matrix reflects the effect of size and insulation for *Coelophysis* in a hot and cold microclimate. Dark blue = >10% above target ME; light blue = 5-10% above target ME; green = +/- 5% of target ME; light orange = 5-10% below target ME; and dark orange = >10% below target ME.

The uninsulated model resulted in extreme cold-stress for the lower three sizes in the hot microclimate, and all 6 mass estimates in the cold microclimate. When increased to top-only insulation the severity of cold-stress decreased, but the model was still excessively cold-stressed in the cold microclimate. The hot microclimate resulted in more months where the model was able to maintain its target metabolic energy, including summer months of the lower three size estimates and all months for the largest three mass estimates. The fully insulated model shows heat-stress during the summer months in the hot microclimate, gradually increasing in severity and temporal extent with increased mass. Under the cold microclimate, the model met target metabolic energy for all but the 15 kg *Coelophysis*, which exhibited minor cold-stress within the winter months (between 5 and 10% above target).

Our linear dimensions for *Plateosaurus* were taken from GPIT/RE/7288, a six meter long skeleton with a femoral length of 635 mm. With an estimated density of 0.97 kg/l the model has a mass of 850 kg. This is in line with mass estimates for moderate sized *Plateosaurus* specimens (i.e., 5.67 m skeleton [595 mm femoral length] with a mass between 660-782 kg using a 0.89-1.05 kg/l density respectively; [98]). Other *Plateosaurus* mass estimates include a 6.5 m long skeleton (1073 kg using polynomial method [99]) and a 920 kg mass determined by stylopodial circumference using a 685 mm femoral length [100].

To test the effect of different mass estimates, we chose to take the same skeletal dimensions and increase or decrease the diameter of body segments (assigned densities remained constant). We ran experiments assuming a total mass of 600, 850, 1150, 1600, 2000, and 3000 kg (Fig. 6; see also S4 Figure 8). The first three states (600-1150 kg) more likely capture a realistic mass estimate range for the skeleton and are representative of mass estimates in the literature for this specimen [97–100]. The last three states (1600-3000 kg) were used to observe how an extreme overestimate of mass would affect the model results. Results for monthly metabolic energy with a high (ratite-like) RMR and CTR (relative to target) for the *Plateosaurus* model demonstrate the effect mass has for hot and cold microclimates (Fig. 6). Results for varied RMR and CTR for the 850 kg *Plateosaurus* follow.

**Figure 6.**
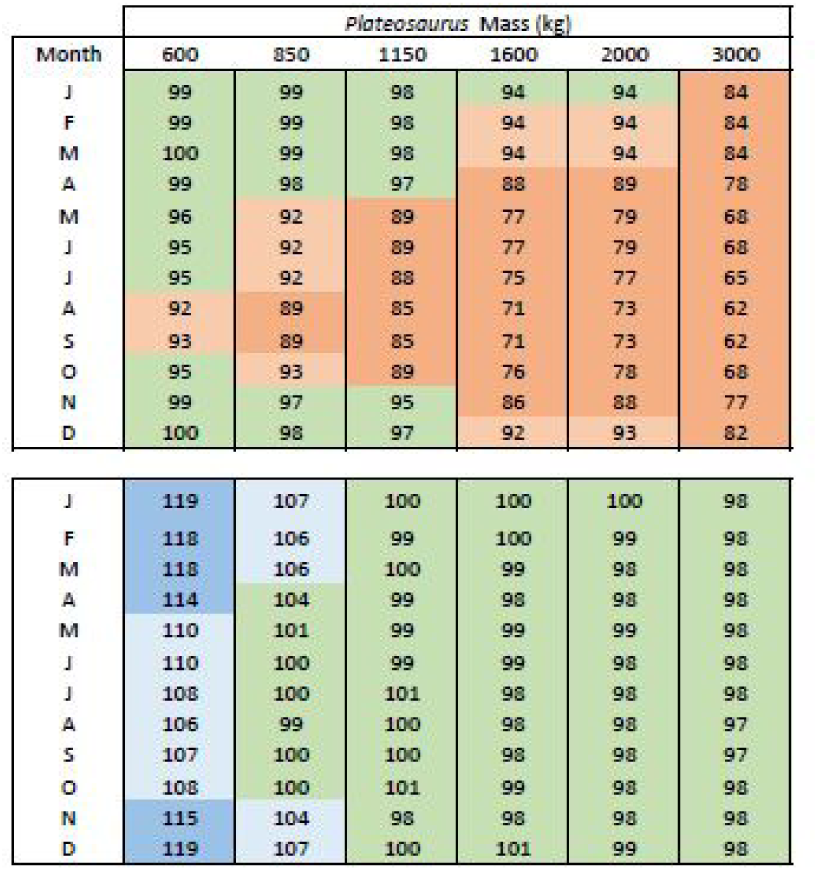
Effect of mass estimate (*Plateosaurus*) on annual energy. The matrix reflects the effect of size and insulation for *Plateosaurus* in a hot and cold microclimate. Dark blue = >10% above target ME; light blue = 5-10% above target ME; green = +/- 5% of target ME; light orange = 5-10% below target ME; and dark orange = >10% below target ME.

The 600 kg model was mildly heat stressed in the hot microclimate during peak summer temperatures, however it was excessively cold (ME ∼15-20% above target) in the cold microclimate. Under the hot microclimate, the 850 kg model we identified as most likely for our 6 m skeleton met its target ME during the cooler winter and spring/fall seasons, but experienced significant heat stress (ME ∼10% below target) during peak summer temperatures. As mass increased, this trend was amplified in the hot microclimate producing excessive heat stressed models. The 850 kg model experienced modest cold stress in the winter months, while the four largest mass estimates met expected target values within the cold microclimate.

### Diet

It has been suggested [87] that *Equisetum* would have been a favored food source (from a nutritional point of view) due to its higher degradability (e.g., 11.6 MJ/kg dry matter) relative to various conifers or *Ginko* (8.3, 8.6 MJ/kg dry mass, respectively). However, given the high water content of *Equisetum* (∼70% [79]) relative to conifers (∼44% [79]) the degradable energy per kilogram of wet mass (what the animal actually consumes) is nearly identical: 3.5 MJ/kg wet mass (*Equisetum*) vs 3.6 MJ/kg wet mass (various conifers) [79, 87]. The various ferns reported by Hummel and others [87] have nearly 75% water content and yield 7.7 MJ/kg dry mass (or 2.1 MJ/kg wet mass). Thus, an animal eating dominantly ferns will need to consume 60% more vegetative mass than an organism whose diet is primarily composed of conifers or horsetails.

The diet component of the model, although extremely useful for certain questions, is calculated based on the resulting metabolic energy outputs. Factors such as digestive efficiency, food nutrient composition, waste products (urea/feces), and gut retention time affect the food and water requirements, but do not directly affect metabolic energy calculations. In the two modeled herbivorous diet scenarios the high-browsing animals display substantial differences in the volume of food required per day relative to low browsing animals. This is due to differences in the %dry mass, where the higher the %dry mass, the greater non-water component is available for digestion (see above, and Fig 8). *Plateosaurus*, as a high browser with ratite RMR and CTR in the cold microclimate meets the calculated target food intake (blue filled pentagon, Fig 8). The lower than target values for high browsing in the hot microclimate demonstrate a decrease in activity below 2 times RMR, likely due to heat stress.

The incremental addition of insulation to *Coelophysis* produced a corresponding decrease in overall food requirement. The uninsulated *Coelophysis* (ratite RMR/CTR) with a carnivorous diet in the hot climate requires ∼300 kg/y, which is near the calculated target food intake requirement of 310 kg/y (blue filled pentagons of Fig. 7). However, with full insulation the annual intake is only 200 kg/y, suggesting heat stress has an impact on activity through a reduction in metabolic heat production during some parts of the year, thus requiring less food intake. Under cold climate conditions the uninsulated *Coelophysis* with a carnivorous diet requires more than twice the target food intake to maintain core temperatures, while a fully insulated individual is slightly heat stressed requiring less than the target food intake.

**Figure 7.**
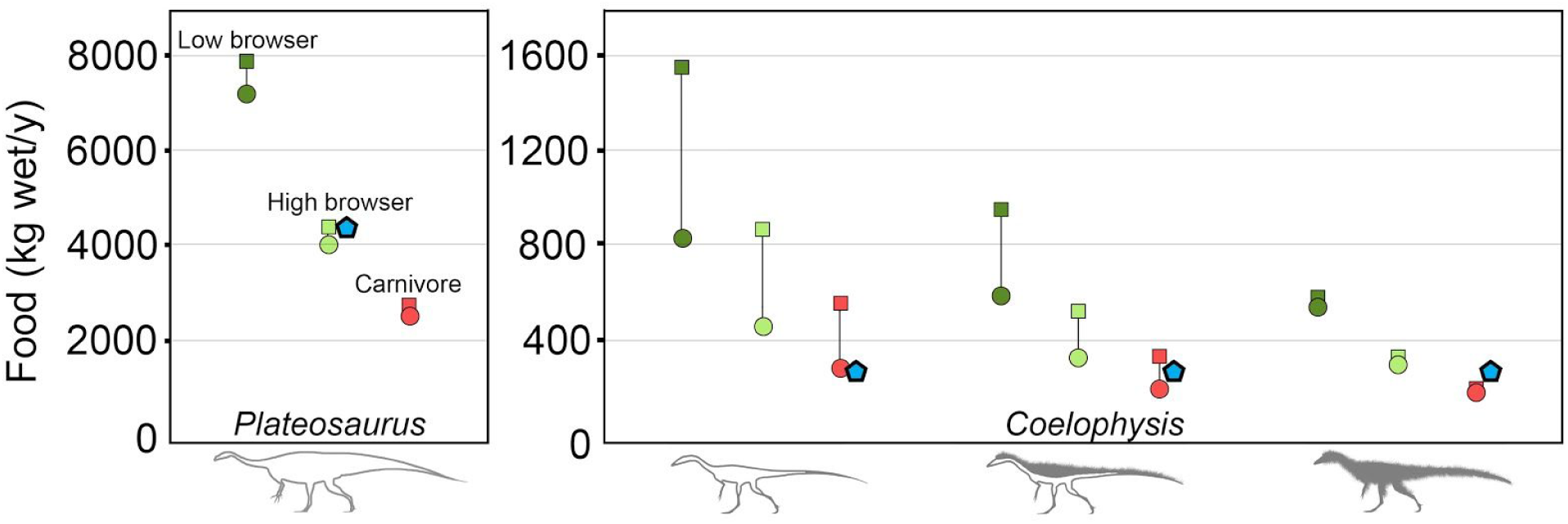
Dietary variability with diet type and insulation (*Coelophysis*). The amount of food needed to maintain the specified (*target*) core body temperature throughout the year varies with diet type. Diet types: low browser herbivore (dark green); high browser herbivore (light green); and carnivore (red). Climate conditions also affect the quantity of food required to maintain core temperatures in hot (closed circles) and cold (closed squares) climates; annual target food intake in kilograms for each species is denoted by a closed blue pentagon when *Plateosaurus* = high browser and *Coelophysis* = carnivore. Data represent each species with a ratite RMR and CTR.

It is notable that the absolute difference between the cold and hot climate annual food requirements decreases non-linearly as insulation increases similar to that reported by Porter [101]. There is a 6% difference in the annual food budget between hot and cold climates for the fully insulated *Coelophysis* and an 8.8% difference for *Plateosaurus* for all diets (carnivorous/herbivorous). In contrast, the difference in annual food budget under cold and hot climates for the top-only insulated and uninsulated *Coelophysis* increases to 36% and 46% respectively for the cold climates relative to warm climates. These differences in food requirements for small dinosaurs with little to no insulation are directly related to the decrease of thermal heat flux from the body due to increased insulation for fully insulated *Coelophysis* or having a large adult body size like *Plateosaurus*.

### Metabolic Chamber Simulations

Within the metabolic chamber simulation that spanned 0-50°C, *Plateosaurus* displayed a greater range of temperatures where it could remain in its active-thermoneutral zone relative to the small bodied uninsulated and top-only insulated *Coelophysis*. The fully insulated *Coelophysis* exhibited a similar breadth of thermoneutrality range as the *Plateosaurus* (Fig. 8). For each stepwise increase in resting metabolic rate (RMR; from squamate to eutherian) two general trends were observed: 1) thermoneutral zone breadth increased and 2) the maximum and minimum thermoneutral temperature values each shifted to lower values. This trend is more apparent in the larger bodied *Plateosaurus*. For example, *Plateosaurus* with a ratite-like core temperature range (CTR) of 38±2°C can maintain thermoneutrality with a squamate grade RMR in air temperatures between 32-45°C and between 15-36°C with a ratite grade RMR. There is an 8°C increase in the absolute thermoneutral range from squamate to ratite RMR, while the maximum air temperature shifts negatively by 9°C.

**Figure 8.**
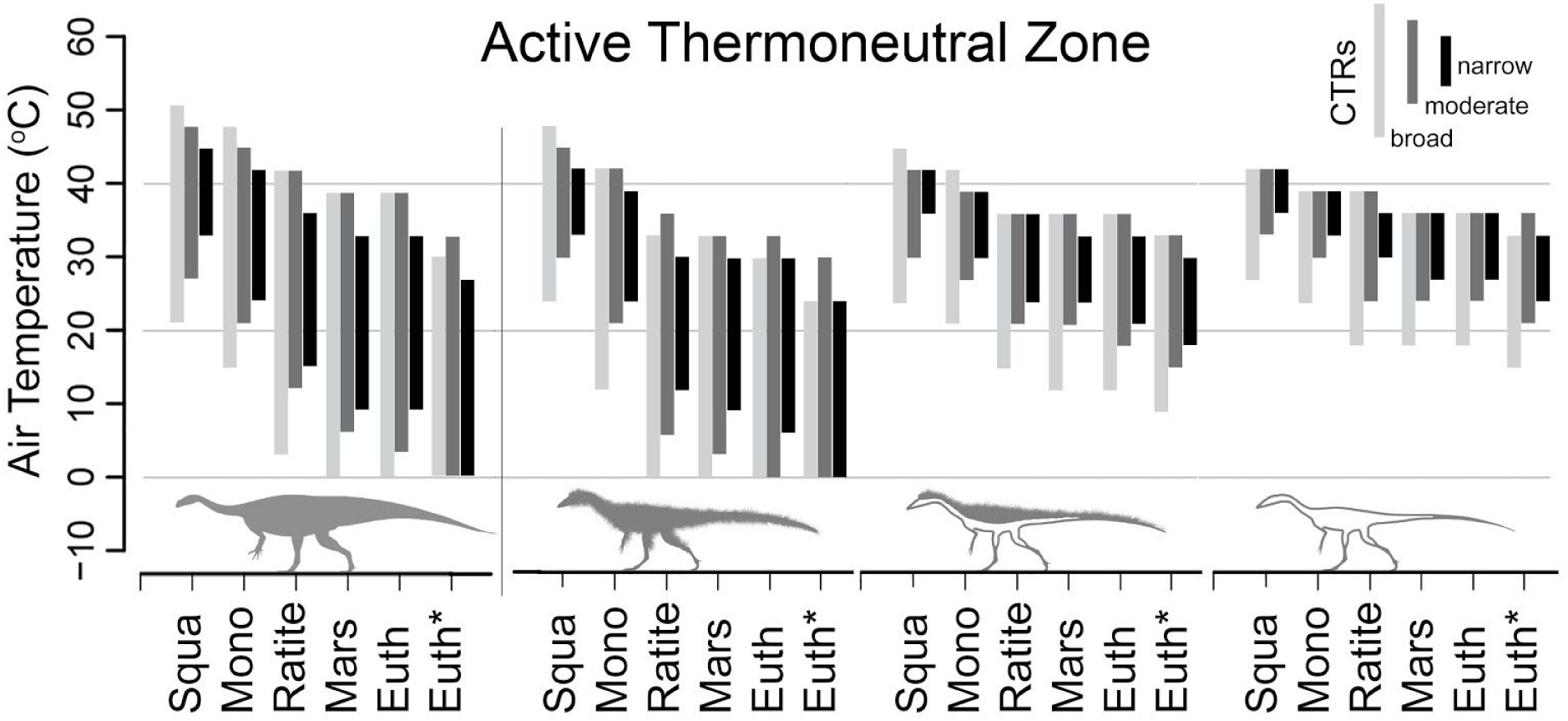
Active thermoneutral zones of *Plateosaurus* and variably insulated *Coelophysis*. Shaded areas represent the active thermoneutral zone determined from 18 metabolic chamber experiments for *Plateosaurus* and *Coelophysis* (fully insulated, top-only, and uninsulated) with RMR ranging from squamates to eutherians based on published regression equations [7,75,76]. Light gray = broad CTR (26-40°C); dark gray = moderate CTR (32-40°C); black = high CTR (36-40°C). Target T_core_ = 38°C. The active thermoneutral zones for the top-only and fully insulated *Coelophysis* were calculated with the ptiloerection behavioral function enabled.

Varying the amount of external insulation in the form of filamentous ‘proto’-feathers made a substantial difference in thermoneutral temperature ranges. An uninsulated *Coelophysis* could maintain thermoneutrality over an 6-10°C temperature range (Fig. 8). With a ratite RMR and CTR (38±2°C) the thermoneutral range of an uninsulated *Coelophysis* was 30-36°C. As dermal insulation was added the overall pattern observed was similar to that seen with increased BMR; i.e., there was a stepwise decrease in maximum and minimum thermoneutral temperature, but an overall increase in total range. The thermoneutral range relative to the uninsulated model was extended moderately 0-3°C (depending on metabolic rate) in the top-only insulated *Coelophysis*. A fully insulated *Coelophysis* had a substantial decrease in the lower end of its thermoneutral range while minimally decreasing its upper bound (12-30°C); the fully insulated *Coelophysis* more than doubled its active thermoneutral air temperature range. The net effect of insulation allows a fully insulated *Coelophysis* to maintain thermoneutrality across a much broader temperature range in colder environments compared to the non-insulated *Coelophysis*, although this is at the cost of lowering the maximum tolerable air temperature.

To test the effect of variable CTRs as well as RMRs we simulated a broad (26-40°C), moderate (32-40°C), and narrow (36-40°C) core temperature range for each of the 6 RMRs (Fig. 8). The same trends were observed with the broad and moderate CTR as seen in the narrow CTR simulations, however the absolute range was greatest in the broad CTR and intermediate in the moderate CTR, and lowest in the narrow CTR discussed above. We also tested the model with four different *target* core body temperatures, 38, 35, 32 and 29°C with a narrow (±2°C) and broad(+2/-13°C) CTR for both species to compare their active thermoneutral zones under these conditions. As the *target* core temperature was stepped down, the overall thermoneutral range remained effectively the same, but the absolute minimum and maximum air temperature values shifted negatively ∼ 2-3°C for each 3°C step down in *target* core temperature. The results for the four target core temperatures under ratite-like and squamate-like CTR are shown in S4 Figures 9 & 10, respectively.

### Effects of resting metabolic rate and core temperature range

Each model simulation paired different physiological combinations of resting metabolic rate (RMR; squamate, monotreme, and ratite grade metabolic rates) with a broad, moderate, or narrow core temperature range (CTR), each with a 38°C target core temperature (26-40°C, 32-40°C, and 36-40°C, respectively), under cold, moderate, and hot climates for the two dinosaur species. The results of these experiments yielded hourly outputs that were plotted as annual heatmaps for core body temperature, metabolic energy (contoured in multiples of RMR), and hours in open versus shaded conditions (see Fig. 9 for explanation of heatmaps).

**Figure 9.**
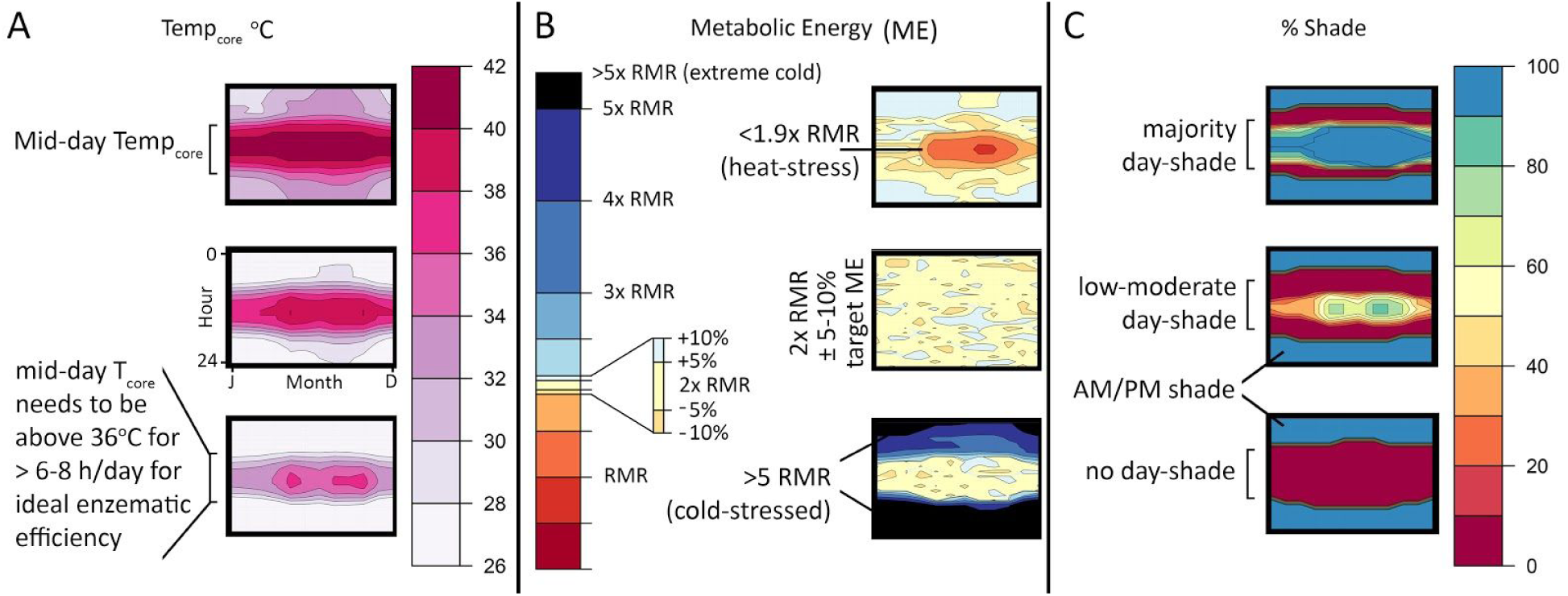
Heatmaps of T_core_, metabolic energy (ME), and % shade. Heatmaps provide a quick quantitative tool for visualizing results on an hourly basis across the year. A) Top; example of narrow CTR heatmap where 36<T_core_<40 ∼6-8 hrs per day. Bottom; 36<T_core_<38 ∼0-3 hrs per day (e.g., cold stressed). B) Top; Metabolic energy (ME) heatmap displaying heat stress during mid-day hours, mid-year. Middle; ME heatmap displaying a reasonable range around 2x RMR. Bottom; results of a cold-stressed model with ME exceeding 5x RMR. C) Heatmaps demonstrating a high (top), moderate (middle), and low (bottom) daylight hours shade requirement.

#### *Coelophysis* (uninsulated)

The uninsulated *Coelophysis* model results show a high degree of cold stress for all but 3 of the 27 possible RMR/CTR/microclimate combinations (Fig. 10). The two best fits are the moderate and upper RMR with broad CTR in the hot microclimate. However, under all RMR/CTR combinations *Coelophysis* is cold stressed in the cold microclimate. Even if paleotemperatures of high latitude localities were only slightly cooler (moderate) than equatorial (hot) conditions modeled herein, the uninsulated *Coelophysis* still shows signs of cold stress (i.e., T_core_ does not reach 35°C for more than 4 hours a day, for over half of the year; 3 months of the year never reach 35°C at all).

**Figure 10.**
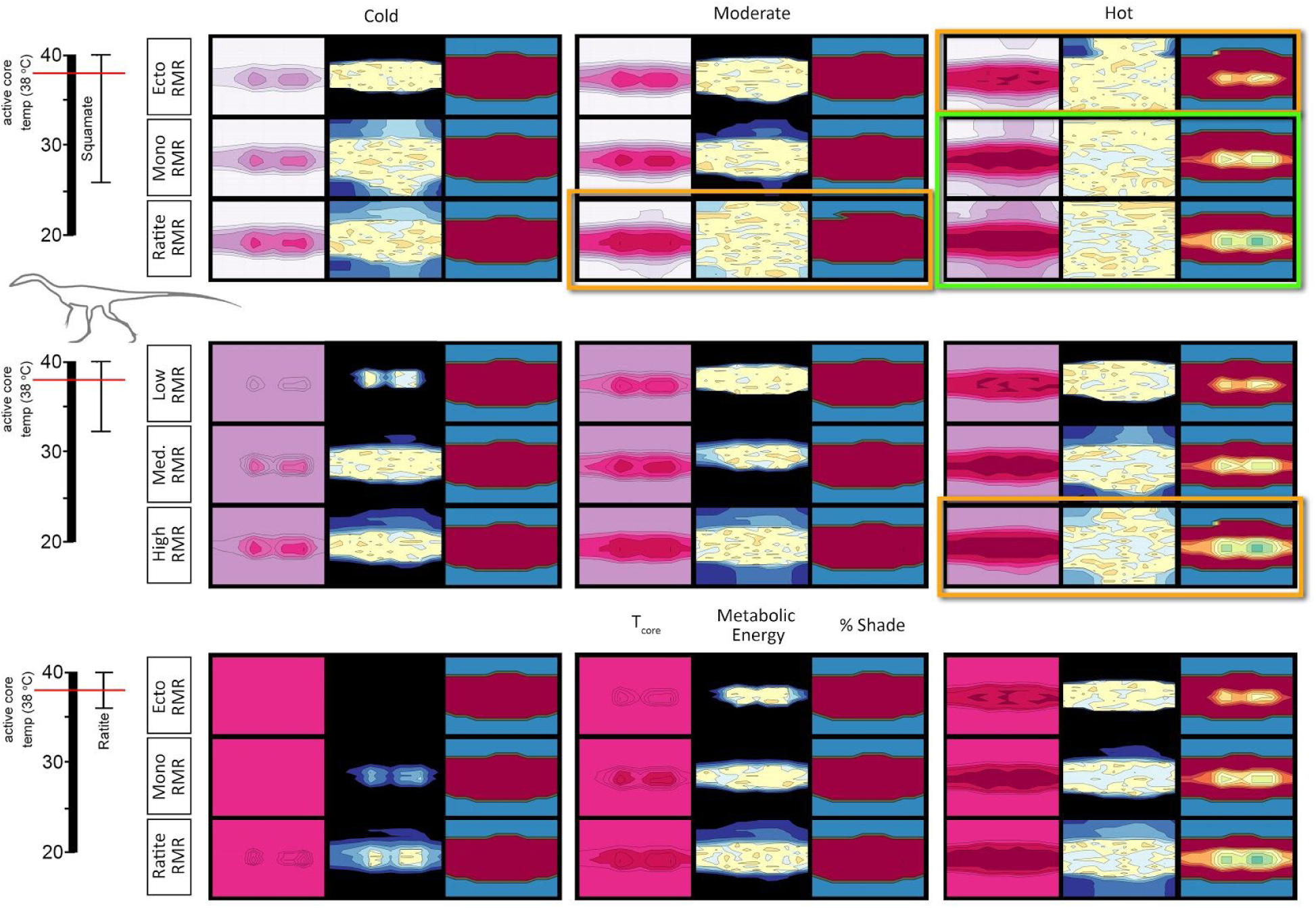
T_core_, ME, and %shade heatmaps for *Coelophysis* (uninsulated). Heatmaps representing the hourly results across the modeled year the three dominant variables: microclimate (hot, moderate, and cold), RMR (low, moderate, and high), CTR (broad, moderate, and narrow) for an uninsulated *Coelophysis*. See figure 10 for key. Two most likely scenarios for survivability are outlined in bright green, the three edge conditions are outlined in orange; all other conditions are considered to be non-viable.

Because many ectothermic animals have the potential to decrease their internal temperatures below the 26°C lower bound we used in the broad, squamate-grade CTR, we also modeled the uninsulated *Coelophysis* with a 10°C lower temperature bound to ensure we capture the lowest extremes of core temperature. *Coelophysis* was modeled in the hot and cold microclimate for the month of May (northern hemisphere early summer). These data were plotted along with the November (southern hemisphere early summer) temperature profile for the largest known extant ectothermic terrestrial vertebrate, *Varanus komodoensis*, as a frame of reference (Fig. 11). In the hot microclimate T_core_ for *Coelophysis* responded similarly to *V. komodoensis*. In the cold microclimate T_core_ does not exceed 30°C for more than 5 months of the year demonstrating severe cold stress (non-viable).

**Figure 11.**
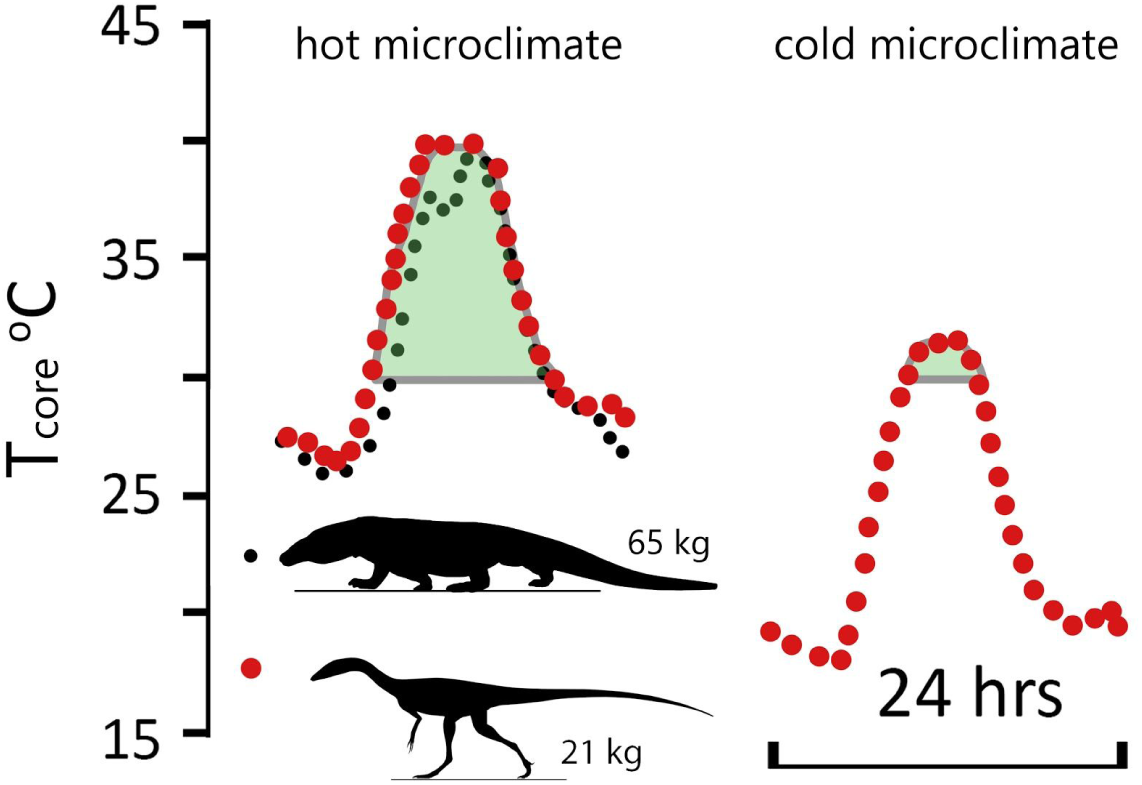
Comparison of daily temperature curves for *Varanus* and *Coelophysis* (uninsulated). Daily temperature curves for hot and cold microclimates for the 15th of May (uninsulated *Coelophysis*) and November (*Varanus komodoensis*) [40]. There is strong agreement between the low RMR and broad CTR *Coelophysis* and *V. komodoensis* in the hot microclimate. *Coelophysis* modeled in the cold microclimate was significantly cold stressed. Green shaded area represents duration of day with T_core_ > 30°C.

With a 10-40°C CTR the lowest ambient air temperature in the cold microclimate was above the lower (10°C) CTR threshold, thus, it was possible for the organism to thermoregulate and maintain its target ME by dropping its core temperature rather than increase its metabolic rate (see S4 Figure 11). This did not affect the daily core temperature result between 26 and 40°C which are identical as the prior broad CTR experiment above; the animal is still significantly cold-stressed.

#### *Coelophysis* (top-only insulation)

With the addition of insulation to the top-half of *Coelophysis* the severity of cold-stress decreased and the number of viable RMR/CTR/microclimate combinations increased to 6 of 27 (Fig. 12). Under the hot microclimate with a broad CTR, all RMR conditions met ME targets and were able to maintain a core temperature above 35°C. As the microclimate shifted to the moderate condition, the lower RMR was excluded; all RMR were excluded under the cold microclimate. As the CTR reached the moderate range, only the moderate and upper RMR were considered feasible under the hot microclimate. The narrow CTR excluded all RMR in all microclimates.

**Figure 12.**
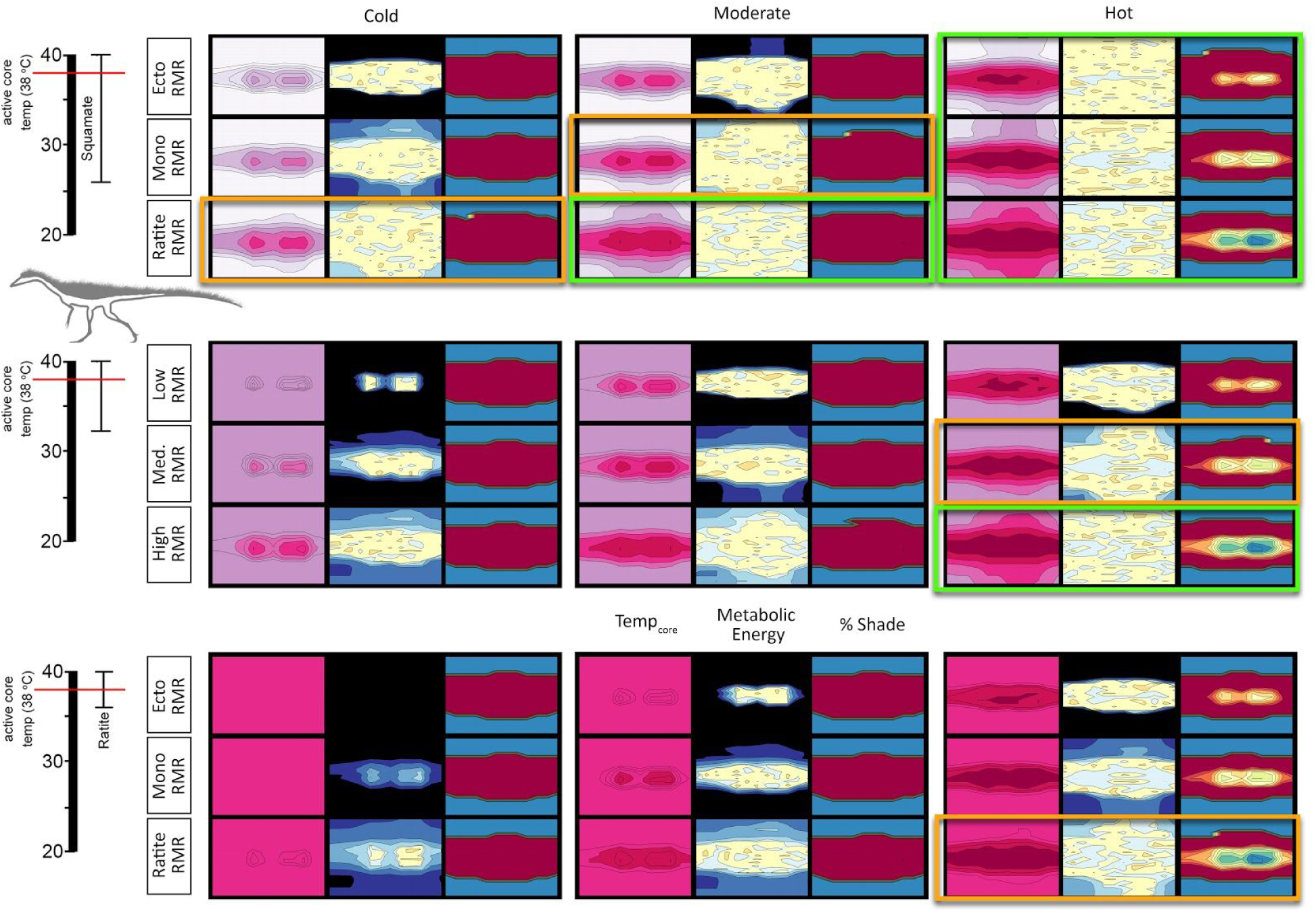
T_core_, ME, and %shade heatmaps for *Coelophysis* (top-only insulated). Heatmaps representing the hourly results across the modeled year the three dominant variables: microclimate (hot, moderate, and cold), RMR (low, moderate, and high), CTR (broad, moderate, and narrow) for an top-only insulated *Coelophysis*. See figure 10 for key. Five most likely scenarios for survivability are outlined in bright green, the four edge conditions are outlined in orange; all other conditions are considered to be non-viable.

#### *Coelophysis* (full insulation)

With a fully insulated *Coelophysis* the severity of cold-stress further decreased and the number of viable RMR/CTR/microclimate combinations increased to 10 of 27; heat stress was evident in all 3 CTRs with an upper RMR under the hot microclimate (Fig. 13). The fully insulated *Coelophysis* was cold stressed in the cold and moderate microclimates with a broad CTR. However, it was able to maintain its ME target and sustain a core temperature greater than 34-36°C for at least half of dial hours with a broad CTR and: lower RMR in the hot microclimate; moderate RMR in moderate and hot microclimates; upper RMR in cold and moderate microclimates. Raising the CTR to the moderate condition excluded all lower RMRs as well as the moderate RMR condition in the moderate microclimate. A narrow CTR resulted in the loss of the remaining moderate RMR in the hot microclimate; only the upper RMR in moderate and cold microclimates were able to meet their ME target.

**Figure 13.**
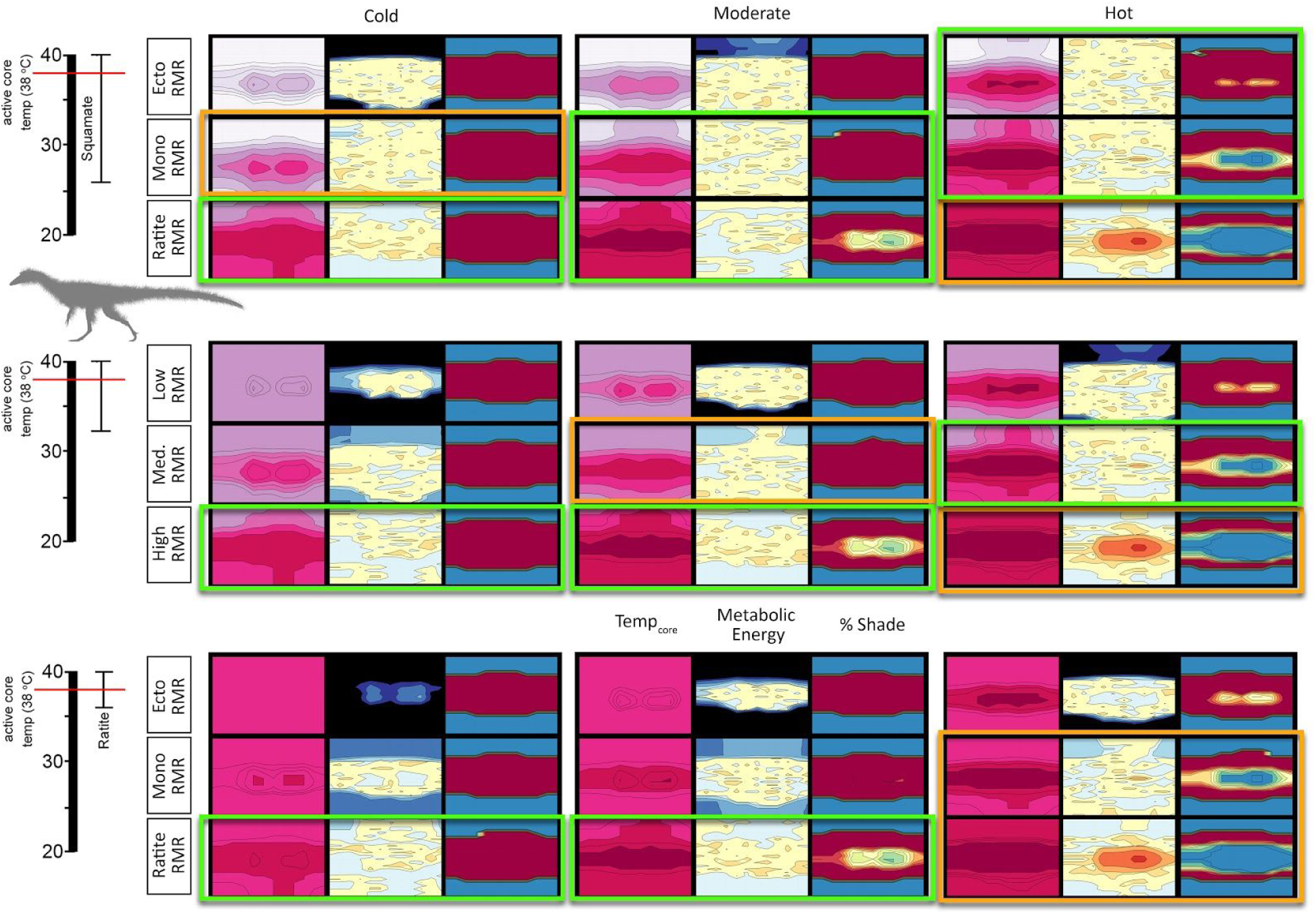
T_core_, ME, and %shade heatmaps for *Coelophysis* (fully insulated). Heatmaps representing the hourly results across the modeled year the three dominant variables: microclimate (hot, moderate, and cold), RMR (low, moderate, and high), CTR (broad, moderate, and narrow) for a fully insulated *Coelophysis*. See figure 10 for key. Tenmost likely scenarios for survivability are outlined in bright green, the six edge conditions are outlined in orange; all other conditions are considered to be non-viable.

#### Plateosaurus

*Plateosaurus* exhibits a similar response to that seen in the fully insulated *Coelophysis*; the number of viable RMR/CTR/microclimate combinations was 10 of 27; heat stress was evident in all 3 CTRs with a ratite RMR under the hot microclimate (Fig. 14). With a lower RMR and broad CTR, *Plateosaurus* was cold stressed in the early morning hours under cold conditions and didn’t exceed 30°C body temperature for more than half of the calendar year. The moderate microclimate fared only slightly better, but the ME still exceeded target by ∼10%. This same physiology modeled in the hot microclimate demonstrated a core temperature of 28-30°C during morning hours and reached target core temperatures by midday.

**Figure 14.**
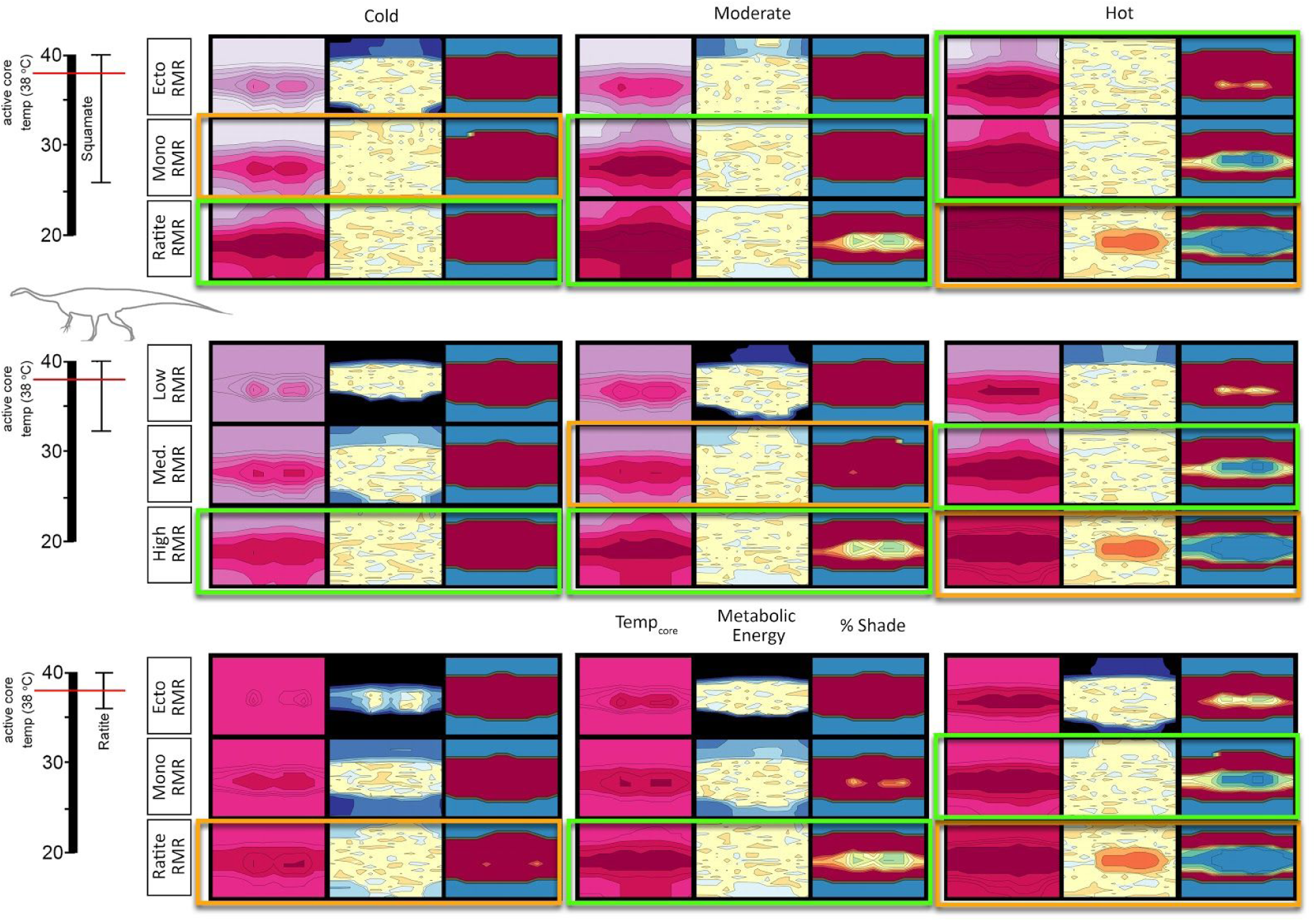
T_core_, ME, and %shade heatmaps for *Plateosaurus*. Heatmaps representing the hourly results across the modeled year the three dominant variables: microclimate (hot, moderate, and cold), RMR (low, moderate, and high), CTR (broad, moderate, and narrow) for a *Plateosaurus*. See figure 10 for key. Ten most likely scenarios for survivability are outlined in bright green, the six edge conditions are outlined in orange; all other conditions are considered to be non-viable.

When the CTR reached the moderate level, all lower RMR were excluded due to significant cold stress, as was the moderate RMR in the cold microclimate. The moderate RMR met its ME target in the hot microclimate, but its ME slightly exceeded its target goal in the moderate microclimate and exceeded its ME target under the cold microclimate. The final step to a narrow CTR increased the cold stress previously observed in the moderate and cold microclimate with a moderate RMR, the model slightly exceeded its ME target under the cold microclimate with an upper RMR. The model met its ME target under the moderate microclimate with an upper RMR.

### Microclimate wind effects

Because the wind was the second strongest main effect in our yates analysis (see S4) we further explored this effect using *Coelophysis* and *Plateosaurus* with an upper RMR and narrow CTR. For *Coelophysis*, the magnitude of wind effects varies substantially depending on the degree of insulation, ptiloerection, and climate (Fig. 15). Daily variation in wind speeds from 0.1 to 2.0 m/s affects total annual energy requirements from approximately 2000 (hot microclimate) to 3400 MJ/y (cold microclimate) without insulation down to approximately 1400 (hot microclimate) to 1800 MJ/y (cold microclimate) when fully insulated *without* ptiloerection (30 mm insulation depth); the fully insulated (with ptiloerection enabled) was ∼1500 MJ/y. Ptiloerection was not activated until the model required >2x RMR to maintain *target* core temperatures.

**Figure 15.**
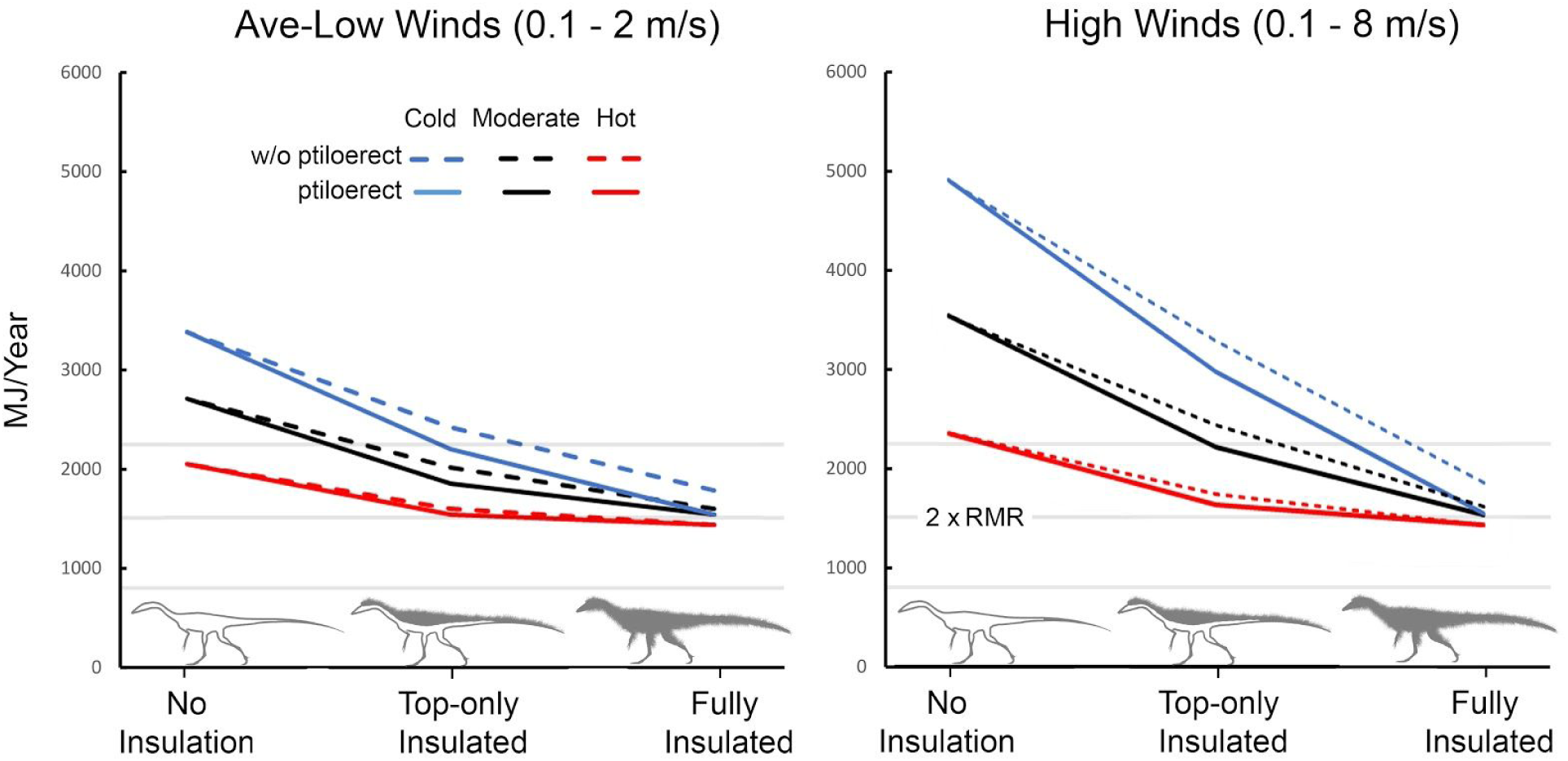
Energetic cost of wind exposure for *Coelophysis*. As temperature increases (blue, black, and red lines, respectively) ptiloerection was less beneficial with increased insulation volume. (e.g., fully insulated *Coelophysis* does not significantly benefit by implementing ptiloerection at hot temperatures but the presence of feathers broadens its active thermoneutral zone). The three light gray horizontal lines represent, from bottom to top, resting, twice resting (e.g., ME target), and three times resting metabolic rate to indicate the likely range of activity levels for the size, shape, and degree of insulation for *Coelophysis*.

We tested 5 insulatory conditions for each climate: 1) no insulation, 2) 15 mm depth top half only (only the top half of the animal had insulation), 3) 30 mm depth top half only with, 4) 15 mm depth fully insulated, and 5) 30 mm depth fully insulated. Fully insulated animals only engaged ptiloerection in the coldest microclimate. When the feather depth of the fully insulated animal decreased from 30 to 15 mm its energetic response was similar to that of the 30 mm top-only insulation; thus decreased insulation depth is equivalent to greater depth with only top surfaces insulated.

Wind did not have as large of an impact on the modeled *Plateosaurus*, although wind was the second strongest effect observed in the Yates analyses. *Plateosaurus* was able to maintain its target core temperature under the moderate and hot microclimates for average and high speed winds. In low speed winds moderate and cold microclimates were at the lower *target* boundary (-5% of 2x RMR), while the hot microclimate caused notable heat stress (-10% of 2x RMR; Fig. 16). This stresses the importance of behavior for the model to seek shelter from or take advantage of higher wind conditions for thermoregulation.

**Figure 16.**
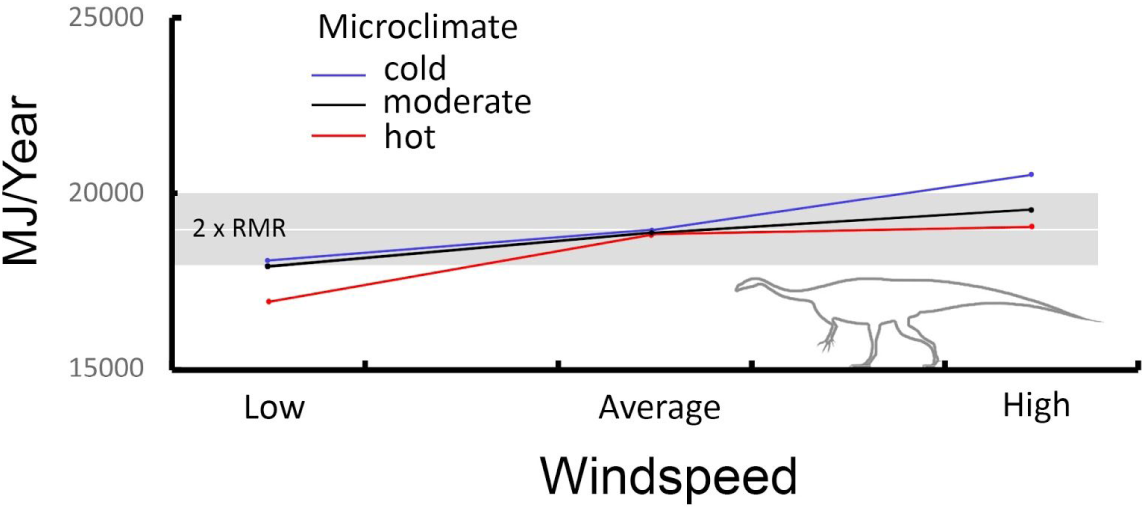
Energetic costs of wind exposure for *Plateosaurus*. Under low wind speeds *Plateosaurus* is moderately to notably heat stressed (cold/moderate and hot, respectively). *Target* ME is maintained in all microclimates for average winds, and in the hot microclimate with high winds. *Plateosaurus* becomes cold stressed with high winds in the cold microclimate. Red line = hot microclimate, blue line = cold microclimate, black line = moderate microclimate.

## Discussion

Modeling an organism’s interaction with its environment effectively predicts the environmental range of modern animals with high fidelity [19,20,31,32,69]. This has been leveraged to generate hypotheses of how organisms respond to habitat expansion, contraction, and altered geographic ranges associated with changing climate on local and global scales [12,18,40,102]. Our efforts have focused on extending this tool to test biophysical hypotheses in deep time.

Niche Mapper has demonstrated the ability to predict metabolic expenditure as a function of environmental conditions for a broad sample of vertebrates in microclimates that range from arctic to tropical [13,17,20,32,34]. This study extends Niche Mapper’s applicability to define the range of physiological conditions under which extinct animals functioned within a designated microclimate. While we may not have a detailed empirical physiological profile for extinct animals, this approach allows us to explore different combinations of morphological and physiological options to determine how metabolic rates and geometries would have affected energetics, behavior, and animal distributions for a given microclimate.

We leveraged a variety of stable isotopic and geochemical systems to infer mean annual temperature, mean annual precipitation, atmospheric O_2_ and CO_2_, and relative humidity in the Late Triassic of Western North America [55,103–105],. These and other sedimentary paleoenvironmental proxies [e.g., 52] were employed to validate global climate models. Our analyses can also be decoupled from specific microclimate models by using Niche Mapper’s virtual metabolic chamber function to determine an active thermoneutral temperature range for modeled taxa. Determining the overlap in thermoneutral zones of organisms and mapping them against their paleobiographic distributions [e.g., 15, 23] provides and independent test of plausible paleophysiology.

### Plateosaurus

Our results demonstrate that *Plateosaurus* could have maintained its target metabolic energy (ME) in hot environments with either a squamate-like core temperature range (CTR) and resting metabolic range (RMR), or with a monotreme-like RMR at moderate to narrow CTR. A shift from hot to moderate or colder environments required at minimum a ratite-like RMR with a moderate CTR. Modeling *Plateosaurus* with a ratite-like narrow CTR and upper RMR resulted in heat stress in hot environments, full viability in moderate environmental temperatures, and slight cold stress in colder environments (Fig. 17).

**Figure 17.**
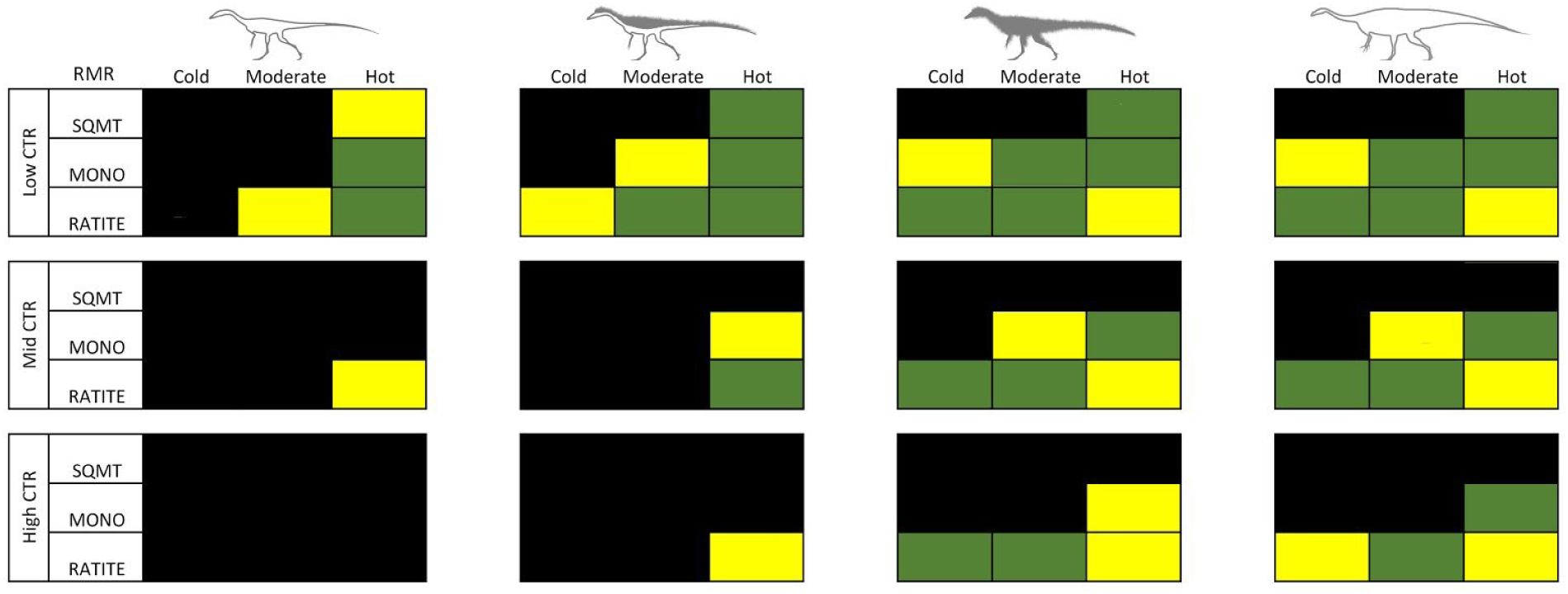
Summary of viable, conditional tolerance, and non-viable results. The matrix provides a summary of viable combinations of resting metabolic rate (RMR) and core temperature range (CTR) within cold, moderate, and hot microclimates. Green = viable; black = non-viable; yellow = conditional tolerance (e.g., a possible but extreme endmember of viability).

Modeling *Plateosaurus* with a squamate RMR was non-viable for all moderate and cold microclimates regardless of CTR. The greater viability of *Plateosaurus* with elevated RMRs in Late Triassic environments is consistent with isotopic estimates derived from dinosaur teeth and eggs which suggests an elevated core temperature between 36-38°C for the sauropod lineage [106–110]. Additionally, a squamate-like broad core temperature would be pushing the lower limits for enzymatic efficiency in large herbivores. This would translate to an inhibition of rapid growth, which is counter to rates of growth reported from *Plateosaurus* bone histology [111, 112].

Considering the paleogeographic range of *Plateosaurus* and it’s close relatives (see discussion below) extend well into temperate latitudes, squamate-level RMRs and CTRs appear to be non-viable. If *Plateosaurs* was trying to defend a highly stable internal core temperature a ratite-like RMR and narrower CTR would be required, in addition to physiological acclimatization such as seasonally variable metabolic rates, variable fat stores, or changes in conductivity to the ground as found in other endothermic animals [e.g., 113, 114]. We conclude that *Plateosaurus* most likely had a moderate core temperature range coupled with an elevated ratite-like resting metabolic range.

### Coelophysis

When modeled without insulation and possessing either a narrow or moderate CTR, *Coelophysis* is non-viable due to excessive cold stress in all environments regardless of RMR. Non-insulated *Coelophysis* modeled with a broad CTR is able to meet its target needs in hot environments with either a monotreme or ratite-like RMR, but remains non-viable due to cold stress under all other conditions. The daily temperature profile for the month of May with a squamate-like RMR and CTR is strikingly similar to that seen for a small adult komodo dragon (e.g., Fig. 11). Winter months show minor cold stress for this condition; this can be alleviated by decreasing the lower CTR bound by as little as 2°C, although decreasing the CTR range does not alter peak temperatures and *Coelophysis* remains cold stressed in the moderate and cold environments as a squamate. In short, a non-insulated *Coelopyhsis* would be viable only in hot environments with a broad CTR, regardless of RMR.

When modeling epidermal insulation along the top half of the neck, torso, and tail leaving the ventral surfaces bare, the modeled *Coelophysis* saw increased capacity to maintain its daily target ME in moderate environments with a broad CTR and ratite-like RMR. All RMRs were viable in the hot microclimate with a broad CTR. The addition of dorsal insulation made a moderate CTR viable, but only with a ratite-like RMR within hot environments. Half-insulated *Coelophysis* was still non-viable in all cold environments regardless of CTR.

The final biophysical scenario, the fully insulated *Coelophysis*, was viable in a broader range of temperatures. A fully insulated *Coelophysis* was only viable with a squamate-like RMR in the hot environment with a broad CTR. Modeling a monotreme-like RMR resulted in non-viability due to cold stress in colder environments regardless of CTR; *Coelophysis* with a monotreme-like RMR was viable in hot (moderate to broad CTR) and moderate (broad CTR only) microclimates. Cold and moderate environments were accessible to the fully insulated *Coephysis* with moderate to narrow CTRs with a ratite-like RMR, although the model faced heat stress during peak summer temperatures in the hot microclimate.

Theropod isotopic paleothermometry indicates elevated RMRs and core temperature ranges above the levels of extant squamates [107,109,115]. Eagle and others [107] performed a clumped-isotope analysis on oviraptorosaur eggs and concluded the egg-layer had an average core temperature of around 32°C. Oviraptorosaurs had a mass broadly similar to *Coelophysis*, though they were significantly more derived theropods (e.g., more deeply nested within Coelurosauria); the analyzed specimens were found in deposits with a high paleolatitude (> 45°N) during time of deposition [116]. An average core temperature much lower than 32°C would likely inhibit metabolic efficiencies necessary for elevated growth rates reported for *Coelophysis* [117, 118].

Lowering the *target* core temperature from 38°C to 32°C resulted in an average (viable) core temperature of 32°C for *Coelophysis* modeled with squamate-like RMR and CTR within hot environment models only; under both moderate and cold microclimates a squamate-like *Coelophysis* becomes non-viable due to extreme cold stressed (with T_core_ rarely exceeding 30°C for more than a few hours a day, S4 Figure 12 & Table 1). The half-insulated *Coelophysis* with a ratite-like RMR and CTR was viable in moderate to hot environments. With a ratite-like RMR fully insulated *Coelophysis* viability increased to include cold and moderate environments; it was still heat stressed in the hot microclimate. Results from both elevated (38°C) and lowered (32°C) *target* core temperatures are consistent with viability in cold to hot environments with a ratite-like RMR as a fully to half insulated *Coelophysis* (S4 Table 1). These results support an elevated RMR for *Coelophysis* with a moderate to narrow CTR, and suggest that *Coelophysis* did not need to vigorously defend an elevated stable (i.e., narrow) core temperature.

### Integrating model results with the fossil record

Body fossils of coelophysoids are known from equatorial through temperate paleolatitudes, while larger bodied plateosaurid body fossils are known from subtropical and temperate climates, but are notably lacking in equatorial paleolatitudes (Fig. 18). Given that the paleogeographic range of the two modeled species extends to temperate latitudes we used our cold microclimate model as a conservative surrogate for temperatures at subtropical to temperate latitudes [see 55,59,60]. The results of *Plateosaurus* modeled at 45°N paleolatitude demonstrates that air temperature has a much greater effect than does increased variability in insolation, supporting the use of our cold microclimate (modeled at lower latitude) as an analog for temperate latitudes.

**Figure 18.**
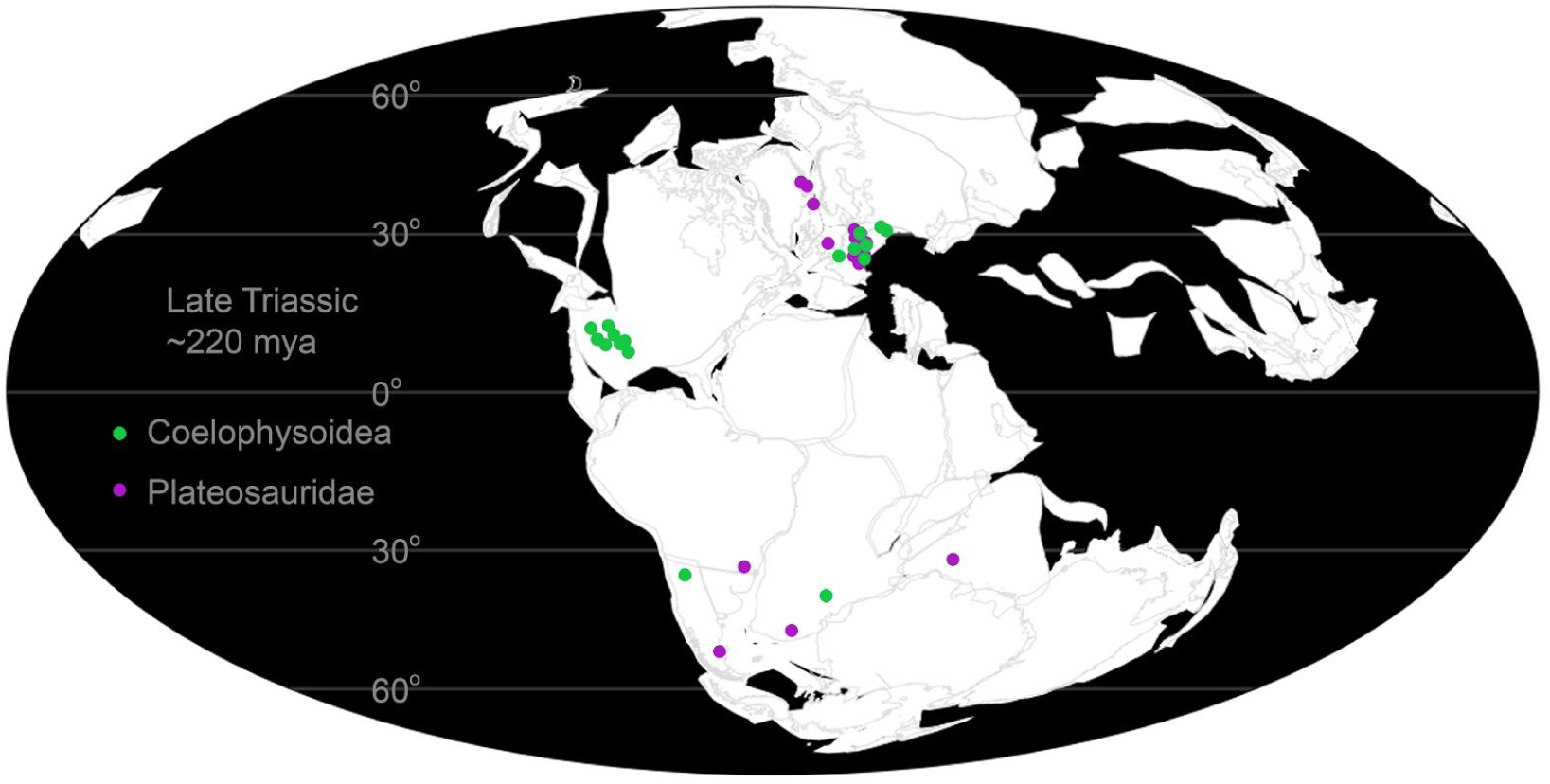
Paleogeographic distribution of body fossils for members of Coelophysoidea and Plateosauridae. Note the absence of Plateosauridae at tropical latitudes. Data from paleobiodb.org.

Examining fossil distribution patterns, there is a gap extending through tropical latitudes where skeletons of large bodied *Plateosaurus* and other large Late Triassic prosauropods (e.g., *Antetonitrus* (Yates and Kitching 2003), *Unaysaurus* (Leal and others 2004), and *Efraasia* (von Huene 1908)) are not represented in the fossil record. Conversely, large bodied dinosaurs are well known from cooler subtropical to temperate latitudes, consistent with our results for *Plateosaurus* in moderate to cooler environments.

Although no body fossils have been found, the Late Triassic vertebrate track record of North America contains traces that have been attributed to medium-sized prosauropods (i.e., *Evazoum* (Nicosia and Loi 2003) formerly *Pseudotetrasauropus* (Ellenberger 1965), see [119]). The trackmakers would have been similar in size to the Early Jurassic skeletons of *Seitaad* (Sertich and Loewen 2010) and *Sarahsaurus* (Rowe and others 2011). There remains some doubt that the taxonomic identities of the trackmakers are dinosaurian at all [119]. However, tracksites from northeastern New Mexico have been more confidently attributed to a larger sauropodan dinosaur [119]. Notably, the locations of all described prosauropod/sauropodan trackways are near the edges of local highlands on the Late Triassic landscape (Fig. 19).

**Figure 19.**
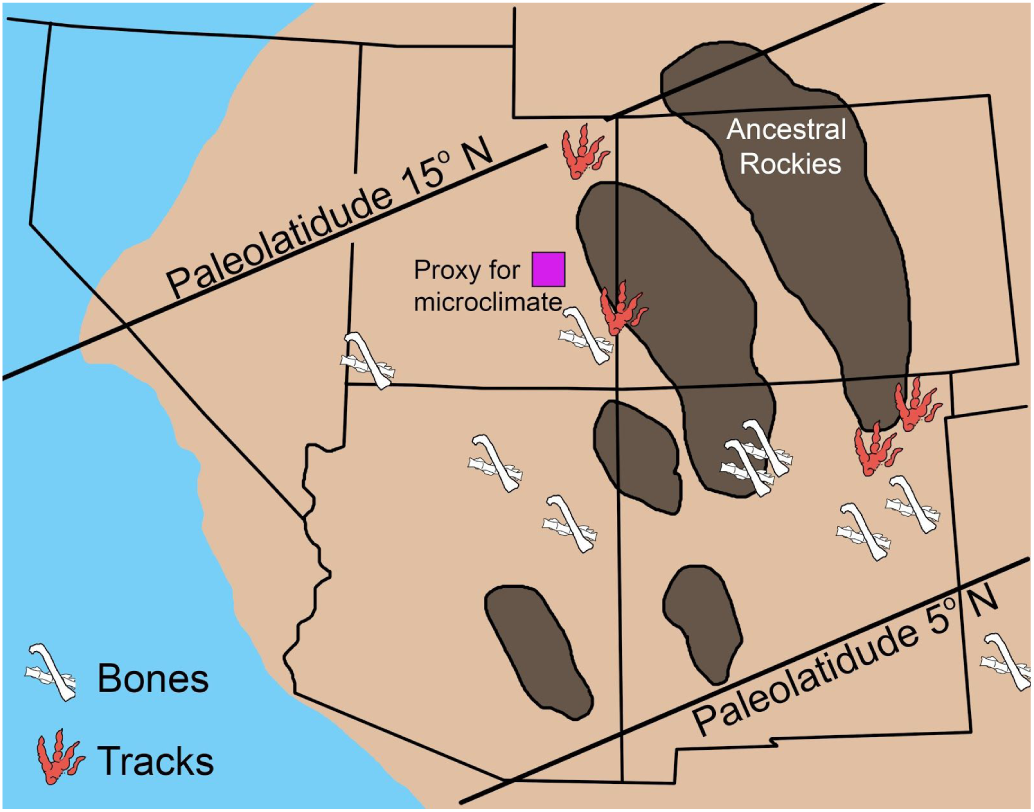
Track locations attributed to prosauropods and bones of *Coelophysis* in the late Triassic of the western USA. Purple square = field localities from which proxy microclimate data used in this study were previously published [52]. Bones = localities with known coelophysoid body fossils.

Our results for *Plateosaurus* modeled in hot environments indicate heat stress was the dominant constraint restricting *Plateosaurus* to cooler environments. Thus, we suggest environmental temperatures as the primary mechanism limiting prosauropods from greater utilization of tropical latitudes during the Late Triassic. Within lower latitudes prosauropods may have been present in cooler environments such as forested areas or higher elevations, but these depositional environments are less conducive to fossil preservation compared to the hotter environments encountered in lowland floodplains of the Chinle Formation [120, 121]. Utilizing dense vegitative cover or higher elevations would explain both the absence of body fossils in Chinle floodplain deposits and the rare occurrence of trackways attributed to prosauropod-like track makers surrounding elevated cores of the ancestral rockies (Fig. 19).

Large size alone is not a limiting factor for the Late Triassic Chinle paleoecosystem. Several large bodied vertebrates inhabited this region, including the ∼1000 kg dicynodont *Placerias* (Lucas 1904) and the ∼2000 kg phytosaur *Rutiodon* (Emmons 1856). Both of these taxa are capable of terrestrial locomotion but are thought to exhibit hippo-like and crocodile-like aquatic behavior and niche occupation respectively [122–124]. Spending time in water enhances heat dissipation and allows for their large size in hot equatorial climates [e.g., 40].

### Insulation in Triassic theropods

In contrast to the skeletal and ichnological fossil record of prosauropods at tropical latitudes, early theropod occurrences are more numerous. The 21 kg *Coelophysis* is well known from the body fossil record of the Chinle Formation (Fig. 19), and other coelophysoids are known globally at higher latitudes (Fig. 18).

The basal saurischian *Chindesaurus* (Long and Murray 1995) which at the least filled a theropod-like ecological role, basal theropods *Daemonosaurus* (Sues and others 2011) and *Tawa* (Nesbit and others 2011), and neotheropod *Camposaurus* (Hunt and others 1998) is known from body fossils within the Chinle Formation; there are abundant trackways attributable to theropod dinosaurs throughout the region. According to several independent methods of mass estimation [e.g. 94, 100] adult theropod taxa from this formation would have been in the 20 kilogram weight range. Body fossils of coelophysoids such as *Zupaysaurus* (Arcucci and Coria 2003), *Liliensternus* (Welles 1984), *Procompsognathus* (Fraas 1913), and *Coelophysis rhodesiensis* (Raath 1969) are known from subtropical to temperate paleolatitudes. *C. rhodesiensis* was discovered in temperate southern paleolatitudes and is similar in size to Chinle coelophysoids. *Zupaysaurus* and *Liliensternus* would have been around 100 kilograms heavier than *Coelophysis*.

*Coelophysis* and other primitive theropods had a bipedal upright stance and a narrow, laterally compressed body that reduces solar cross-section when the sun is overhead and increases surface to volume ratio, enhancing radiative cooling relative to more round-bodied taxa [11]. When the sun was low in the sky laterally compressed animals have greater behavioral flexibility in adjusting their solar radiation cross-section, either by facing towards the sun to minimize their cross-section or by orienting themselves perpendicular to the sun, making their solar absorption equivalent to more rotund organisms. In both cases the change allows for a reduction in solar absorption during peak thermal stress, allowing for higher metabolic rates during the day [101]. Our model results show this bauplan is appropriate in warm environments, but without insulation individuals would have been at a distinct disadvantage in cooler climates. A major difference between our *Plateosaurus* and *Coelophysis* models is the inclusion of three aforementioned states of dermal insulation in the form of primitive filamentous structures for *Coelophysis*. Filamentous and/or quill-like structures are known in a wide range of coelurosaurian theropods, basal ornithischians such as *Tianyulong* (Zheng and others 2009) and *Kulindadromeus* (Godefroit and others 2014), as well as the more derived ornithischian *Psittacosaurus* (Osborn 1923). Recent evidence suggests the origin of primitive feathers extends beyond Dinosauria to the base of Ornithodira (i.e the most recent common ancestor of pterosaurs, dinosaurs and all their descendants; [125]). This supports the idea that insulatory epidermal structures were a plausible thermoregulatory solution for *Coelophysis* [126–128] despite the absence of skin impressions in basal theropods. It should be noted that there are no prosauropod body (or trace) fossils that demonstrate insulatory epidermal structures. It is likely that that these structures were lost as sauropodomorphs increased in size - increased mass alone can expand tolerance of cooler temperatures and stabilize internal temperature variation, but not without its own energetic costs. Similarly, epidermal insulatory structures are known from some (phylogenetically disparate) ornithischians, however several smaller members of this clade from temperate latitudes are known or suspected burrowers [129, 130]. Fossorial behavior allows for the exploitation of more stable microhabitats in environments with higher variance in daily or annual air temperature.

Skeletal remains of *C. bauri* (our model) are well known from Chinle deposits best represented by our hot microclimates, and similarly sized *C. rhodesiensis* is well known from the Elliot Formation (Zimbabwe) which was deposited in a temperate southern hemisphere paleolatitude. Thus, the paleogeographic range of small-bodied coelophysoids extends from northern low tropical latitudes through temperate latitudes of both hemispheres. This is a broad latitudinal and environmental range for coelophysoids in the 20 kilogram size range. Given the lack of evidence of significantly different metabolic adaptations in these closely-related basal theropods, any biophysical scenario must satisfy both the hot and cold microclimates. While no single biophysical condition (e.g., combination of CTR/RMR) satisfies the disparity in climate regimes that *Coelophysis* inhabited, varying the amount and location of insulation covering its body solves this apparent paradox.

The addition of complete insulation coverage of filamentous structures to our modeled *Coelophysis* produced similar results to those of the much larger non-insulated *Plateosaurus*. Our results demonstrate that both *Plateosaurus* and the fully insulated *Coelophysis* would have been heat stressed in the hot Chinle microclimate, limiting their distribution to temperate and boreal latitudes or high elevations (or dense forested areas) at more equatorial latitudes as mentioned above.

In extant taxa the density of insulatory structures vary with ontogeny and with seasonal variability [126,131,132]. Temperature acclimatization to both hot and cold climates is a well-documented phenomenon in birds and mammals. It has been shown that cold-acclimated birds can have greater feather density, higher resting metabolic rates, and reduced evaporative cooling compared to heat-acclimated birds of the same species [see 114].

Recent studies of growth and postnatal development in early dinosaurs (e.g., *C. bauri* and *C. rhodesiensis*) and dinosauromorphs (e.g., the silesaurid *Asilisaurus* (Nesbitt and others 2010)) suggest high variation in developmental sequence and body size at skeletal maturity was likely the ancestral condition [127, 133]. This differs from the moderate to low intraspecific variation in growth seen in extant archosaurs [133]. Griffin and Nesbitt [133] suggest anomalously high variability in *Coelophysis* body size at skeletal maturity may be epigenetically controlled. Higher variability in metabolic rate and core temperature range (and thus physiological efficiencies such as digestion) is consistent with increased variability in size at skeletal maturity. This variability, as well as evidence of increased respiratory efficiency [134–136] and locomotion energetics [137–139] suggests increased RMR within dinosaurs (or their immediate ancestors) was linked to a need to increase aerobic scope, as opposed to enzymatic efficiency, parental care, or increased eurythermy [e.g., 140–142].

Mechanistic modeling of physiological and environmental conditions to test for viable physiological combinations in multiple environments is a relatively new tool for deep time applications. The simulations described above outline the primary components necessary for exploring paleophysiology in deep time with Niche Mapper. The relative effect of temperature, resting metabolic rate, core temperature range, size, and epidermal insulation are much greater than those of skin/fur/feather color, or muscle, respiration, and digestive efficiencies. We do not suggest these other parameters are trivial, rather they are more suited to ‘fine tuning’ the model, such as the extant examples previously mentioned or where circumstances are favorable for a given extinct taxa [i.e., 102]. Niche Mapper is a powerful tool that can be leveraged to address a diverse array of evolutionary questions in deep time such as testing hypotheses pertaining to our understanding of paleoecological carrying capacity, paleobiogeographic distribution, and survivorship across major extinction boundaries.

## Conclusions

Mechanistic models in Niche Mapper use phylogenetically-constrained physiological parameters to determine habitable microclimates for a given taxon. Based on our results, prosauropods like *Plateosaurus* would have had a resting metabolic rate close to that of modern ratites, although we cannot rule out the variance in this clade’s core temperature range being intermediate to that predicted for extant ratites and squamates of their size. This is not unexpected given their phylogenetic position relative to known ectothermic and endothermic crown members and is suggestive of an acquisition of elevated metabolic rates prior to the narrowing of core temperature ranges in defense of a stable core temperature. Similarly, we suggest that *Coelophysis* was more likely to maintain a ratite-like resting metabolic rate than a monotreme or squamate-like RMR. A core temperature range intermediate to extant ratites and squamates is also suggested, similar to *Plateosaurus*. The presence of variable depth, density, and distribution of epidermal insulation would not only allow for a broader range of environmental tolerances, it appears to be a physiological necessity for *Coelophysis* and likely for most small ornithodirans as they began to increase their resting metabolic rates above ancestral levels.

Our results illustrate the interconnected nature of morphology, physiology, environmental variables and how they constrain organismal energetics, behavior, and geographic distribution. Niche Mapper is a flexible tool that can be applied to extinct organisms in deep time whose body shapes have no direct modern analog.

## Acknowledgements

The authors would like to thank Aaron Kufner and Adam Fitch for discussion and productive feedback on earlier versions of the manuscript and Andrew Zaffos for help troubleshooting R code. Thanks also goes to the numerous colleagues who have acted as sounding boards and provided critical evaluations of this work as it progressed over the last few years; their efforts are greatly appreciated. A special thanks to the two anonymous reviewers whose suggestions vastly improved an earlier version of this manuscript.

## Supporting Information

### S1 Appendix Niche Mapper Variables

#### Niche Mapper Variables

The number of variables (691; representing all data for 1 day each month for 1 year) in the combined microclimate and physiological models seems large. However, the microclimate model can be broken into 13 discrete categories. For instance, the microclimate model may boast 198 variables, but the vast majority represent the range of a given category, such as minimum and maximum values for each modeled day (24 states). In this way 5 categories (relative humidity, air temperature, cloud cover, wind, and percent shade) account for 60% of microclimate variables. In other words, there are 24 variables for relative humidity (a high and low for each of the 12 modeled days of the year), etc.

The biophysical model can be broken into 12 categories for body shape (99 variables), 10 categories for heat generation and transient properties (179 variables), 4 categories for dietary/digestive/water properties (131 variables), and 4 categories for behavior (89 variables).

##### Microclimate model variables

| Model Parameters | Quantity | Comments: |
| --- | --- | --- |
| Number of days to model | 12 | A start, stop, and day number are chosen (from 1-365) |
| Substrate conditions | 8 | Roughness, conductivity, |
|  |  | density, heat, depth |
| Surface object profiles (bushes, rocks, etc.) | 4 | Experimental wind profiles<br>Set to zero if none measured. |
| Geographic Information | 10 | Hemisphere, latitude,<br>longitude, slope, azimuth,<br>elevation, |
| Maximum/Minimum % shade | 24 | Potential shade range for<br>model to choose from |
| Timing of maximum minimum | 8 | (e.g., air/wind max~solar<br>noon; relative<br>humidity/cloud~sunrise |
| Maximum/Minimum relative humidity | 24 | Aridity/humidity (min/max per<br>modeled day) |
| Maximum/Minimum cloud cover | 24 | Insolation (min/max per<br>modeled day) |
| Maximum/Minimum wind speed | 24 | (min/max per modeled day) |
| Maximum/Minimum air temperature | 24 | (min/max per modeled day) |
| Snow or free water presence/absence | 12 |  |
| Substrate reflectivity (albedo) | 12 |  |
| % of surface acting like free water | 12 |  |

##### Allometry (modeled organism dimension properties)

| Model Parameters | Quantity | Comments: |
| --- | --- | --- |
| Animal group/locomotion | 2 | Clade, and biped vs. quad |
| Head geometry | 9 | Length:width:height; fur depth<br>(dorsal and ventral); density<br>and head geometry |
| Neck geometry | 9 |  |
| Torso geometry | 9 |  |
| Front legs | 9 |  |
| Back legs | 9 |  |
| Tail/additional appendage geometry | 9 | (this is same as above but can be used to model a trunk as well) |
| Total length/scaling adjustments | 3 |  |
| Subcutaneous fat distribution/mass change | 7 | Distribution and region(s) of subcutaneous fat storage |
| Postures | 7 | Start/End postures for shade, sleep, and inactive (modeled body parts in contact with ground or as single lump) |
| Flesh conductivity (head-tail) (min) | 13 | Also accounts for torso overhang (shading legs) |
| Core temperature relation with appendages | 8 | average core temperature that is somewhere between ground and torso core temperature |

##### Physiological and behavioral properties

| Model Parameters | Quantity | Comments: |
| --- | --- | --- |
| General coding and output parameters | 10 |  |
| Heat storage; specific heat | 6 | Transient vs stead state; specific heat for flesh |
| Mass, subcutaneous fat, density, appendages | 11 | Whole body parameters, number of appendages; substrate interaction, fur/feather compression when lying down |
| Metabolic rate; activity multiplier; thermoregulation triggering | 4 | *how the model uses heat generated from activity |
| Forage rate; muscle efficiency; production heat | 4 | Contribution of activity to heat balance model |
| Insulation properties: head, neck, torso, tail dorsal/ventral | 33 | Diameter, length, depth, density, reflectivity, |
| Hair/feather length/depth dorsal/ventral | 48 | *allows for seasonal changes |
| Temperature and water loss from skin and metabolism | 15 | Core temp: min, max, target; skin wet/sweat, pilo/ptiloerect; skin/fur thermal conductivity; depth within fur/feather for radiant exchange |
| Core temperature (target) (hibernation) | 24 | Tc per modeled day |
| Time dependant change in mass/body fat | 24 | For each modeled day |
| Lungs and gut | 5 | O2 extraction min/max; gut passage time; fecal and urea, |
| Digestive efficiency | 12 | For each modeled day |
| Times basal for activity energy & food for it (1-7) | 24 | This can be used to simulate phenology of food available and reproduction timing. |
| Food: %protein,carbs, fat, dry matter | 48 |  |
| Diurnal, nocturnal, crepuscular, hibernate (fraction of day hibernating) | 60 | For each modeled day |
| Active/inactive on land or water | 24 | For each modeled day |
| Thermoregulation behavior | 13 | Use nest; burrow shade; climb, wind protection; night shade; dive; wade; thermoregulation order |
| Flight, burrow, nest properties, shelter | 34 | Specific to nest, flight,and burrowing behaviors including shape and thermal properties of structures and (burrow gas properties) |

### S2 Appendix Model test using Varanus komodoensis, Komodo Is

#### Model test using *Varanus komodoensis* on Komodo Island

Although Niche Mapper has been tested for reptiles ranging in size from small iguanids [1–3] to land iguanas in the Galapagos [4] and small and large birds [5–7] and mammals [8–10] we include here an additional test using the Komodo dragon, *Varanus komodoensis*, on Komodo Island. We obtained data for monthly climates for this location using Adagio in niche-mapper.com to point-and-click on Komodo Island to extract climate data from New and others [11]. Komodo morphology/proportions were obtained from web photos of wild komodos from Komodo Island and implemented in the alomvars.dat input file for Niche Mapper. Key physiological properties, e.g. maximum and preferred core temperatures, were obtained from Harlow and others [12]. We set the model to allow 24 hour activity, diurnal only, and diurnal + crepuscular to see if there were any noticeable differences in energy requirements.

We assumed a high-level squamate metabolism and a skin reflectivity of 15% based on black reptile skin (Galapagos marine iguana) reflectivities measured by WPP. Since the allometry subroutines in Niche Mapper will automatically scale the allometry isometrically, given the mass and a reference dimension, e.g. snout-vent length or shoulder height, we were able to run annual simulations for a 6.7 kg and 65 kg animal assuming a density of 1030 kg/m^3 for both animals, and compute the body core temperatures, activity patterns and food requirements for the month of November when observations recorded in the literature were made that could be used for comparison to the simulations [13, 14]. **Figure 1 in S2** Appendix illustrates the computed body temperatures that would occur with a squamate metabolism for the large, 65 kg, and small, 6.7 kg Komodo dragon. The measured shade air temperatures reported for this time are also plotted.

Computed activity hours, patterns of annual activity in sun/shade and energy and food requirements for the 65 kg dragon exhibit a consistent pattern of energy and food requirements while activity hours varied under three activity conditions: diurnal only, diurnal-crepuscular, and 24 hour potential activity (**Fig. 2 in S2 Appendix**). For comparison, a simulation of the 65kg dragon was conducted within the Triassic hot microclimate model to illustrate the impact on activity, shade use, and food requirements under our deep-time conditions (**Fig. 3 in S2 Appendix**). We also ran a simulation for an allometrically modified 6.7 kg dragon, since the young have apparent longer legs and narrower torso than adults to determine allometry effects (Fig. 2 in **S2 Appendix**).

**Figure 1.**
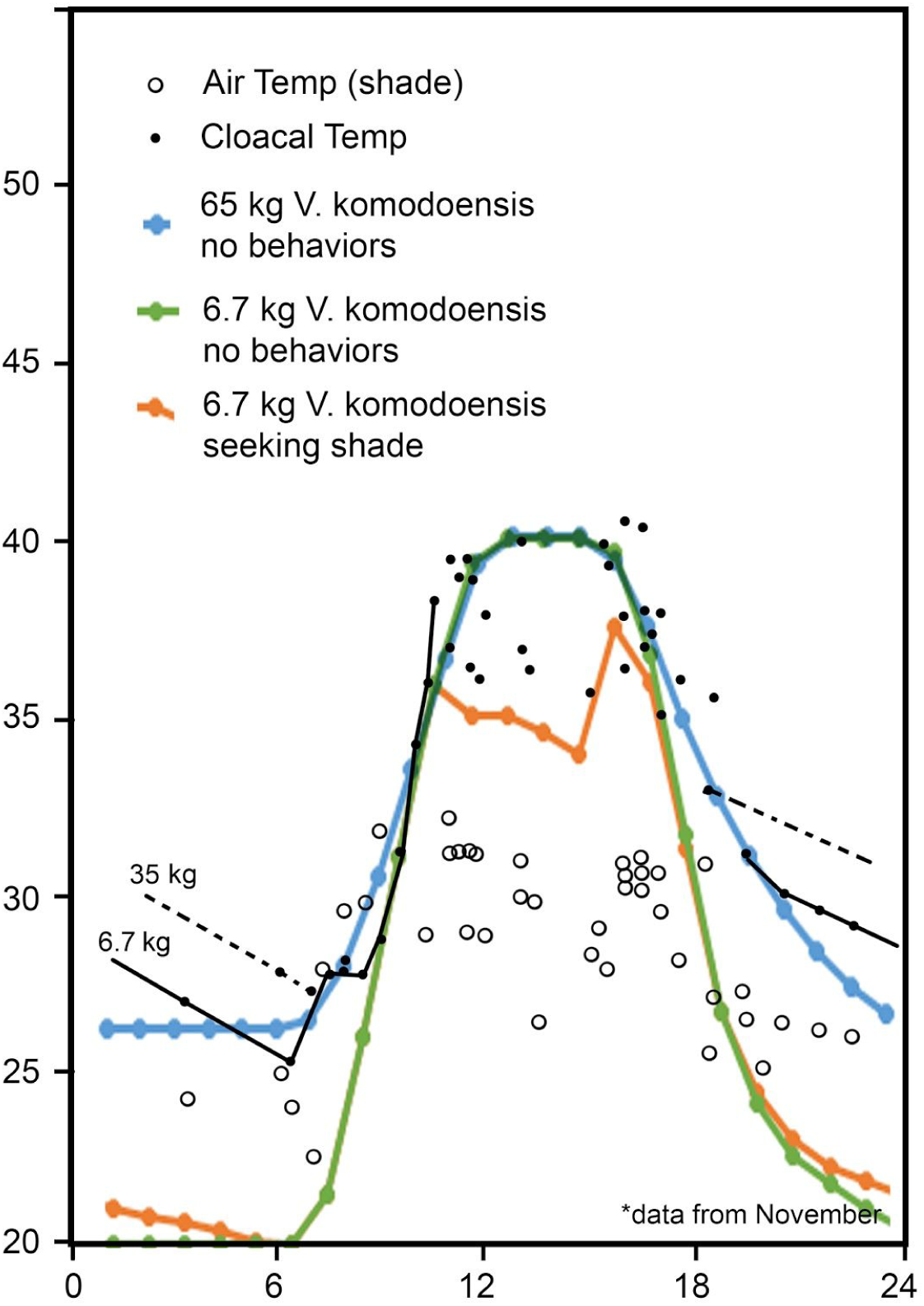
These data represent field cloacal temperatures measured from 28 individuals (closed circles) and heating and cooling curves from a 6.7 (solid black line) and 35 kg (dashed line) *V. komodoensi*s on Komodo Island including shaded air temperatures (open circles); modified after McNab and Auffenberg [13]. The 65kg *V. komodoensis* modeled in NicheMapper (blue line) demonstrates a similar hourly temperature pattern to the empirical data. The modeled 6.7kg *V. komodoensis* (green line) predicts a cooler night time temperature than available empirical data. Empirical data for the 6.7 kg animal is not recorded between the hours of 20:00 and 06:00 and and may be due to fossorial behaviour. The modeled 6.7kg *V. komodoensis* with thermoregulatory behaviors enabled (orange line) predicts a slightly warmer nighttime core temperature (seeking night shade) and cooler mid-day temperatures (seeking shade).

**Figure 2.**
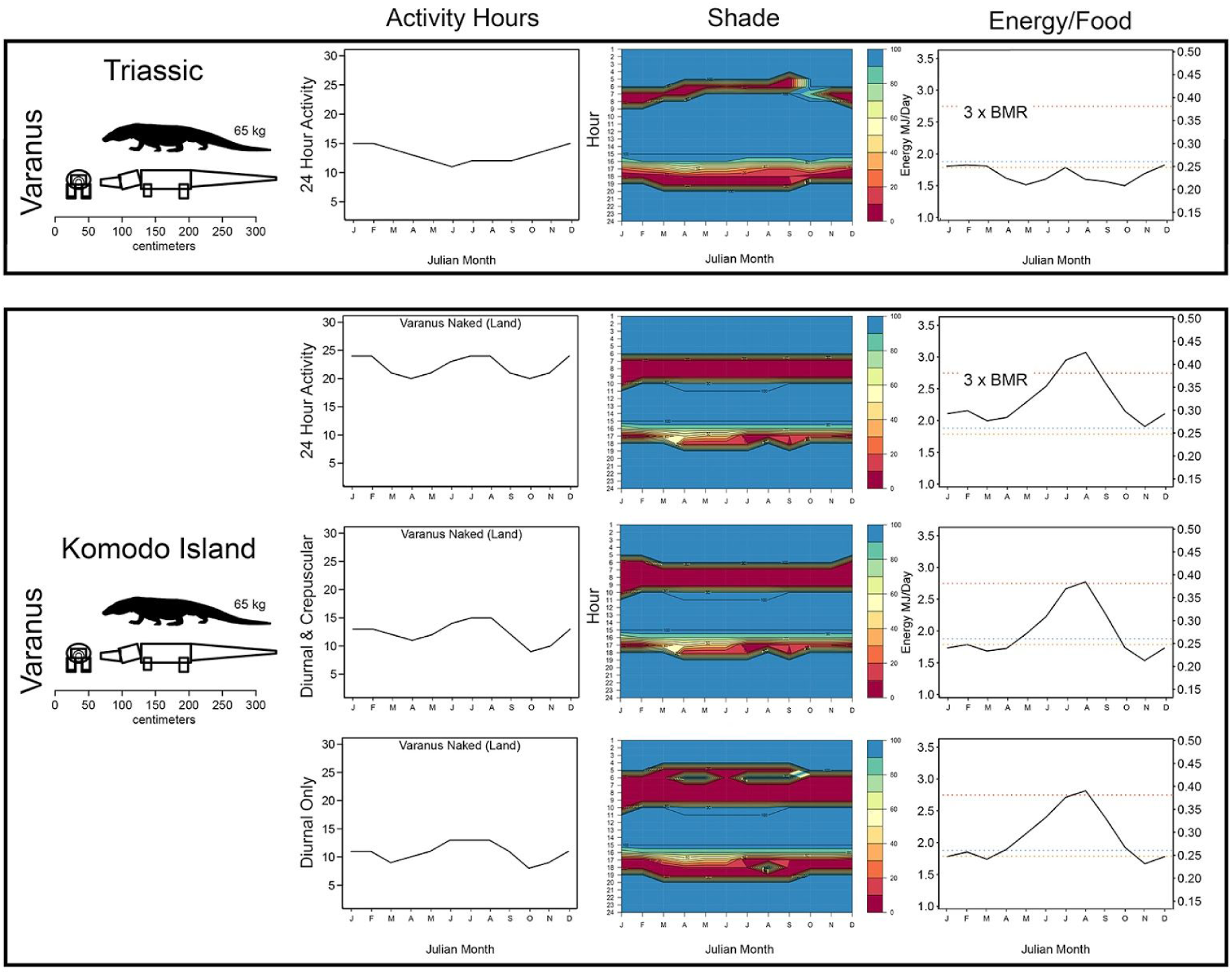
The 65 kg *V. komodoensis* was modeled in 4 different scenarios, 1 under hot Triassic conditions with 24 hours potential activity and 3 under Komodo Island conditions with a 24 hour, diurnal, and diurnal + crepuscular potential activity. In the ‘Energy/Food’ column, the increase in energy requirements mid-year on Komodo Island represents the cooler temperatures of the southern hemisphere winter. The modeled shade ranges from 0-100% (red-blue, respectively), with night shade representing the animal retreating from open air conditions to reduce net radiant heat loss to the night sky and cold ground surface, and day shade representing retreat to overhanging vegetation due to excessive core body temperatures. The modeled values for shade seeking behavior on Komodo Island reflect the observed bimodal activity peaks in the morning and evening [14]. *V. komodoensis* modeled in the Triassic hot monsoon microclimate exhibits a similar pattern to that seen on Komodo Island, though the activity hours over a 24 hour cycle and the energy requirements are shifted toward more nocturnality, likely due to higher T_air_.

**Figure 3.**
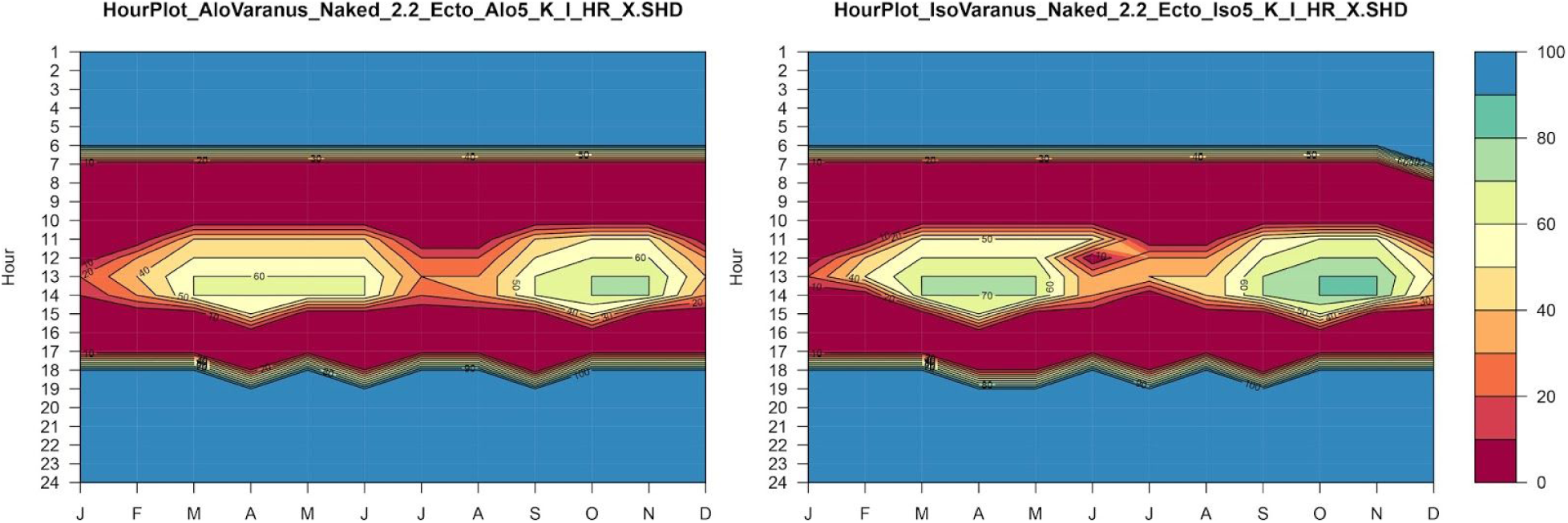
Allometric (left) vs isometric (right) scaling of juvenile Komodo dragon at 6.7 kg from an adult Komodo dragon of 65 kg. Young dragons have smaller bodies and longer legs relative to adult Komodo dragons. These graphs illustrate subtle (but minimal) changes in the amount of shade needed during the middle of the day in the hot times of the year on Komodo Island south of the equator.

**Figure 4.**
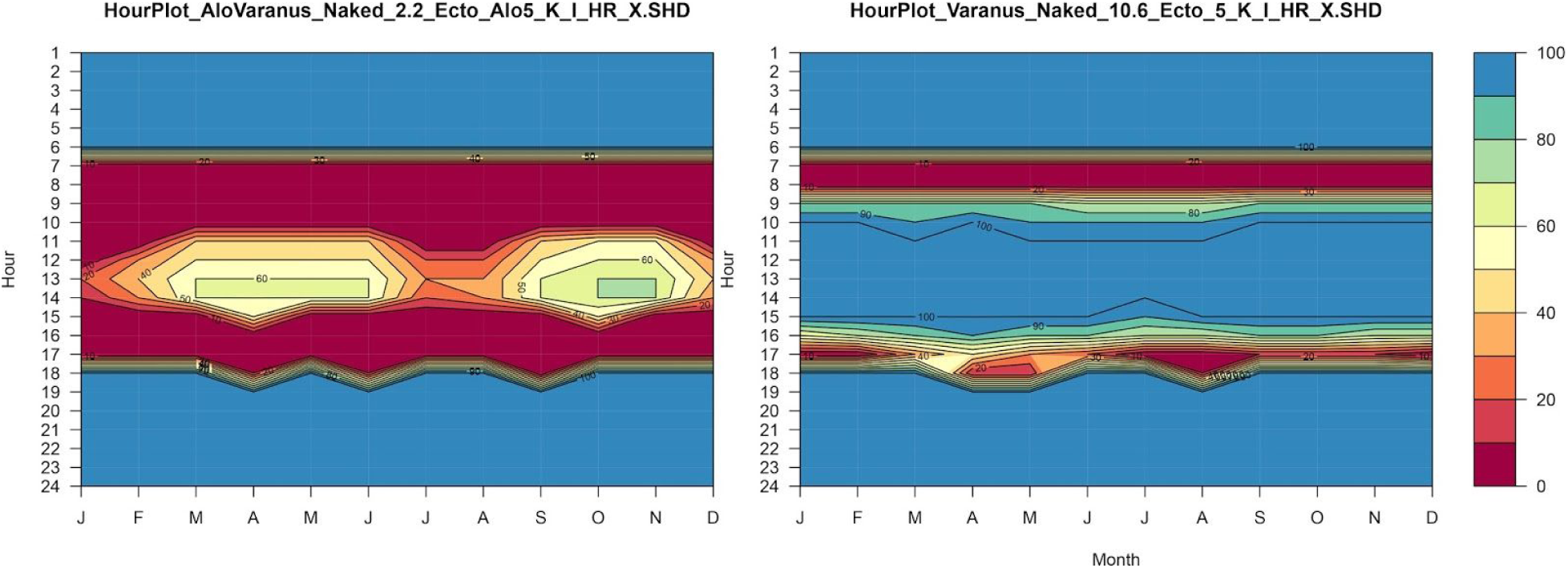
Computed 6.7 kg juvenile Komodo dragon activity pattern (left) vs. an adult 65 kg Komodo dragon activity pattern (right). These graphs illustrate the behavioral separation of activity patterns of small vs. large Komodo dragons during the day and throughout the year on Komodo Island south of the equator.

## S3 Appendix Parameterizing biophysical model dimensions for fossil vertebrates

### Parameterizing biophysical model dimensions for fossil vertebrates

Niche Mapper calculates its hourly heat balance equations utilizing a simplified geometric approximation of research taxa (Fig. 1), allowing for large numbers of simulations to run on relatively modest hardware. The results of this approach have been verified by comparison to high resolution models subject to large-scale fluid dynamics simulations in Ansys Fluent [1, 2]. In extant taxa such as *Varanus komodoensis*, parameterization of physical dimensions relies on direct or reported measurements. Extinct taxa require additional steps to ensure sufficient dimensional accuracy.

**Figure 1.**
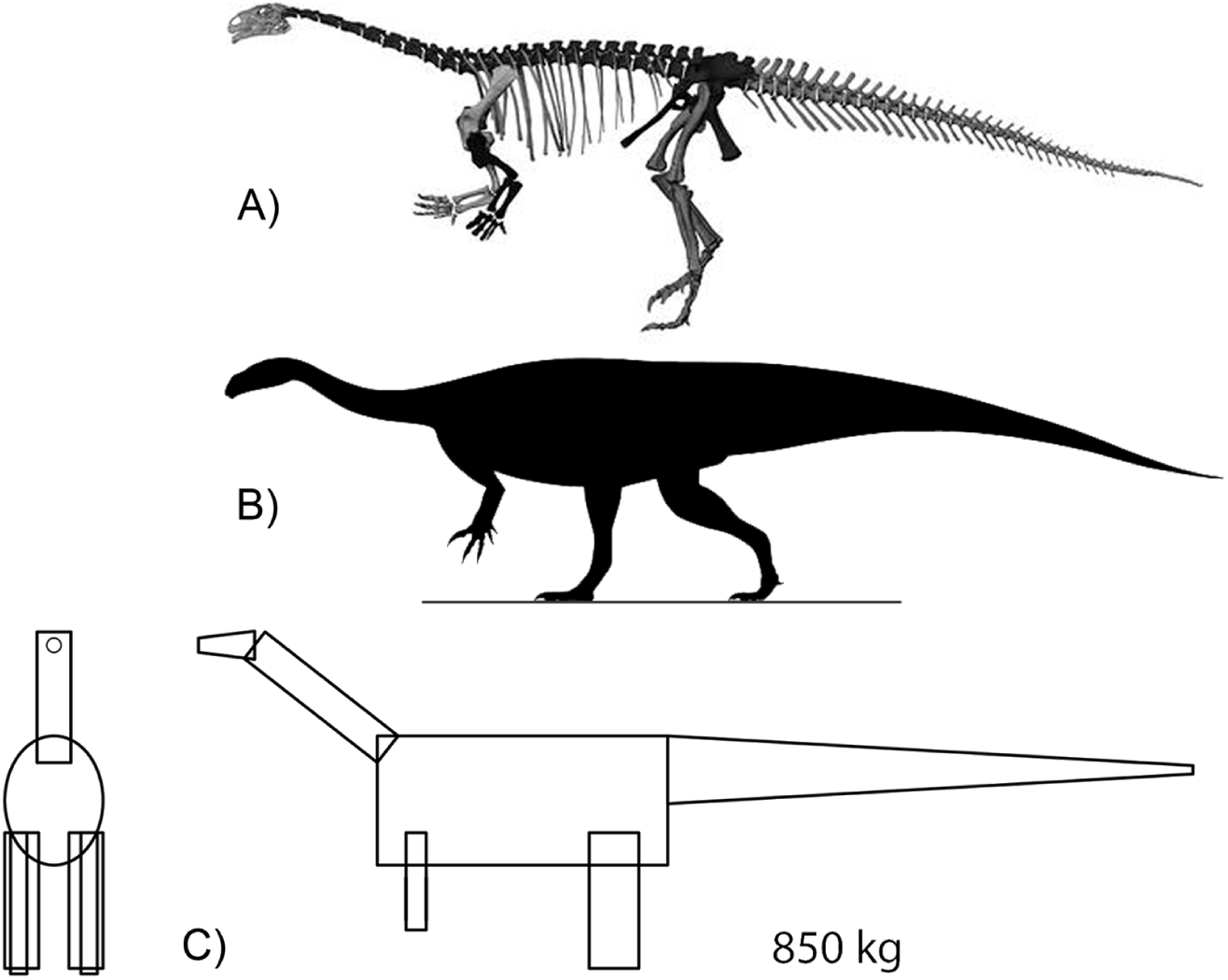
Modeled volume for *Plateosaurus*. Simplified overview of steps followed to create simplified geometric models for Niche Mapper biophysical input. Linear dimensions are taken from fossil data such as (A) this surface scan of GPIT125 [3]. That data is used for mass estimates, e.g. (B) lateral view silhouette used in GDI mass estimate, and the data is input to create a geometrically simplified (C) Niche Mapper model.

Dimensional inputs required by Niche Mapper include linear measurements and estimate of specific gravity for the head, neck, limbs, torso, and tail. An independent mass estimate is useful for comparison to the mass generated by Niche Mapper from the input dimensions and specific gravity of each body segment, as a check on potential errors in data input.

Dimensional parameters required by Niche Mapper are based on proportions of living animals. For *Coelophysis* and *Plateosaurus* estimates of life dimensions started with dimensionally accurate skeletal reconstructions (Fig. 2). Linear dimensions of individual skeletal elements were obtained from direct measurement of specimens and published data, as detailed in Wang, et al. [4]. The largest impact to dimensional proportions are competing interpretations of pectoral girdle placement [5].

**Figure 2.**
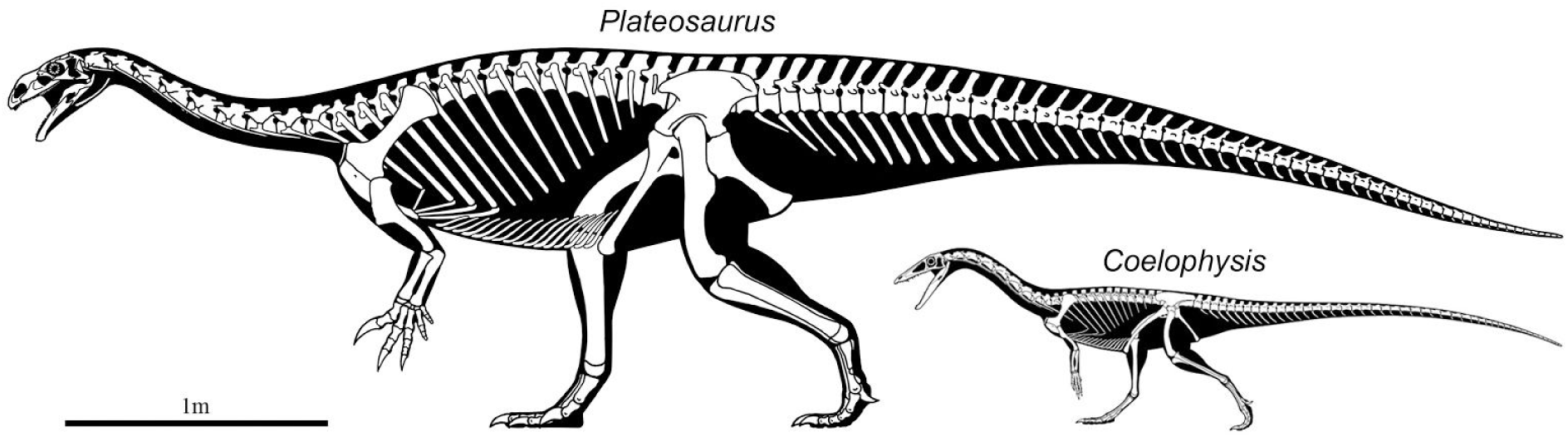
Skeletal reconstructions of *Plateosaurus* and *Coelophysis*.

Mass estimates for dinosaurs have a long history in paleobiology [e.g. 6], and have been attempted via disparate methods including limb bone allometry [7, 8], volumetric measurements of scale models [9,10,11], graphic double integration [12], computational application of minimum convex hulls [13] and various other dimensional analyses based on CT skeletal data [3,14,15,16]. The approach used in Niche Mapper is also a volumetric computational one, where total mass and mass distribution are calculated within Niche Mapper by assigning model volumes and densities for each body segment. Niche Mapper mass results were checked against Graphic Double Integration of rigorous skeletal reconstructions for *Coelophysis* and *Plateosaurus* [Fig. 2; 12,17] as well as previously published mass estimates [11, 15]. Diet composition was inferred from dental morphology. Skin transpiration and breathing efficiency were estimated from extant analogs.

## S4 Appendix Sensitivity analyses and additional figures

### Sensitivity Analyses

The strength our modeled results, in part, relies on understanding how sensitive the model is to ranges of values for variables that are not directly measurable in deep-time. As such, we endeavor to demonstrate that many of the variables (main effects and interactions) have relatively small effects on overall metabolic needs of the modeled organisms. However, we realize these effects can be cumulative and are more significant at the boundaries of a modeled organisms’ temperature tolerance where small changes can be the difference between survival or death. Variables that have a more significant impact, such as temperature, CTR, RMR, and insulation are presented with a range of inputs for each experiment, so that results can be compared and interpreted appropriately. The following sensitivity analyses were conducted to quantify the advantage or disadvantage our chosen values would impart on the model.

### Skin and insulation reflectivity

Reflectivity of the skin and epidermal insulatory structures for our organisms are modeled at 15% (0.15) which approximates a dark coloration. Reflectivity of black and white pelts of dairy cows are documented to be around 0.16 and 0.48, respectively (pers. observation, WP). To test the effect of color selection *Coelophysis* and *Plateosaurus* were modeled with various reflectivities (0.1, 0.15, 0.2, 0.25, and 0.6; see Fig. 5). There was little effect across this range with a fully insulated *Coelophysis* which is demonstrated to be near its target metabolic energy levels (∼1400 MJ/y) under the cold microclimate. Reducing insulatory covering to top-only and non-insulated individuals produced an energetic benefit of lower reflectivity (e.g., darker color) in the cold microclimate - the more cold stressed, the larger the effect. The benefit of darker color is still swamped by the overall cold-stress the modeled organisms experience in the cold microclimate (e.g., >5 x RMR). *Plateosarus* did not show a significant change in annual metabolic need with varying reflectivity. This suggests that thermoregulation behaviors are effective at moderating the effects of reflectivity regardless of color when an organism is near its target ME, and benefits of coloration only become realized well past the boundaries of tolerable temperatures.

**Figure 1.**
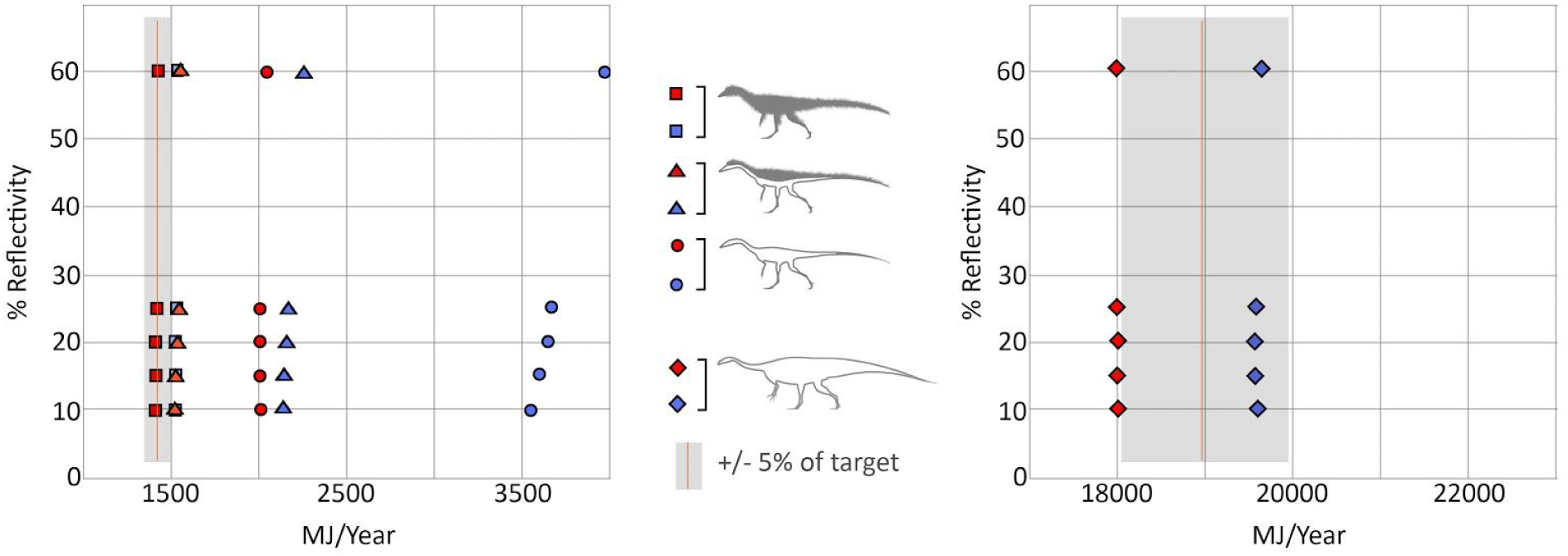
Skin/insulation color sensitivity analysis. The grey shaded area represents +/- 5% of the annual target ME. The further to the right of the target zone the greater the cold stress. Filled shapes: blue = cold microclimate, red = hot microclimate. The filled blue circles (uninsulated *Coelophysis*) exhibit the greatest advantage in decreased reflectivity due to extremely cold stress. When the modeled organism (e.g. fully insulated *Coelophysis*) is within its target, the effect of color is greatly minimized. *Plateosaurus* exhibits little to no benefit with changing reflectivity.

### Muscle efficiency

Most mammals, regardless of size, have a muscle efficiency of 0.25-0.30, and the vast majority of muscle efficiency for vertebrate clades ranges between 0.2 and 0.4 [1]; an efficiency factor of 0.2 in Niche Mapper (i.e., 20%) means that 80% of activity-generated energy is lost as heat instead of powering the activity. We chose to model our dinosaur’s muscle efficiency at 20% for all experimental runs. To test the impact of increased active muscle efficiency (e.g., decreasing muscle heat lost during activity) we ran experiments with 20, 30, 40, and 50% efficiency (Fig. 6). When the modeled organism is near its target metabolic energy the disparity from low to high efficiency is around 2.5% in the hot microclimate and 5-7% in the cold microclimate. The greater the cold stress the modeled organism experienced the greater the disparity between 20 and 50% muscle efficiency.

**Figure 2.**
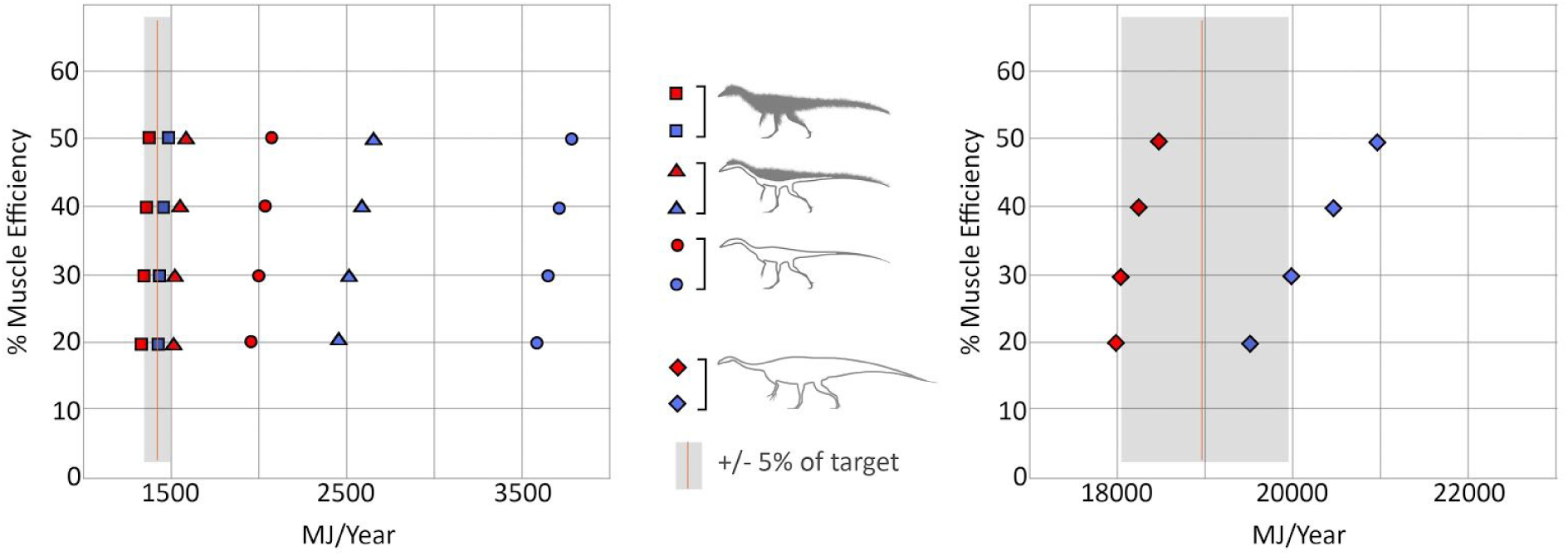
Muscle efficiency sensitivity analysis. The grey shaded area represents +/- 5% of the annual target ME. Filled shapes: blue = cold microclimate, red = hot microclimate. Fully insulated *Coelophysis* meets its target regardless of assigned muscle efficiency. As muscle efficiency is increased *Plateosaurus* is less heat stressed in the hot microclimate, but more cold stressed in the cold microclimate.

### Respiratory efficiency

Oxygen extraction efficiency (EO_2_) is used to calculate respiratory heat and water loss. When the organism is overheating one behavioral parameter in Niche Mapper allows the model to mimic panting by decreasing EO_2_ to ‘force’ the organism to breathe more rapidly leading to greater heat (and water) loss. Phylogenetic bracketing would suggest that early saurischian dinosaurs had a avian-like unidirectional airflow supported by numerous air sacs throughout the respiratory tract [2–4]. The range of EO_2_ values for avian lungs is much greater than other non-volant vertebrates (20-60%; [5]) We chose to use a 20% maximum and 15% minimum EO_2_, which is in line with values reported for ratites and many falconiformes (21-26% respectively; [5]). To test the effect of EO_2_ values we varied the max/min across four ranges 10/5, 20/15, 30/25, and 31/15; the last value range was to see if the max/min disparity had a noticeable effect. Varying the oxygen efficiency or max/min disparity had negligible effect on the annual energy budget (Fig. 7).

**Figure 3.**
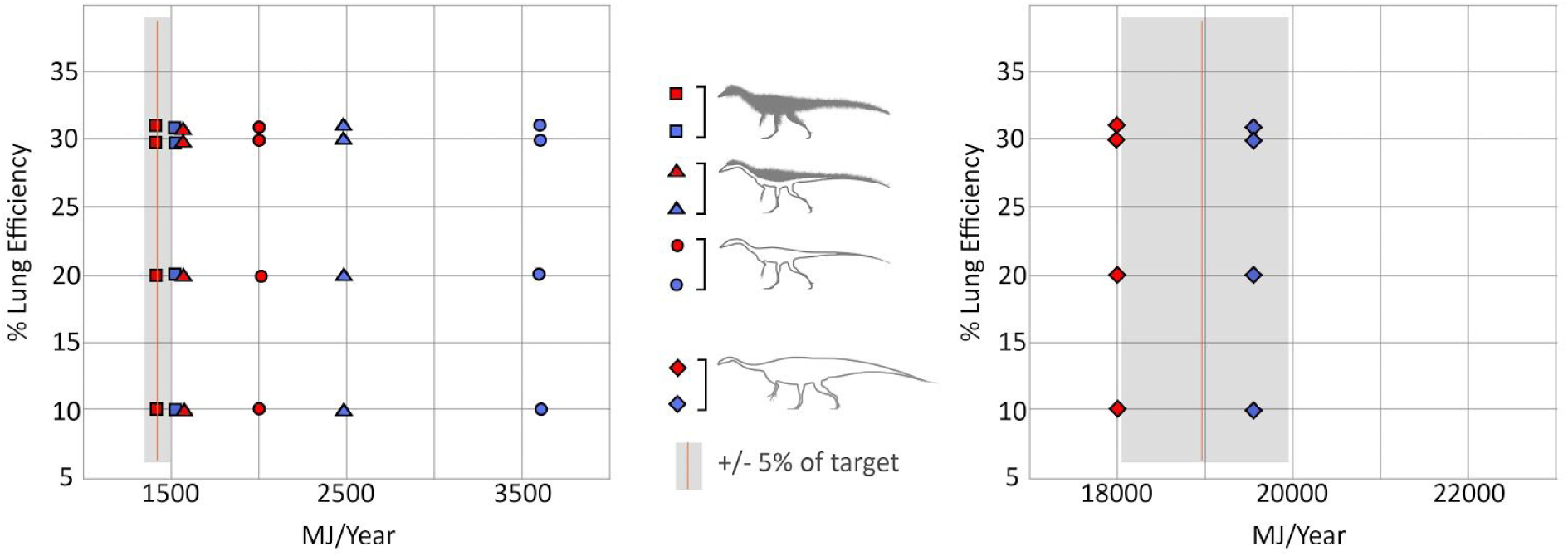
Respiratory efficiency sensitivity analysis. The grey shaded area represents +/- 5% of the annual target ME. Filled shapes: blue = cold microclimate, red = hot microclimate. Varying the lung efficiency had minimal effect on annual ME.

### Digestive efficiency

We modeled *Coelophysis* with a crocodile like digestive efficiency at 85% which is only slightly higher than the average digestive efficiencies of many birds of prey (75-82%; [6]). We varied digestive efficiency between 70 and 85% for our carnivorous taxon to determine the effect on total wet-food intake on an annual basis. *Plateosaurus*, modeled as an herbivore, was assigned a 60% digestive efficiency which is below the 70% efficiency seen in ratites [7], but above 50-55% seen in some herbivorous lizards as well as passerines on a low-quality diet [8, 9]. Others have argued for sauropod digestive efficiencies to be as low as 33% on a low-quality diet [10]. We tested a digestive efficiency range of 30-70% for our herbivorous diet.

As is the case with all dietary parameters in Niche Mapper varying digestive efficiency had no impact on the calculated metabolic energy (e.g., calculations for metabolic energy are independent of dietary calculations). The results (Fig. 8) are based on our assigned nutrient values (percent fat, carbohydrates, protein and dry mass; see Table 3) of the food source for the high browsing herbivorous and carnivorous diet. Changing the digestive efficiency of *Plateosarus* from 70 to 50% (a reasonable range with phylogenetic bracketing) resulted in a 70% increase total wet-food mass; the modeled *Plateosaurus* would need to ingest 3500 to 5000 kg (70-50% digestive efficiency respectively) on an annual basis. At the lower extreme, a 30% digestive efficiency would require ∼8000 kg wet-food per year; this is ∼22 kg of wet-food per day, which is on par with similarly sized extant browsing mammals such as the black rhinoceros [11, 12].

**Figure 4.**
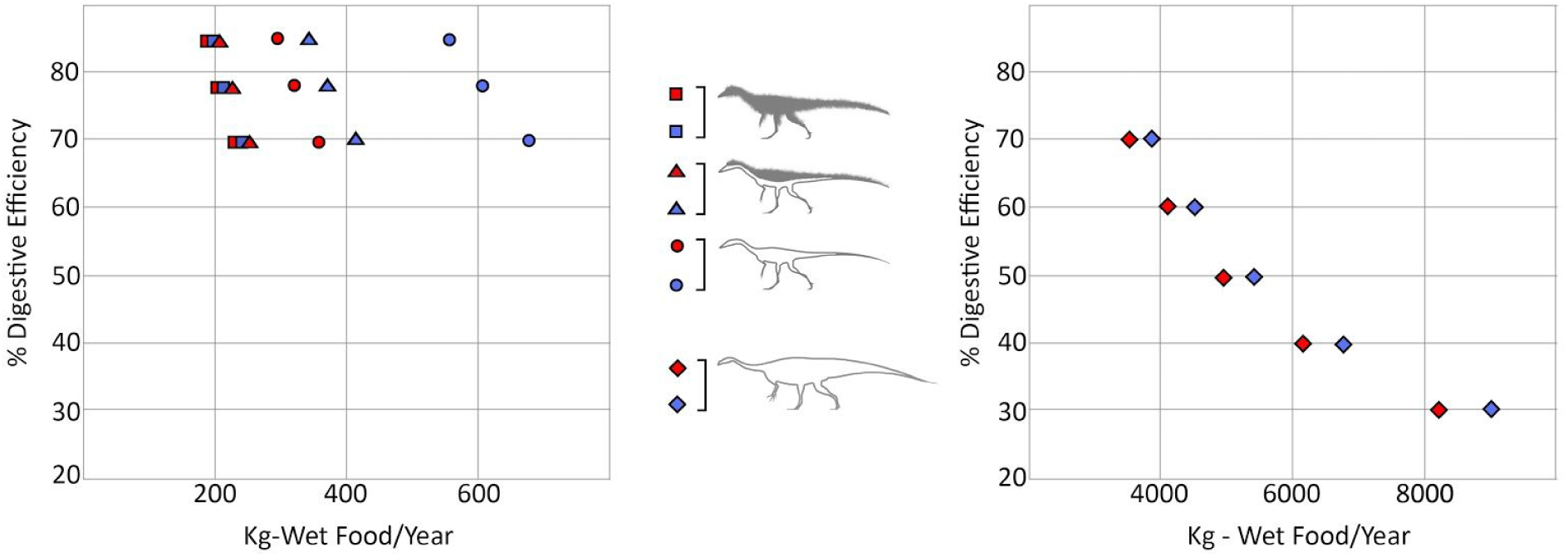
Digestive efficiency sensitivity analysis. Filled shapes: blue = cold microclimate, red = hot microclimate. Varying the digestive efficiency had no effect on annual ME, however it greatly impacted the annual quantity of wet food consumption.

### Effects of latitude

The cold microclimate we use for our experiments is intended to be a lower boundary representative of more temperate latitudes during the Late Triassic; these values are consistent with GCM models [13] for 45°N. However, to test the effect of insolation at higher latitudes we modeled our organisms at 45°N paleolatitude, in addition to the 12°N paleolatitude of the Chinle Formation we use to derive most of our microclimate model data. The primary effect of increasing latitude to 45°N appears to have been a result of increased daylight hours midyear and decreased daylight hours during the winter months. This is most apparent in the increased hours/day that core temperature was maintained, midyear, and decreased during the winter months relative to those observed at 12°N (Fig. 12). The model is more sensitive to microclimate temperatures than variance in insolation due to increased latitude. The remainder of the study uses insolation values from 12°N paleolatitude, and treats the cold microclimate as a surrogate for temperate latitudes. As more paleoclimate proxy data becomes available for higher latitudes this can be more rigorously tested.

**Figure 5.**
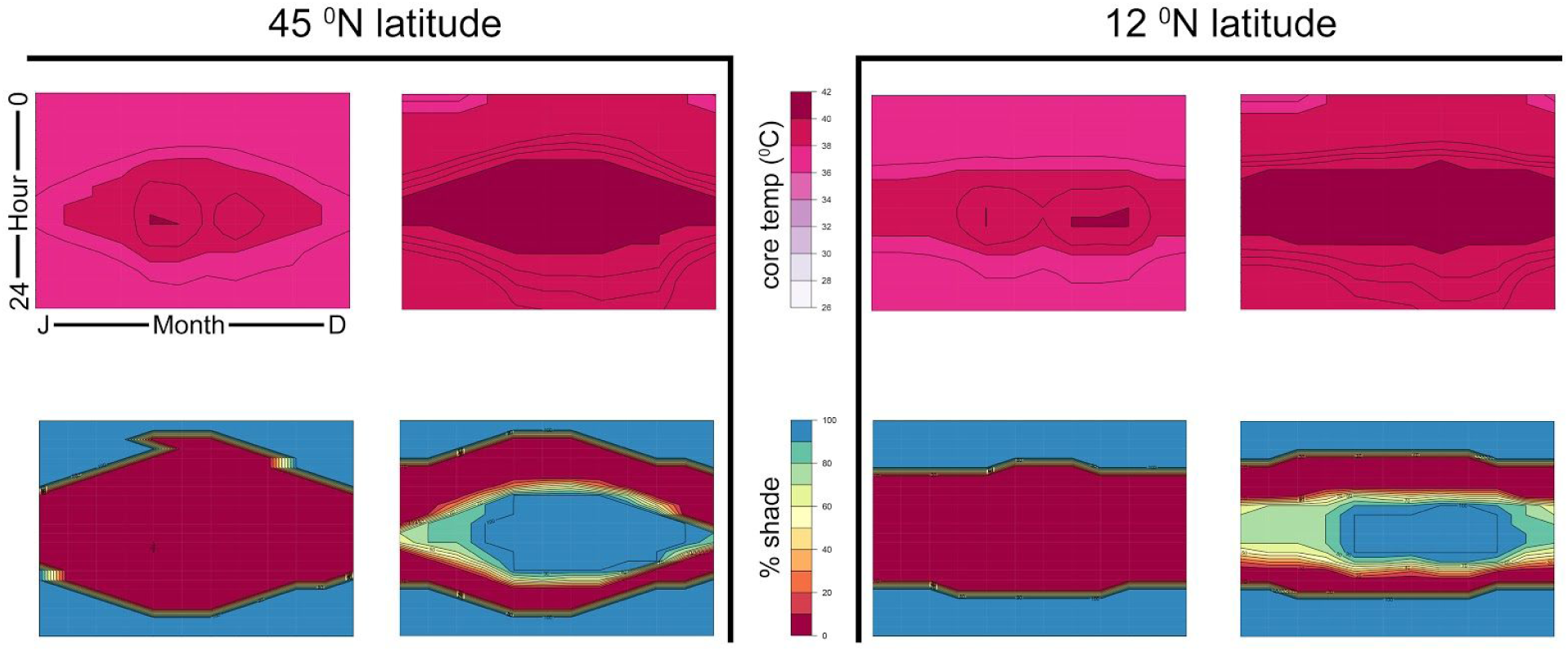
Effect of latitude (*Plateosaurus*) sensitivity analysis. Heat maps of core temperature (top) and shade utilization (bottom) demonstrating the increase in daylight hours midyear and decrease in daylight hours during the winter months at high (45°N) latitude relative to low (12°N) latitude.

### Yates Analyses of climate parameters

To test the main and interactive effects of four primary climate parameters (temperature, humidity, wind speed, and cloud cover) a 2^4^ factorial design and Yates’ algorithm [14] was analysed (Fig. 13). The results show that temperature has the largest effect on our modeled organisms’ annual energy budget. The fully insulated *Coelophysis* was modeled with and without the behavioral ability to ptiloerect. With ptiloerect enabled, *Coelophysis* was less affected by wind speed - likely due to an increased boundary layer provided by ‘fluffing’ up the insulation layer; temperature had an order of magnitude more effect than any other climate variable for the fully insulated *Coelophysis*. Wind speeds had the second greatest effect, while humidity and cloud cover were both negligible. The uninsulated *Coelophysis* and *Plateosaurus* had similar responses to the four climate parameters; temperature was still the dominant effect, but only by a factor of 2. Because temperature and insulation have a large effect on the results, we include all insulatory states, as well as temperature ranges for all of the following analyses.

**Figure 6.**
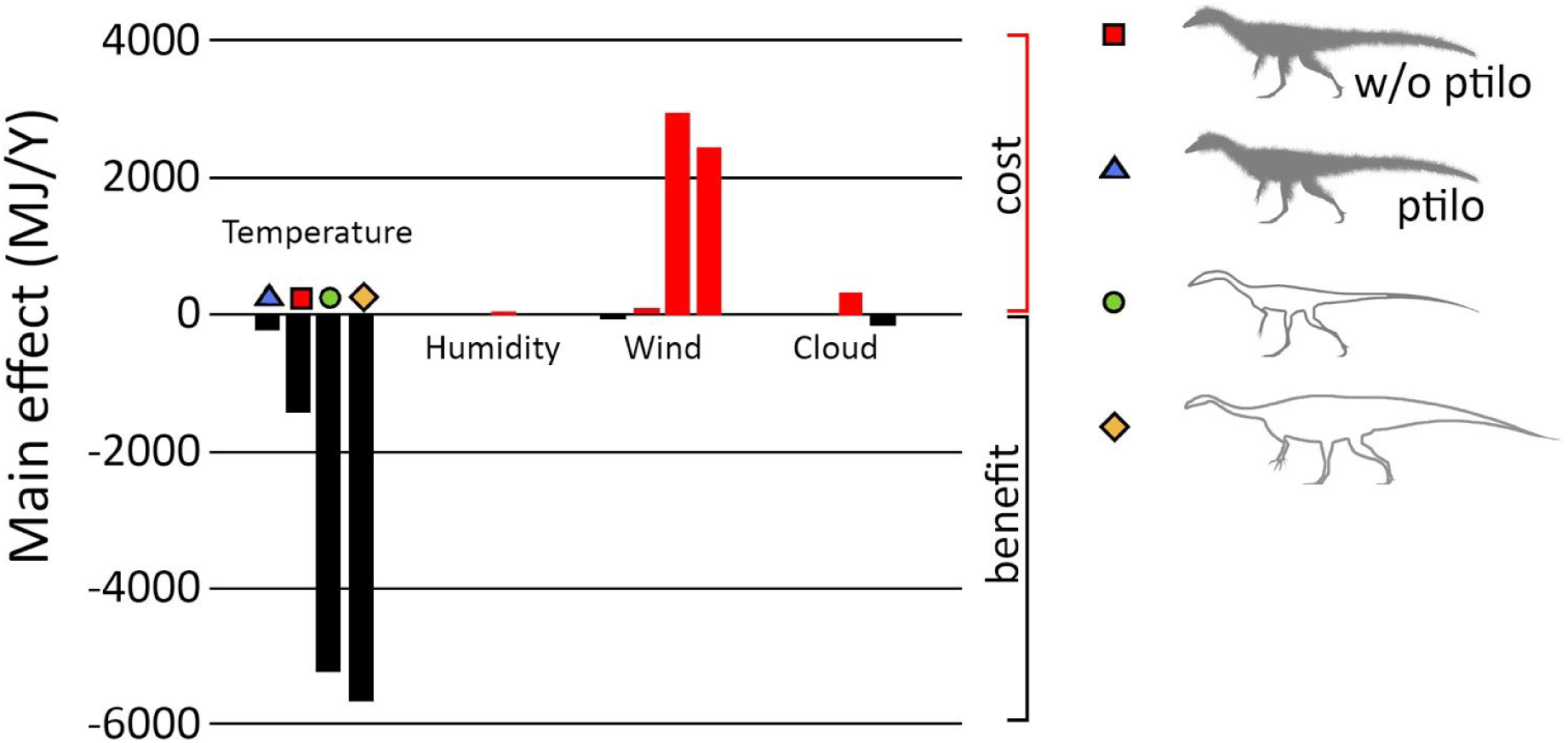
Yates analysis of climate parameters. Climate parameters temperature, humidity, wind, and cloud cover demonstrate the strong effect temperature has on annual metabolic energy. Wind was the second most significant effect, while humidity and cloud cover were both negligible. Note: both uninsulated models (*Coelophysis* and *Plateosaurus*) were strongly affected by temperature and wind; the insulated models were not as greatly impacted.

**Figure 7.**
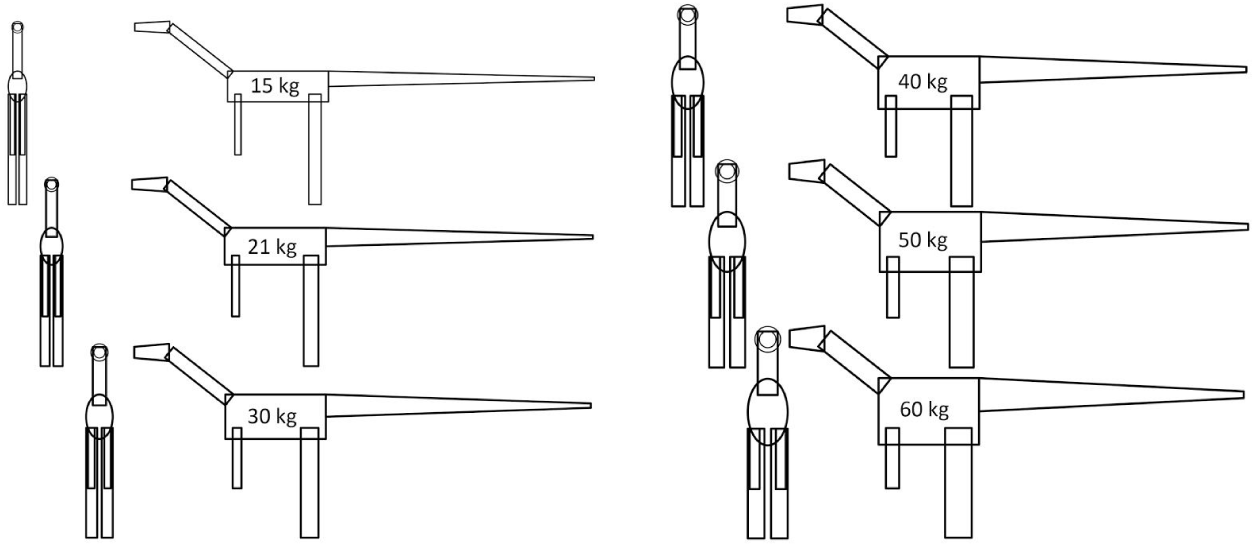
*Coelophysis* mass estimate sensitivity analysis. Six models were generated for a 15, 21, 30, 40, 50, and 60 kilogram *Coelophysis*.

**Figure 8.**
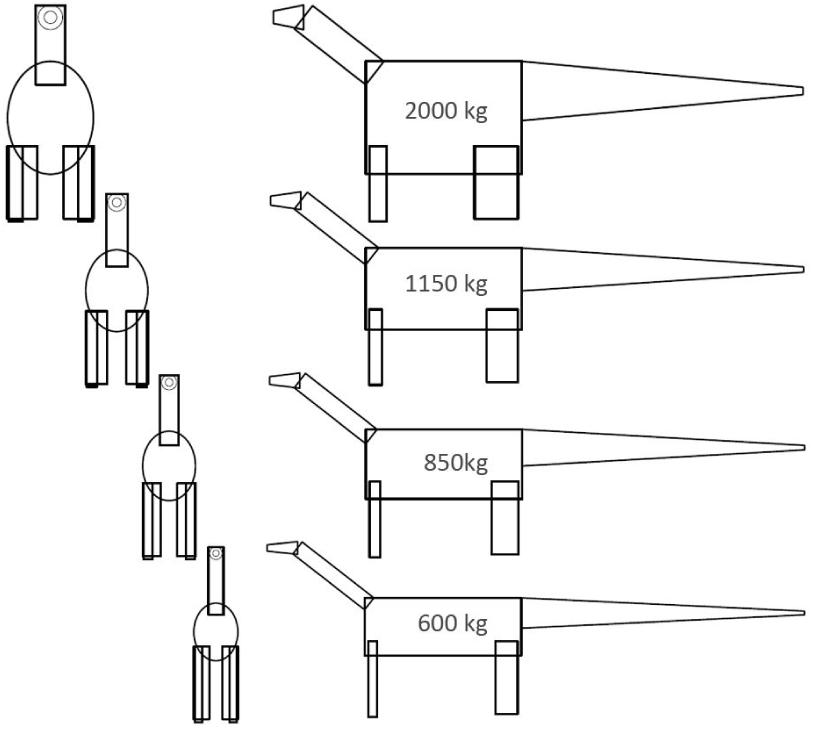
*Plateosaurus* mass estimate sensitivity analysis. Outlines of the Niche Mapper models represent the 600, 850, 1150, and 2000 kilogram *Plateosaurus*.

**Figure 9.**
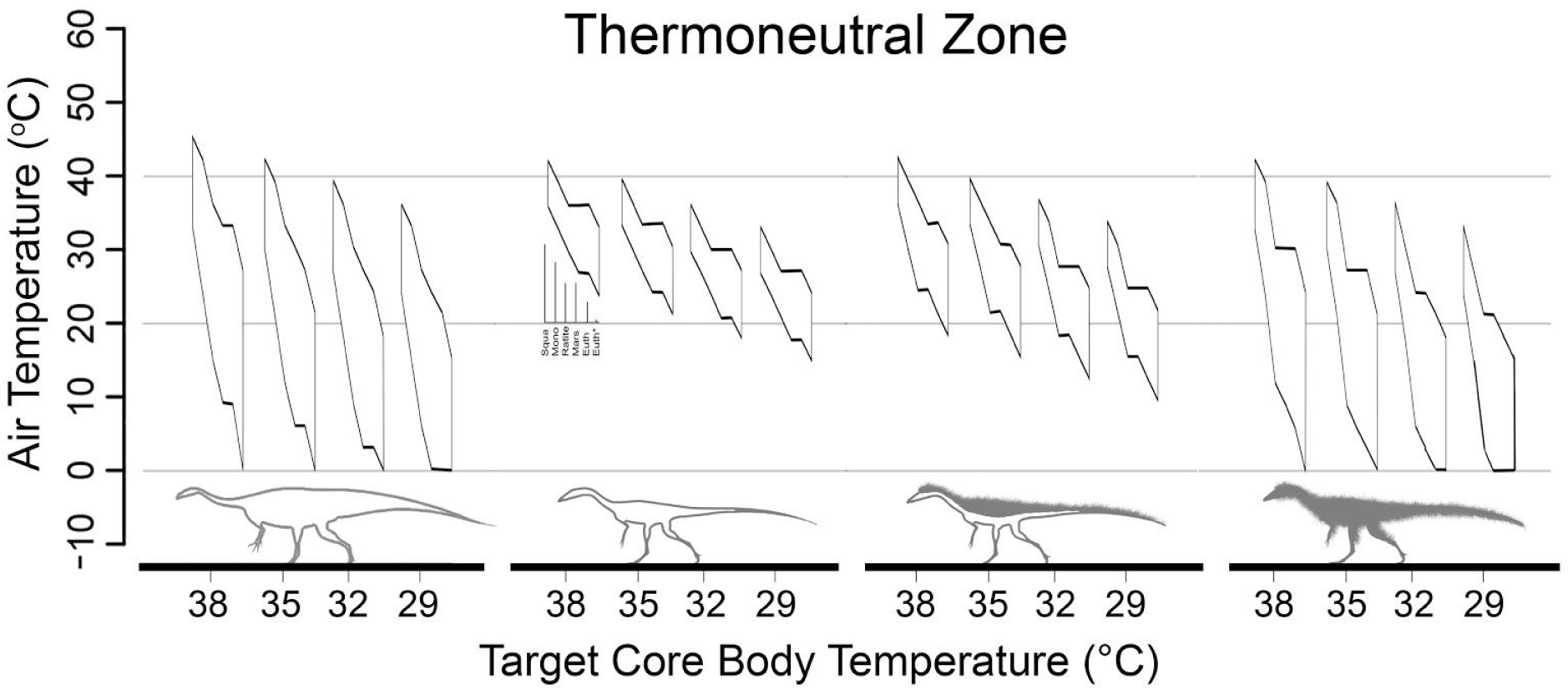
Effect of variable *target* core temperatures (narrow CTR). Active thermoneutral zones of 6 metabolic rates, left to right for each of the 16 polygons, (Squa = squamate [15], Mono = monotreme [16], Mars = marsupial [16], Ratite = ratite [17], Euth = eutherian [16], Euth*=eutherian [15] at four different target core body temperatures with a narrow core temperature range (+/- 2 °C). The thermoneutral zones for the lowest target core temperature for *Plateosaurus* and the lowest two for the fully insulated *Coelophysis* extend below zero °C, but our analyses stopped at 0°C. References [15,16,17] are in the main text.

**Figure 10.**
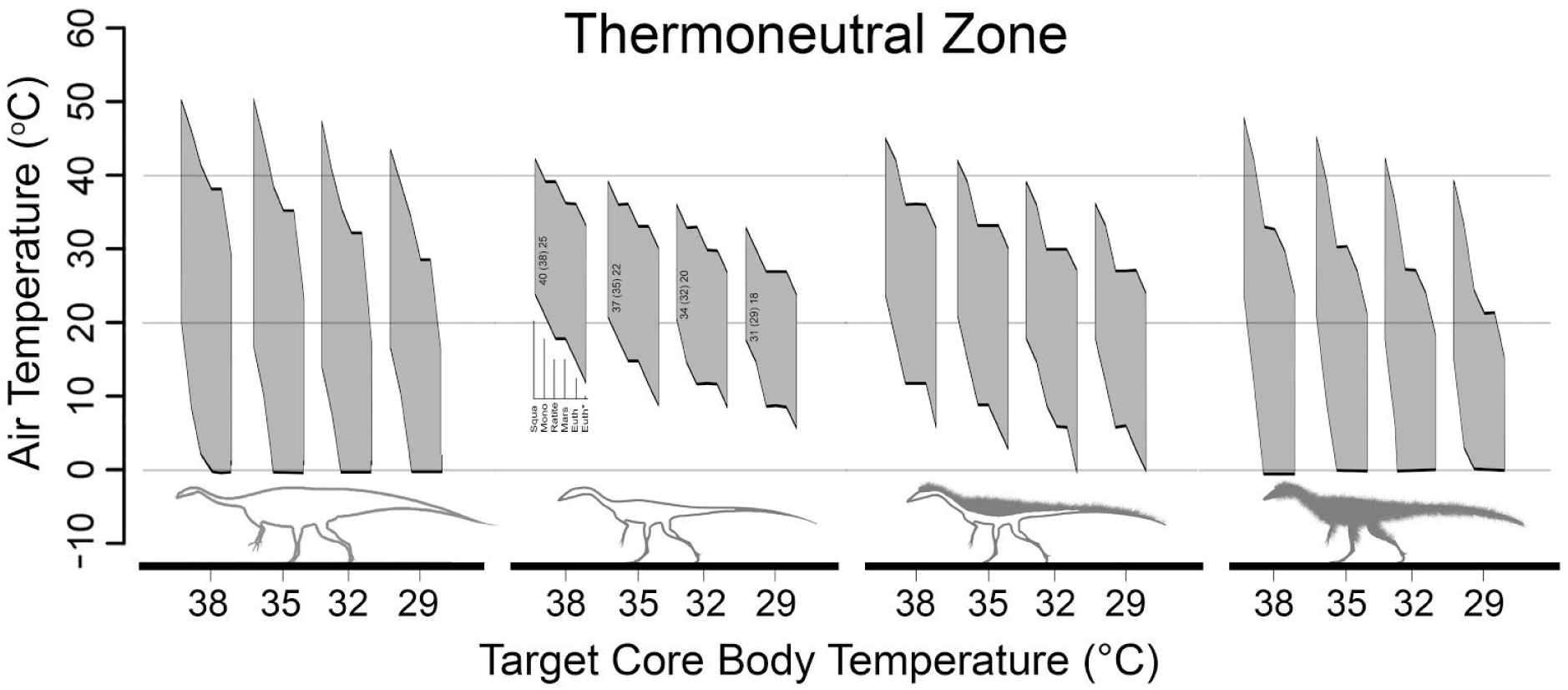
Effect of variable *target* core temperatures (broad CTR). Active thermoneutral zones of 6 metabolic rates, left to right for each of the 16 polygons, (Squa = squamate [15], Mono = monotreme [16], Mars = marsupial [16] Ratite = ratite [17], Euth = eutherian [16], Euth*=eutherian [15] at four different target core body temperatures with a broad core temperature range (+2/-13 degrees C). The thermoneutral zones for *Plateosaurus* and the fully insulated *Coelophysis* extend below zero °C, but our analyses stopped at 0°C. References [15,16,17] are in the main text.

**Figure 11.**
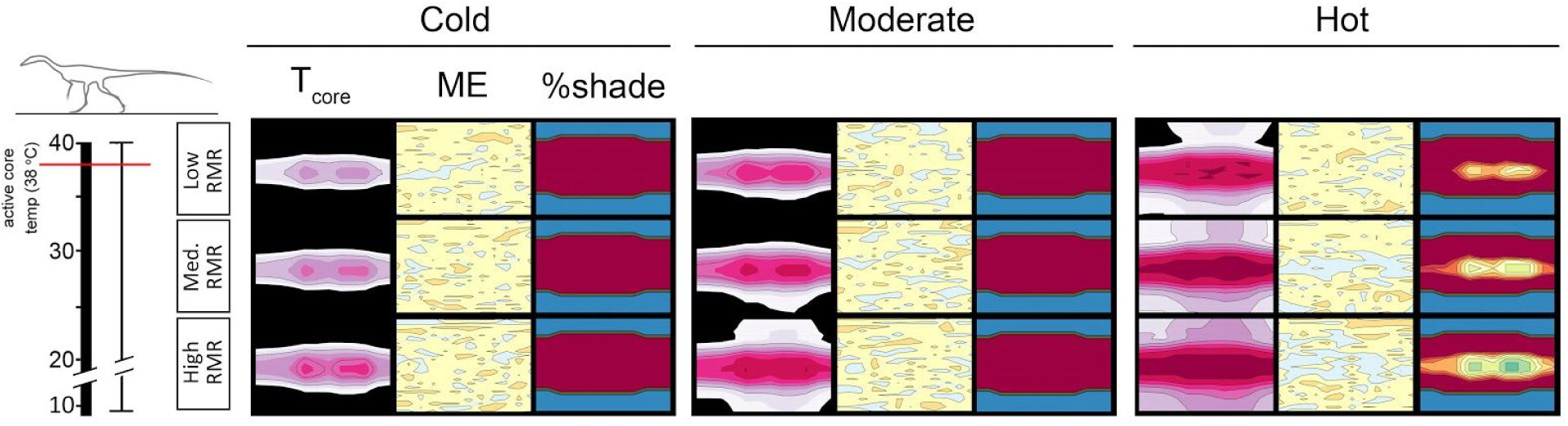
Heatmap of extended CTR for *Coelophysis* (uninsulated). Extending the lower boundary of the broad CTR allowed *Coelophysis* to maintain its target ME (+/- 5% of 2xRMR). However, T_core_ does not exceed 30°C for more than 5 months of the year in the cold microclimate. Black color in the T_core_= temperatures below 26°C.

**Figure 12.**
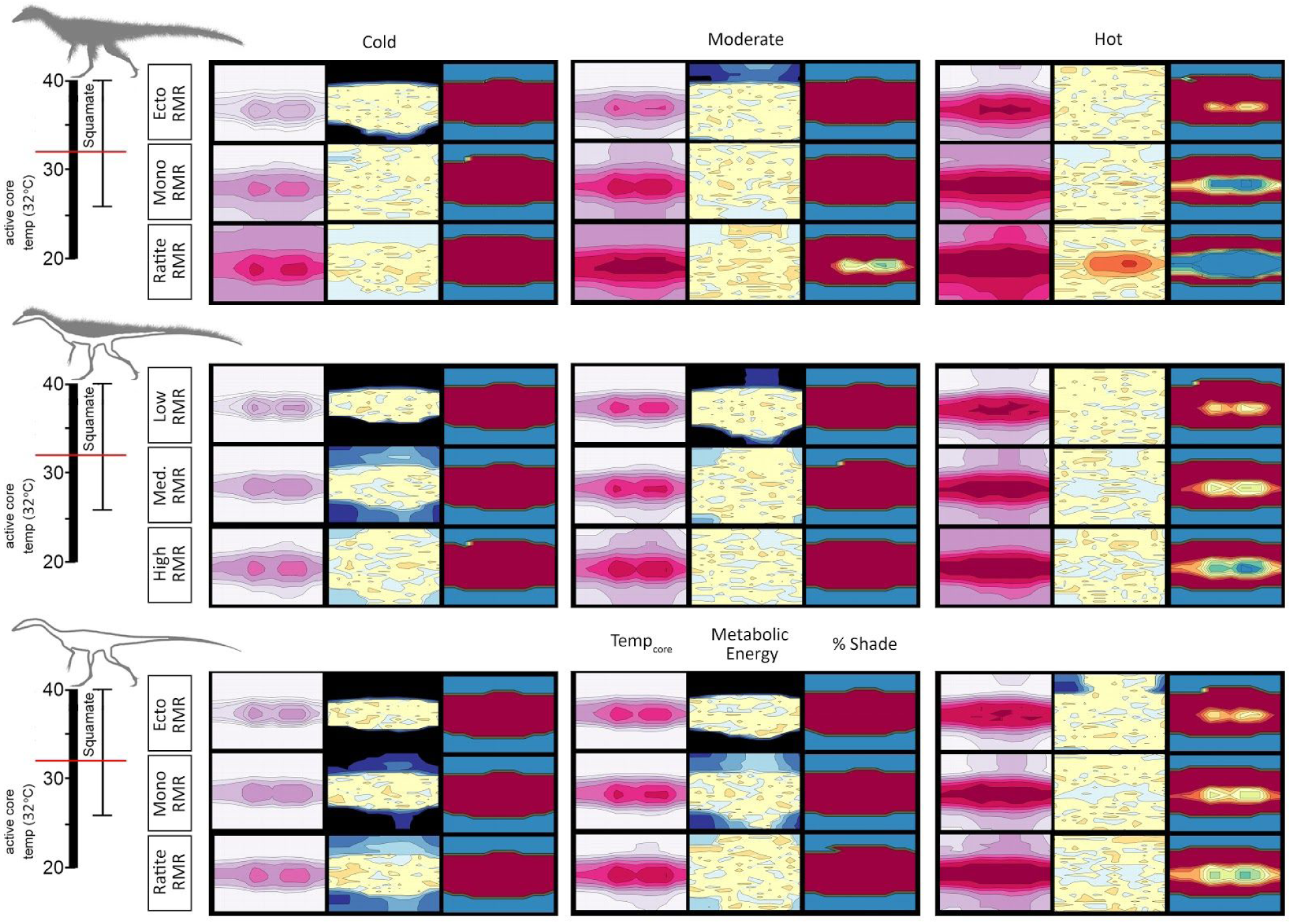
Heatmap of 32°C *target* T_core_ w/ broad CTR; *Coelophysis* (all). With a broad CTR *Coelophysis* was able to maintain a *target* T_core_ of 32°C with a high RMR, fully insulated in cold and moderate microclimates and with moderate to no insulation in the hot microclimates. Average T_core_ was: 33.2°C with high RMR and full insulation in the cold microclimate, 32.5°C with high RMR and top-only insulation in the moderate microclimate, and 34.7°C.

**Table 1.** Average annual T_core_ Coelophysis (all).

| Broad CTR |  | 32 degree Target |  |  |
| --- | --- | --- | --- | --- |
|  | RMR | Microclimate |  |  |
|  |  | Cold | Moderate | Hot |
| Furry | Squa | 27.6 | 29.3 | 32.9 |
|  | Mono | 30.3 | 33.2 | 35.8 |
|  | Ratite | 33.2 | 35.6 | 37.8 |
| FurryTop | Squa | 27.1 | 28.9 | 32.4 |
|  | Mono | 27.9 | 30.1 | 34.4 |
|  | Ratite | 28.9 | 32.5 | 35.6 |
| Naked | Squa | 27.3 | 29.0 | 32.0 |
|  | Mono | 27.9 | 29.7 | 33.4 |
|  | Ratite | 28.5 | 30.8 | 34.7 |

| Broad CTR |  | 38 degree Target |  |  |
| --- | --- | --- | --- | --- |
|  | RMR | Microclimate |  |  |
|  |  | Cold | Moderate | Hot |
| Furry | Squa | 27.5 | 29.5 | 33.2 |
|  | Mono | 31.0 | 34.3 | 37.5 |
|  | Ratite | 36.7 | 38.5 | 39.0 |
| FurryTop | Squa | 27.1 | 28.9 | 32.5 |
|  | Mono | 28.0 | 30.4 | 34.8 |
|  | Ratite | 29.6 | 33.2 | 37.0 |
| Naked | Squa | 27.4 | 29.1 | 32.0 |
|  | Mono | 28.0 | 29.9 | 33.5 |
|  | Ratite | 28.8 | 31.2 | 35.0 |

| Moderate CTR |  | 38 degree Target |  |  |
| --- | --- | --- | --- | --- |
|  | RMR | Microclimate |  |  |
|  |  | Cold | Moderate | Hot |
| Furry | Squa | 32.0 | 32.6 | 34.3 |
|  | Mono | 33.2 | 34.6 | 36.7 |
|  | Ratite | 36.1 | 38.1 | 38.8 |
| FurryTop | Squa | 32.0 | 32.3 | 33.8 |
|  | Mono | 32.0 | 32.8 | 34.5 |
|  | Ratite | 32.3 | 33.7 | 36.0 |
| Naked | Squa | 32.0 | 32.5 | 34.1 |
|  | Mono | 32.0 | 33.0 | 34.5 |
|  | Ratite | 32.3 | 33.6 | 34.8 |

| Narrow CTR |  | 38 degree Target |  |  |
| --- | --- | --- | --- | --- |
|  | RMR | Microclimate |  |  |
|  |  | Cold | Moderate | Hot |
| Furry | Squa | 36.0 | 36.2 | 37.2 |
|  | Mono | 36.4 | 37.0 | 38.0 |
|  | Ratite | 37.5 | 38.6 | 39.0 |
| FurryTop | Squa | 36.0 | 36.1 | 37.0 |
|  | Mono | 36.0 | 36.3 | 37.3 |
|  | Ratite | 36.1 | 36.7 | 37.7 |
| Naked | Squa | 36.0 | 36.1 | 37.1 |
|  | Mono | 36.0 | 36.3 | 37.3 |
|  | Ratite | 36.0 | 36.7 | 37.5 |

---

**S1 Table.**
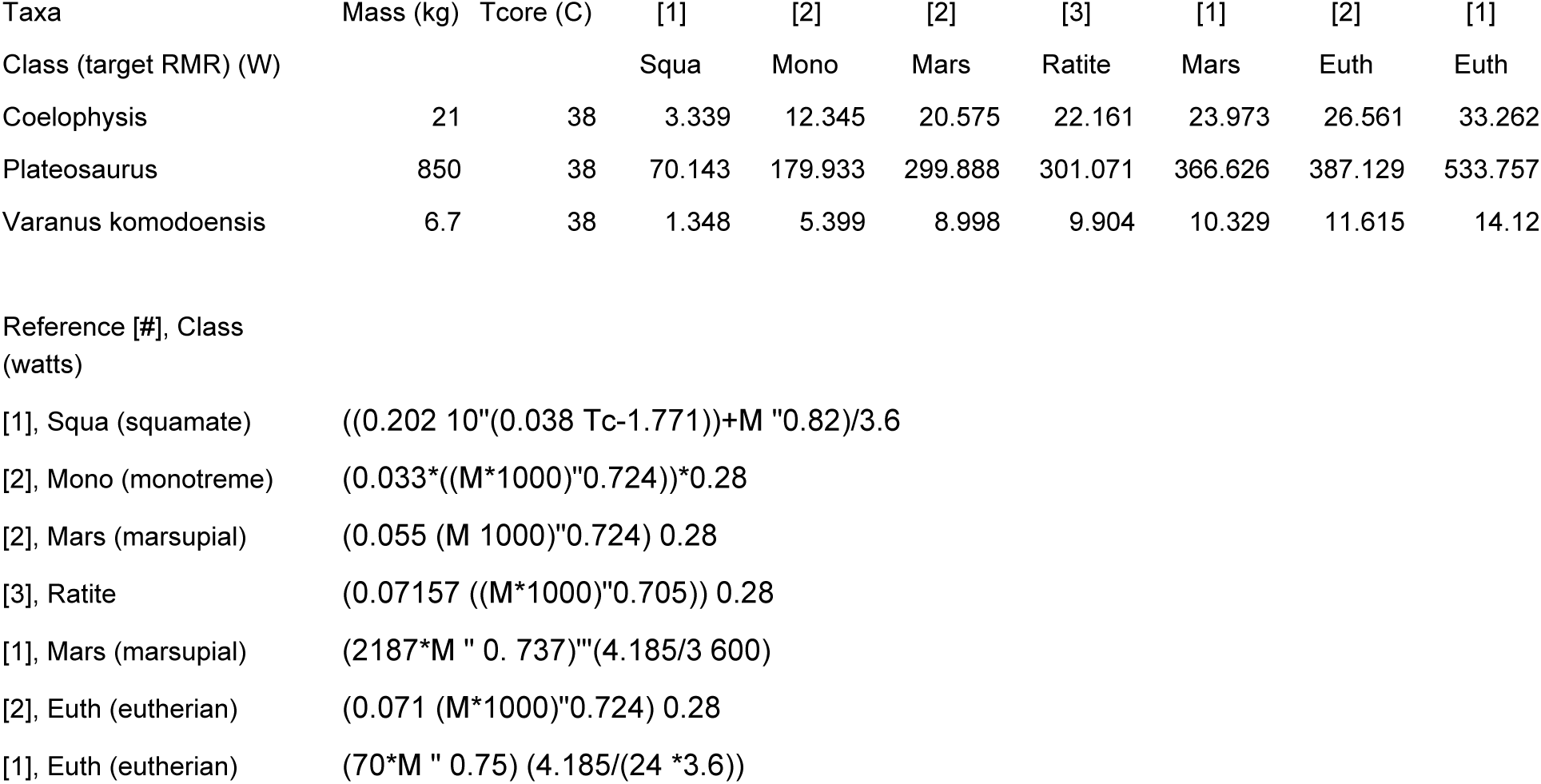
Target resting metabolic rates for classes of animals.

